# Multidimensional analysis of matched primary and recurrent glioblastoma identifies Fcγ receptors upregulation on microglia as a contributor of tumor recurrence

**DOI:** 10.1101/2023.04.21.537308

**Authors:** Tala Shekarian, Sabrina Hogan, Tomás A. Martins, Philip Schmassmann, Alexandra Gerber, Julien Roux, Deniz Kaymak, Célia Durano, Bettina Burger, Matthias Matter, Marie-Françoise Ritz, Gregor Hutter

## Abstract

*Background*: Glioblastoma (GBM) is a lethal brain tumor without effective treatment options. The aim of this study was to characterize longitudinal tumor immune microenvironment (iTME) changes in order to find potential actionable targets to prevent GBM-induced immune evasion mechanisms.
*Methods*: This study included 15 patient-matched treatment-naïve WHO grade 4 primary (pGBM) and recurrent (rGBM) tumors. RNA and proteins extracted from fresh frozen tumor samples from matched pGBM and rGBM were profiled via transcriptomics and proteomics, respectively. A tissue microarray containing paired formalin-fixed paraffin-embedded tumor samples was processed for spatial transcriptomics analysis.
*Results*: Differentially expressed genes and proteins between pGBM and rGBM were involved in pathways responsible for synapse development and myelination which have been shown to play a role in GBM recurrence. By categorizing patients into short and long time-to-relapse (STTR vs LTTR), we identified genes positively or negatively associated with TTR. Expression of Fcγ receptors and complement system genes such as *FCGR1A* (*CD64*), *FCGR3A* and *C3* in rGBM samples were negatively correlated with TTR, whereas expression of *DNMT1/3A*, and *SMARCA4*, involved in DNA methylation, were positively correlated with TTR. Spatial transcriptomic analysis of the tumor cell compartment showed enrichment of oligodendrocytes in rGBM, whereas the myeloid cell compartment switched from quiescent to activated microglia, was enriched in B and T cells, specifically in rGBM with STTR.
*Conclusions:* Our results uncover a role for CD64-expressing activated microglia in GBM recurrence and suggest that interfering with these cells may represent a therapeutic option for hindering GBM relapse.

**Key points:**

- Transcriptomic and proteomic differences exist between patient-paired primary and recurrent GBM tumors
- High expression of Fcy receptors genes on activated microglia at tumor recurrence is associated with shorter time to relapse.

**Importance of this study:** In glioblastoma (GBM), the tumor recurs in almost all cases after standard treatment such as surgery and chemo-radiotherapy. In this study, we longitudinally evaluated the immune- and neoplastic compartments using transcriptomic, proteomic, and spatial transcriptomics in patient-matched treatment-naive and recurrent tumor samples. By correlating gene expression with time-to-relapse, we identified a geneset associated with treatment resistance and faster tumor recurrence. Moreover, this study highlighted the plasticity of the myeloid compartment during disease progression and an unfavorable role of activated microglia in tumor recurrence.

## Introduction

Glioblastoma (GBM) is the most aggressive and common malignant brain tumor in adults. Given the widespread tumor cell infiltration into vital or eloquent structures in the brain, complete surgical excision is impossible. Therefore, despite multimodal standard of care (SoC) therapy consisting of maximal resection followed by concurrent chemo-radiation and maintenance adjuvant chemotherapy (temozolomide, TMZ), the tumor inevitably recurs, with an overall survival of only 15 months^1^. Thus, GBM is considered an incurable, treatment-resistant cancer, necessitating urgent novel therapeutic strategies. In recurrent GBM (rGBM), which emerges after SoC treatment, additional surgery is only possible in up to 25% of patients, depending on the location and infiltrative nature of the recurrence^2^. Subsequently, second-line therapies such as angiogenesis inhibitors or immune checkpoint inhibitors are administered to the patients in the frame of experimental clinical trials^3^. However, all clinical trials failed to display durable remission, prolonged survival and improved quality of life of the affected patients^4^.

Recurrent GBM has occurred as a model of resistance to therapy and is the most aggressive, invasive, and resistant adult brain tumor^5^. The heterogeneity of rGBM at its cellular, molecular, and genetic levels is more pronounced, making it one of the most complex studied tumors and thus hindering the identification of potential therapeutic targets^6^. Study about genetic differences between pGBM and rGBM, has concluded that intratumoral heterogeneity is causing the failure of multimodal therapies^7^.

GBM is characterized by a “cold” immune tumor microenvironment (iTME) containing high numbers of suppressive, pro-tumor immune cells such as regulatory T cells (Tregs), tumor-associated microglia and macrophages (TAMs), myeloid-derived suppressor cells (MDSCs) and exhausted T cells^8, 9^. TAMs are reprogrammed by GBM cells resulting in an ineffective anti-tumor response, and interactions between TAM and GBM cells promote tumor cell proliferation, migration/invasion, leading to a worse overall prognosis^10, 11^. Mechanistically, TAMs secrete growth factors, cytokines and chemokines that remodel the GBM iTME^12^.

Recently, one study investigated the differences between pGBM and rGBM tumor compartments via multi-omic approaches^13^. However, less is known about the changes in the iTME and the regulation of immunogenicity in rGBM. Therapy-resistant tumor clones give rise to rGBM encompassing a new, heterogeneous iTME with distinct features compared to pGBM^14^. Changes in the iTME during disease progression and concurring therapy have been reported, such as decreased GBM associated microglia/macrophages in rGBM samples compared with pGBM, with an increase of undefined CD45^+^ immune cells population in rGBM^15^. Further, radiotherapy increases the immune responsiveness of tumors by driving immune cell infiltration and enhancing immunogenicity^16^.This prompted the use of immunotherapy which showed very promising results in a large variety of solid tumors, but, unfortunately, did not shown favorable results in GBM so far^17, 18^. Thus, alternative therapies combining targeted, rGBM-specific iTME features promoting tumor malignancy should be considered.

Molecular markers identified in the initially resected tumor tissue may only marginally inform us about its progression, resistance to therapy and recurrence potential. Therefore, we investigated differences in gene and protein expression in patient-matched pGBM and rGBM samples, and correlated the most significant changes observed in rGBM with time-to-relapse (TTR). Further, we deciphered TTR-specific changes in cell type composition of tumor and myeloid cells by spatial transcriptomics. Our results expand the understanding of the underlying biological processes on the path to rGBM, and highlight potentially targetable pathways for rGBM treatment.

## Material and Methods

### Ethics approval and consent to participate

All patients provided written, informed consent according to the legislation of the local ethical committee (EKNZ 02019-02358), for material and data collection supporting these analyses. The study was conducted following the ethical principles of the Declaration of Helsinki, regulatory requirements, and the code of Good Clinical Practice.

### Demographic and clinical information of the patient cohort

We retrospectively included 15 patients with pGBM and rGBM (WHO grade IV, *IDH* wild type) operated at the Neurosurgery Clinic, University Hospital of Basel from 2010 to 2020. GBM diagnosis was confirmed postoperatively by board-certified neuropathologists at the Institute of Pathology and Genetics, University Hospital Basel, Switzerland. Twelve patients were MGMT promoter methylation negative, whereas 5 patients had complete or partial MGMT promoter methylation (Table 1). All subjects received perioperative corticosteroids followed by conventional chemo- and radiotherapy before the second surgery. Some patients received adjuvant therapy such as Avastin, Lomustin, Fortemustin, and Optune after the second surgery. Time to relapse (TTR) was determined by calculating the interval between the dates of first and second surgeries (range 148 to 921 days). Patients’ information, including age, gender, treatments and clinical data (OS and TTR) are summarized in Table 1 and **Table 1S**.

**Table 1.** Compiled patients’ information including diagnostic and therapeutic data, dichotomized according to times to relapse.

| **Characteristic** | **Relapse time (cut-off=309d)** |  |
| --- | --- | --- |
|  | **long, n = 9<sup>1</sup>** | **short, n = 9<sup>1</sup>** |
| **Age at diagnosis** | 57 (8) | 57 (9) |
| **Sex** |  |  |
| Female | 2 (22%) | 2 (22%) |
| Male | 7 (78%) | 7 (78%) |
| **Diagnosis** |  |  |
| GBM IV | 9 (100%) | 9 (100%) |
| **IDH status** |  |  |
| Wild type | 9 (100%) | 9 (100%) |
| **MGMT-promotor methylation status** |  |  |
| negative | 6 (75%) | 6 (67%) |
| positive | 2 (25%) | 3 (33%) |
| NA | 1 | 0 |
| **Adjuvant Treatment** |  |  |
| CRT | 9 (100%) | 9 (100%) |
| **time to relapse (d)** | **471 (208)** | **210 (32)** |
| **Survival since 1st surgery (d)** | **801 (317)** | **457 (167)** |
| **Subtype** |  |  |
| Classical | 1 (11%) | 3 (33%) |
| Mesenchymal | 1 (11%) | 1 (11%) |
| Proneural | 7 (78%) | 5 (56%) |
| **Transcriptomic analysis** | **8** | **8** |
| **Proteomics analysis** | **2** | **4** |
| **GeoMX Spatial transcriptomic analysis** | **7** | **3** |

<sup>1</sup>Mean (SD); n (%); n; CRT chemo-radiotherapy; NA: not available

### Total RNA extraction and gene expression assay

GBM tissue samples were collected directly in the operating theater during primary (first) and recurrent (second) surgeries and immediately stored in RNAlater stabilization solution (Thermo Fisher Scientific, USA) and stored at −80 °C. For RNA extraction, tumors were mechanically dissociated in lysis buffer and total RNA was extracted using the AllPrep DNA/RNA/protein mini Kit (Qiagen, USA, Cat. No. / ID: 80004) according to the manufacturer’s protocol. RNA purity and concentration were assessed using the 2100 Bioanalyzer (Agilent Technologies, USA). Gene expression was assessed using the nCounter PanCancer IO 360 and the Neuroinflammation panels comprising 770 genes each (NanoString Technologies, Seattle, WA, USA), using 50 ng of total RNA per sample, according to the manufacturer’s protocol.

### Transcriptomic data analysis

Data collection was performed using the nSolver Analysis system and nCounter Advanced Analysis 2.0 software (Nanostring Technologies, Seattle, WA). Samples passing the normalization steps were considered for further statistical evaluations. Final dataset contains 30 samples (primary and recurrent) from 15 patients and 1320 unique genes. Comparison of expression levels between pGBM and rGBM was performed using limma from the Bioconductor package, patient effect was included in the model. The same model was used to assess the differences between samples from patients with short and long TTR. The threshold was set at 309 days in order to have similar sized groups. The Benjamini-Hochberg (BH) correction was used to obtain adjusted p-values. Hierarchical clusterings of gene expression were adjusted for patient effect using the *removeBatchEffect* function from the *limma* package. The gene set enrichment analysis and over representation analysis of GO terms were performed using the *clusterProfiler* package and the adjusted p-values were calculated using the BH procedure.

### Proteomic and data analysis

Paired fresh-frozen tissue samples were available from a subset of 6 patients. After microscopic quality control of cryosections stained with hematoxylin to eliminate tissue with excessive blood or necrosis, proteins were extracted using the standard protocol of the Protein Core Facility, Biocenter, University of Basel, and processed for proteomic analysis. Details and experimental parameters are outlined in **Supplementary Methods 1**.

Comparison between pGBM and rGBM expression levels was performed using limma from the Bioconductor package, patient effect was included in the model. The BH correction was used to obtain adjusted p-values. Hierarchical clusterings of gene expression were adjusted for patient effect using the *removeBatchEffect* function from the *limma* package. The gene set enrichment analysis and over representation analysis of GO terms were performed using the *clusterProfiler* package and the adjusted p-values were calculated using the BH procedure.

### Spatial transcriptomics

A tissue microarray (TMA) of a subset of 7 patient-paired primary and recurrent FFPE GBM samples was constructed (overview shown in **Fig. S3**) and subjected to spatial transcriptomic analysis using the NanoString’s digital spatial profiling technology (GeoMx DSP, NanoString, Seattle, WA, USA) as described in **Supplementary Methods 2**.

### GeoMx RNA data processing

GeoMx DSP data were evaluated and prepared for downstream analysis following the GeoMx-NGS gene expression analysis workflow until the filtering step. Segments with less than 4% of the genes detected were removed and the cut-off for gene detection was set at 2%. Reads from duplicated samples were summed.

The package limma was used to perform between-samples cyclic loess normalization and to test for differential expression using a linear model (lmFit).

Principal component analysis (PCA) was performed on the filtered gene set using log2 cpm (count per million) values and was plotted using the fviz_pca function from the factoextra Rpackage. SpatialDecon (v1.4.3) was used to determine the cell type composition.

### Data and material availability

All data needed to evaluate the conclusions in the paper are present in the paper and/or the Supplementary Materials. Transcriptomic and spatial transcriptomic data have been deposited to xx under the reference numbers xx and xx respectively. Proteomic data has been deposited to xx under the reference xx.

## Results

### Genes associated with synapse formation are enriched in rGBM

To explore the transcriptional immune signatures of patient-matched pGBM and rGBM specimens, we performed a targeted analysis of 1320 genes involved in iTME and neuroinflammation using specific transcriptomic panels in 15 GBM, with an age range of 39-65 years at the time of diagnosis (Fig. 1A).

**Figure 1.**
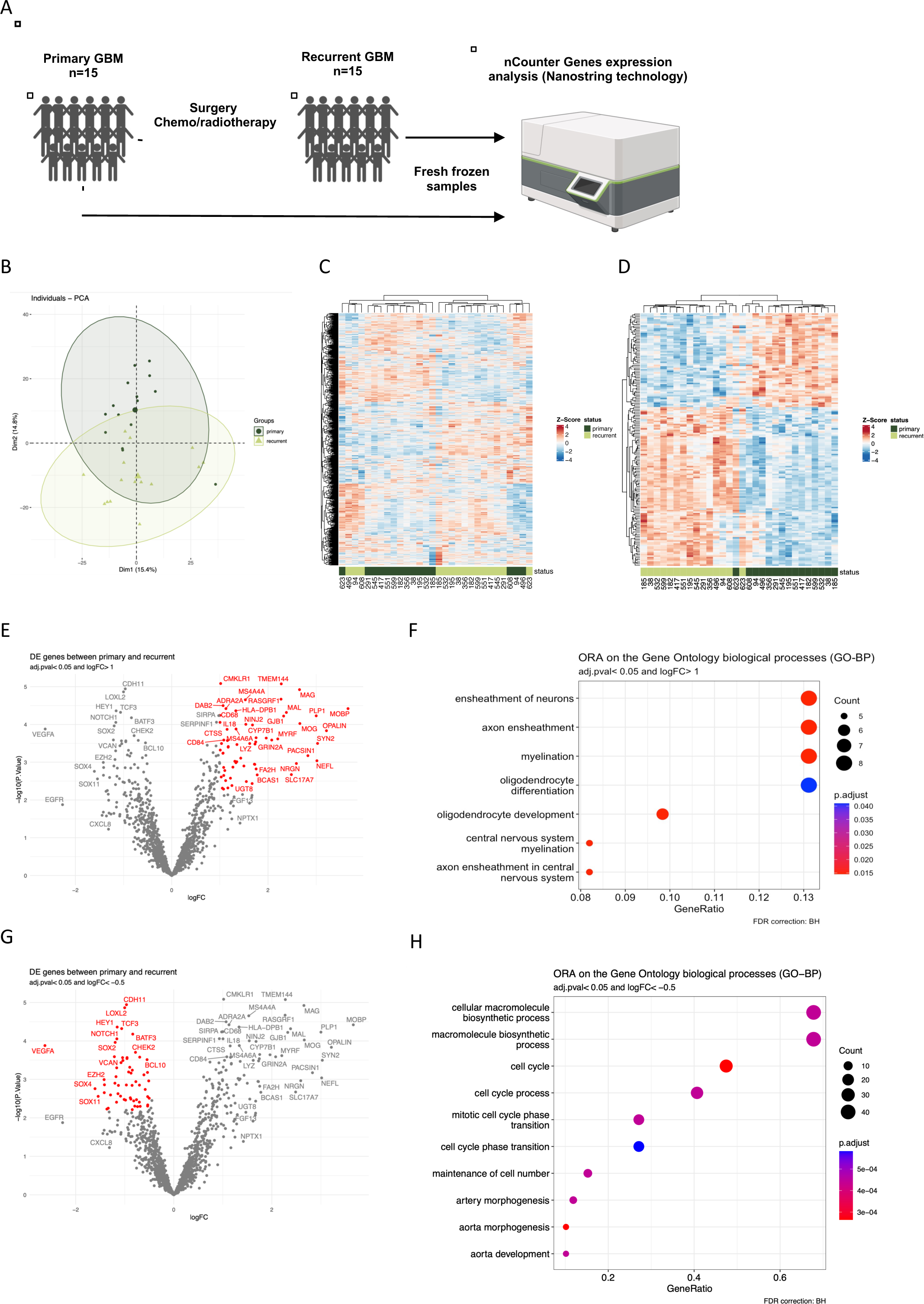
Targeted transcriptomic analysis of patient-matched pGBM and rGBM reveals synapse formation and oligodendrocyte differentiation as pathways activated in rGBM. **A)** Experimental setup for gene expression analysis using RNA extracted from fresh-frozen samples from 15 patient-matched pGBM and rGBM tumors using Nanostring nCounter system. **B)** PCA plot of all samples included in the analysis, each dot represents a patient, color coded according to pGBM or rGBM. **C)** Hierarchical clustering heatmap of the genes expressed across all samples. Z-score transformation was performed for each protein. Genes with similar expression patterns were clustered together. Relative expressions were scaled from red (high expression) to blue (low expression). Each column represents a sample (dark green: pGBM, light green: rGBM), and each row represents a gene. **D)** Hierarchical clustering heatmap of the genes differentially expressed across all samples. **E**) Volcano plot showing significantly differentially expressed genes in rGBM (red) (adjusted p<0.05 and log_2_FC>1). **F**) dot-plot of the corresponding enriched biological processes (GO-BP) determined by ORA. **G)** Volcano plot showing significantly differentially expressed genes in pGBM (red) (adjusted p<0.05 and log_2_FC>1) and **H**) dot-plot of the corresponding enriched GO-BP determined by ORA.

Principal component analysis (PCA) encompassing all investigated genes displayed partial segregation of pGBM and rGBM (Fig. 1B). Unsupervised hierarchical clustering reinforced the PCA results by displaying a consistent separative signature between pGBM from rGBM (Fig. 1C). Up to 167 differentially expressed genes (DEGs) between pGBM and rGBM were identified (Fig. 1D, and **Table S2**). Significantly upregulated DEGs in rGBM encoded for myelin associated proteins/glycoproteins such as *MAG*, *MAL*, *MOBP*, *MOG* and *PLP1*, while downregulated DEGs in rGBM encoding for proteins involved in angiogenesis such as *VEGFA* and in extracellular matrix formation such as *CDH11* and *LOXL2*, and reduced stemness markers *NOTCH1, HEY1, and SOX2*.

The significantly upregulated DEGs in rGBM samples shown in Fig. 1E were submitted to Gene set enrichment analysis (GSEA) and over representation analysis (ORA) of gene ontology classification biological processes (GO-BP) (Fig. 1F) and were enriched in the terms “ensheathment of neurons”, and “oligodendrocyte differentiation and development” suggesting higher myelination in rGBM compared to pGBM. GO-BP for the DEGs upregulated in pGBM (Fig. 1G) highlighted “cell cycle process” as a distinctive pathway in pGBM suggesting a more proliferative characteristic of tumor cells in pGBM than at its recurrence (Fig. 1H).

Collectively, these targeted transcriptomic results indicate that despite the vast inter- and intratumoral heterogeneity observed in GBM, rGBM and pGBM transcriptomic profiles differ in signatures, suggestive of enhanced myelination and oligodendrocyte differentiation at recurrence.

### Proteomic analysis highlights synaptic signaling as a major feature enriched in rGBM

To reinforce our findings, we performed LC-MS based proteomics in a subset of 6 patient-matched tumor samples (Fig. 2A). This proteomic analysis identified and quantified up to 5214 different proteins PCA separated pGBM and rGBM samples, with only one rGBM sample co-segregating with pGBM samples (Fig. 2B). In agreement, hierarchical clustering of all the assessed proteins (Fig. 2C), and of the high number of differentially expressed proteins (DEPs, Fig. 2D) separated rGBM from pGBM samples. Pathway analysis revealed that significantly upregulated DEPs in rGBM (Fig. 2E, listed in **Table S3**) were enriched in “synaptic signaling” and “vesicle-mediated transport in synapse” in rGBM (Fig. 2F). The top DEPs included in these features comprised NEUG, TPPP, CPLX2, ADCY1 and SYPH.

**Figure 2.**
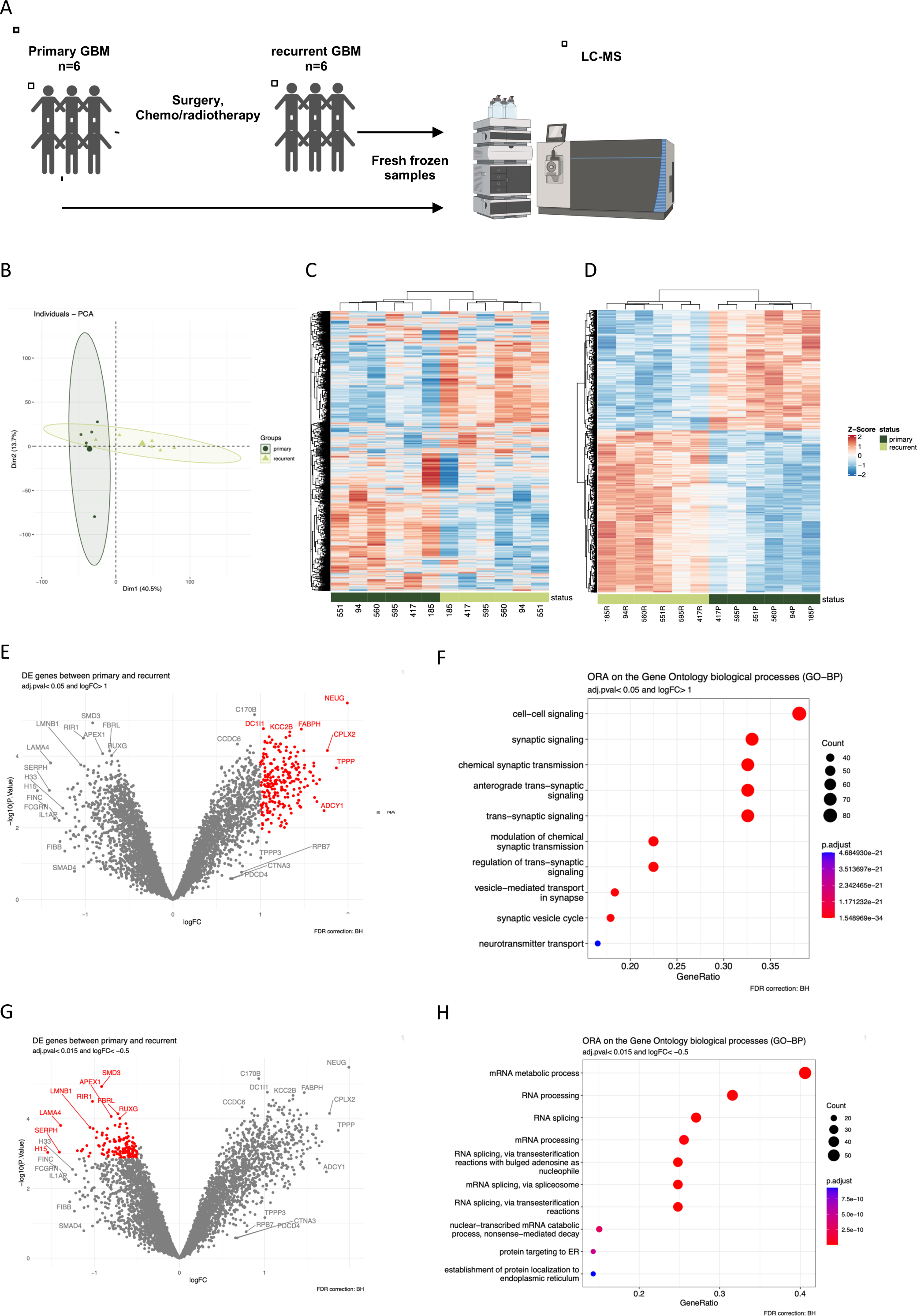
Unbiased LC-MS based proteomic confirms synapse signaling as a major pathway in rGBM. **A)** Experimental setup for LC-MS based proteomic analysis of 6 fresh frozen patient-matched pGBM and rGBM samples (clinical parameters highlighted in Table 1 and Table S1). **B)** PCA plot of all samples included in the analysis, each dot represents a patient, color coded according to pGBM or rGBM. PC1 (40.5% of the variance) and PC2 (13.7% of the variance) are plotted. **C)** Hierarchical clustering heatmap of all the identified proteins across all samples. Z-score transformation was performed for each protein. Proteins with similar expression patterns were clustered together. Relative expressions were scaled from red (high expression) to blue (low expression). Each column represents a sample (dark green: pGBM, light green: rGBM), and each row represents a protein. **D)** Hierarchical clustering heatmap of all DEPs across all samples. **E**) Volcano plot showing significantly differentially expressed proteins in rGBM (red) (adj. p<0.05, logFC>1), **F**) dot-plot of the corresponding enriched biological processes (GO-BP) determined by ORA. **G**) Volcano plot showing significantly differentially expressed proteins in pGBM (red) (adj. p<0.05, logFC>1) and **H**) dot-plot of the corresponding enriched biological processes (GO-BP) determined by ORA.

Pathway analysis using the upregulated DEPs in pGBM (Fig. 2G) highlighted proteins enriched in “mRNA metabolic processes” and “RNA processing” (Fig. 2H), such as SMD3, RIR1 and FBRL.

### High expression of Fcγ receptors (FCGRs) and complement component genes is associated with STTR in rGBM

In order to identify mechanisms leading to recurrence, we focussed on the DEGs upregulated in rGBM. We dichotomized rGBM samples into 2 groups, comprising samples from patients with short TTR (STTR < 309 days) orlong TTR (LTTR ≥ 309 days) (Fig. 3A). We identified 171 significant DEGs between both groups (with p<0.05, Fig. 3B, upper part and **Table S4**). GO-ORA analysis performed with DEGs overexpressed in STTR highlighted “activation of immune response”, “leukocyte-mediated immunity” and “myeloid cell activation”, and DEGs overexpressed in LTTR were enriched in “cell cycle” and “DNA metabolic process”’ as some prominent features (Fig. 3B, middle part). The DEGs present in the most enriched terms (Fig. 3B, lower part), comprise genes coding for the complement system (*C1QA, C1QB, C1QC, C3* and *C3AR1*) and FCG receptors (*FCGR1A (also named CD64), FCGR2A, FCGR3A, FCGR3B*) considering upregulated genes in samples with STTR, and *DNMT3A* and*DNMT1*, *SMARCA4*, *MLH1*, and *TLK2,* considering genes highly expressed in samples with LTTR.

**Figure 3.**
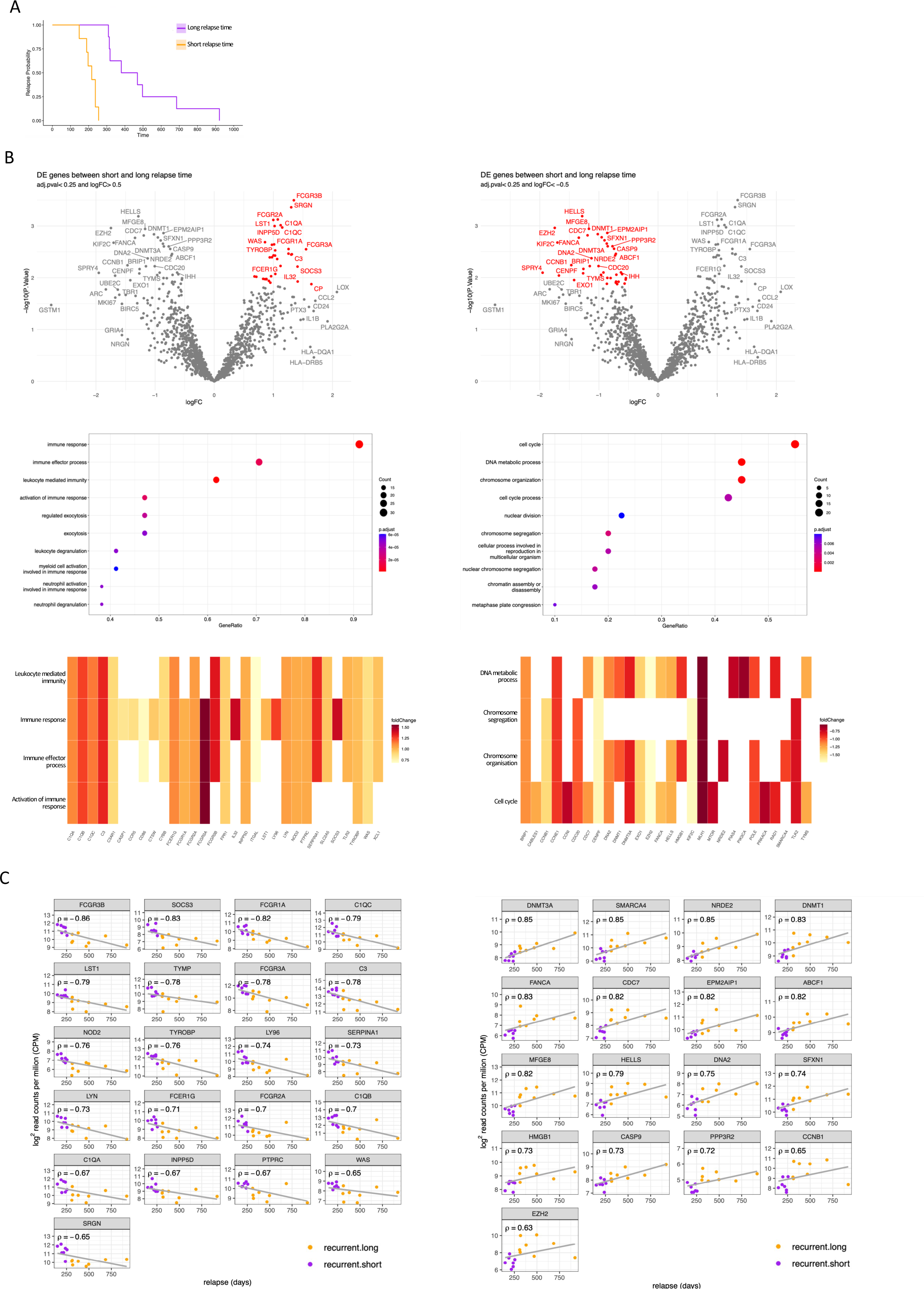
Expressions of immunoglobulin binding receptors and complement system components are higher in the subset of patients with STTR and inversely correlate with TTR. **A)** rGBM samples were subdivided according to their TTR (short time (STTR)<309 days, long time (LTTR)>309 days) and transcriptomics compared between both groups. **B**) Volcano plots showing DEGs in rGBM samples from patients with STTR vs LTTR (upper part). Genes overrepresented in rGBM with STTR (left) or LTTR (right) are highlighted in red (adj. p<0.25, logFC>0.5), dot-plots of the corresponding enriched biological processes (GO-BP) determined by ORA (middle part) and expression of genes enriched in at least one of the 4 most enriched processes (lower part). **C)** Correlation scatterplots of genes negatively (r<-0.63, left) or positively (r>0.63, right) correlated with TTR in rGBM samples.

Analysis of the correlation between expression levels and TTR identified DEGs overexpressed in STTR and strongly negatively associated with TTR, suggesting an unfavorable impact on patients’ overall outcome, whereas genes showing expressions correlating positively with TTR in rGBM (Fig. 3C, and **Fig. S1A and B**) may provide a benefit for patients’ survival.

### Among DEPs, expression of FCGR1 and SHIP1 correlated with STTR

We next correlated the expression levels of DEPs in rGBM with TTR (**Fig. S2A,** listed in **Table S5**). Despite the small number of samples (4 with STTR, and 2 with LTTR), significant negative correlations for FCGR1 and SHIP1 expressions with TTR were observed. Some proteins also displayed higher expressions in LTTR rGBM, however, none of them showed correlations with TTR (**Fig. S2B**).

### Spatial transcriptomic analysis identified cell type composition differences in the CD64+ myeloid population during disease progression

To corroborate and validate our findings, we performed spatial transcriptomics analysis on GFAP^+^ tumor cells and CD64^+^ myeloid cells to gain a more granular impression on iTME/tumor transcriptional state changes on the path from pGBM to rGBM. A cohort of 11 of our patient-paired pGBM and rGBM tissue cores were assembled on a TMA. Tumor cell population was identified by GFAP expression, whereas myeloid cells were identified based on CD64 expression, because of its strong association with tumor relapse in our transcriptomic study. Although fluorescent cells were intimately intertwined in the tumor tissue, both fluorescent signals could be well distinguished, allowing capture of both populations (Fig. 4A). The quality/quantity of the sequencing data was however under the limit of standard for a large number of regions of interest (ROIs), so that only 3 matched pairs, 2 primary samples only and 2 recurrent samples only could be fully exploited for the CD64+ ROIs, and 5 matched pairs, and 5 primary samples only for the GFAP+ ROIs.

**Figure 4.**
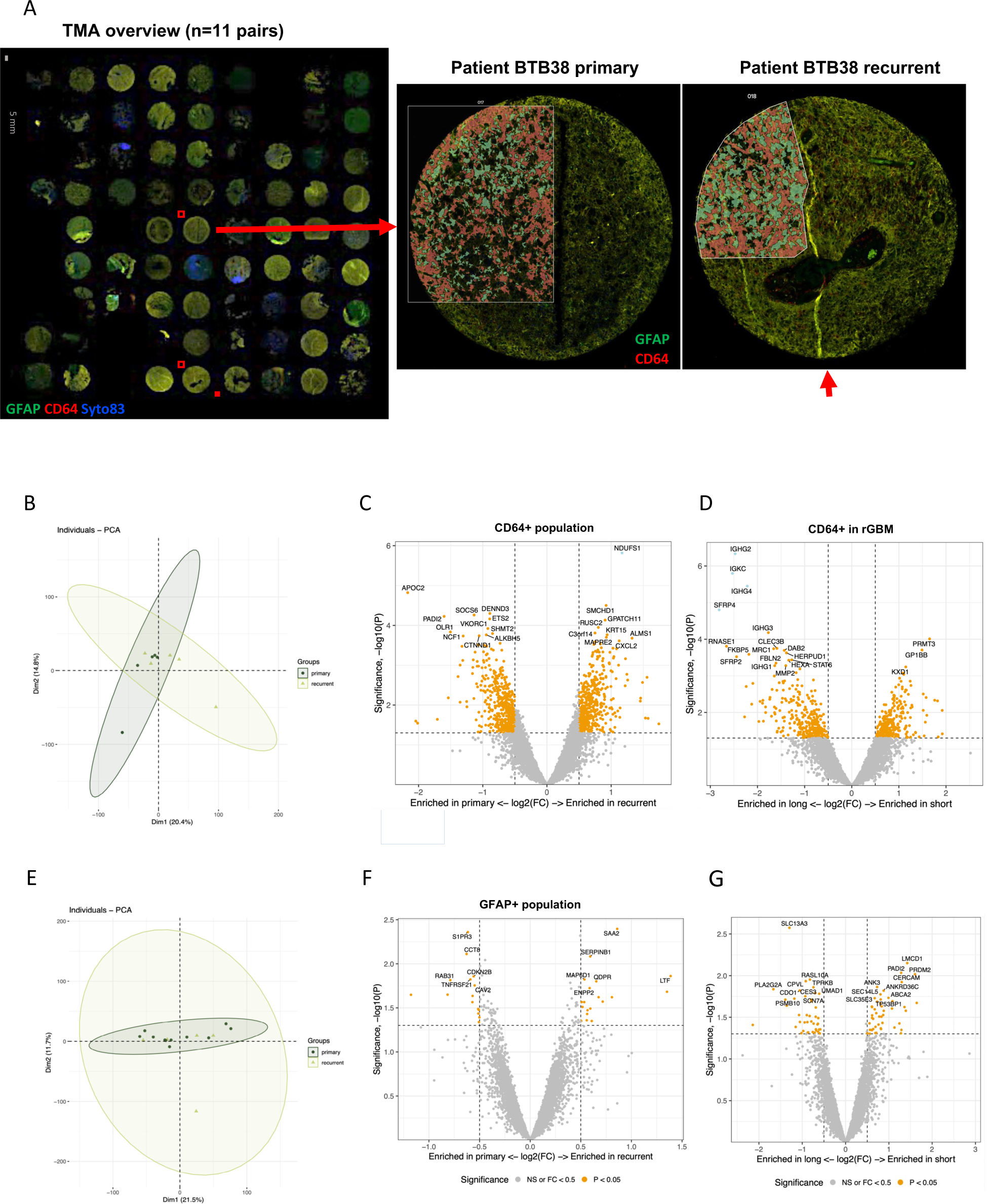
Spatial transcriptomic of CD64- and GFAP-expressing cell populations revealed subtle changes in microglia subtypes in short relapsing rGBM. **A)** Representative TMA section containing 7 pairs of patient-matched pGBM and rGBM samples stained for GFAP (red), CD64 (green) and nuclei (blue) identifying ROIs for GFAP+, CD64+ cellular fractions and nuclei, respectively, and selected for whole transcriptome sequencing. Inserts show magnifications of ROIs from pGBM and rGBM from patient BTB38 as an example. **B)** PCA analysis across CD64+ samples in pGBM and rGBM. **C**) Volcano plot showing DEG between CD64+ cells in pGBM and rGBM (p<0.05 in orange, adj. p<0.05 in light blue, logFC>0.5 or <-0.5). **D**) Volcano plot showing DEG between rGBM samples from patients with STTR and LTTR (p<0.05 in orange, adj. p<0.05 in light blue, logFC>0.5 or <-0.5). **E**) PCA analysis across GFAP+ samples in pGBM and rGBM. **F**) Volcano plot showing DEG between GFAP+ cells in pGBM and rGBM (p<0.05 in orange, logFC>0.5 or <-0.5). **G**) Volcano plot showing DEG between GFAP+ cells in rGBM samples from patients with STTR and LTTR (p<0.05 in orange, logFC>0.5 or <-0.5).

The PCA of the CD64^+^ samples did not show a clear separation of pGBM and rGBM samples (Fig. 4B). We identified several DEGs among CD64^+^ cells in pGBM vs rGBM, however none reached statistical significance (Fig. 4C, p<0.05, but adj. p>0.05, listed in **Table S6**). When comparing the expression of genes between samples with STTR and LTTR in rGBM (listed in listed in **Table S7**), two genes were significantly overexpressed in CD64^+^ population with LTTR, *SFRP4* coding for a Wnt antagonist and *IGHG2* coding for IgG2 (Fig. 4D). As the PCA of GFAP+ ROIs already suggested (Fig. 4E), gene expression analysis failed to uncover significant DEGs between pGBM and rGBM (Fig. 4F, complete list in **Table S8**) and between samples with LTTR and STTR in rGBM (Fig. 4G, **Table S9**).

Since both markers used for spatial transcriptomics are expressed by various cell types, we focussed on changes in specific cell types using cellular deconvolution. Among CD64+ population, we identified mainly quiescent microglia, macrophages and smooth muscle cells in pGBM (Fig. 5A). The most prominent changes in cell type composition upon recurrence were observed in 2 patients with STTR: quiescent microglia disappeared for the benefit of activated microglia, with the appearance of a small proportion of B and T cells. Macrophages were constantly present in pGBM and rGBM, with no major changes in their proportion during disease recurrence. An increase in the proportion of endothelial cells and smooth muscle cells in rGBM samples was also observed (Fig. 5B and **Fig. S5**).

**Figure 5.**
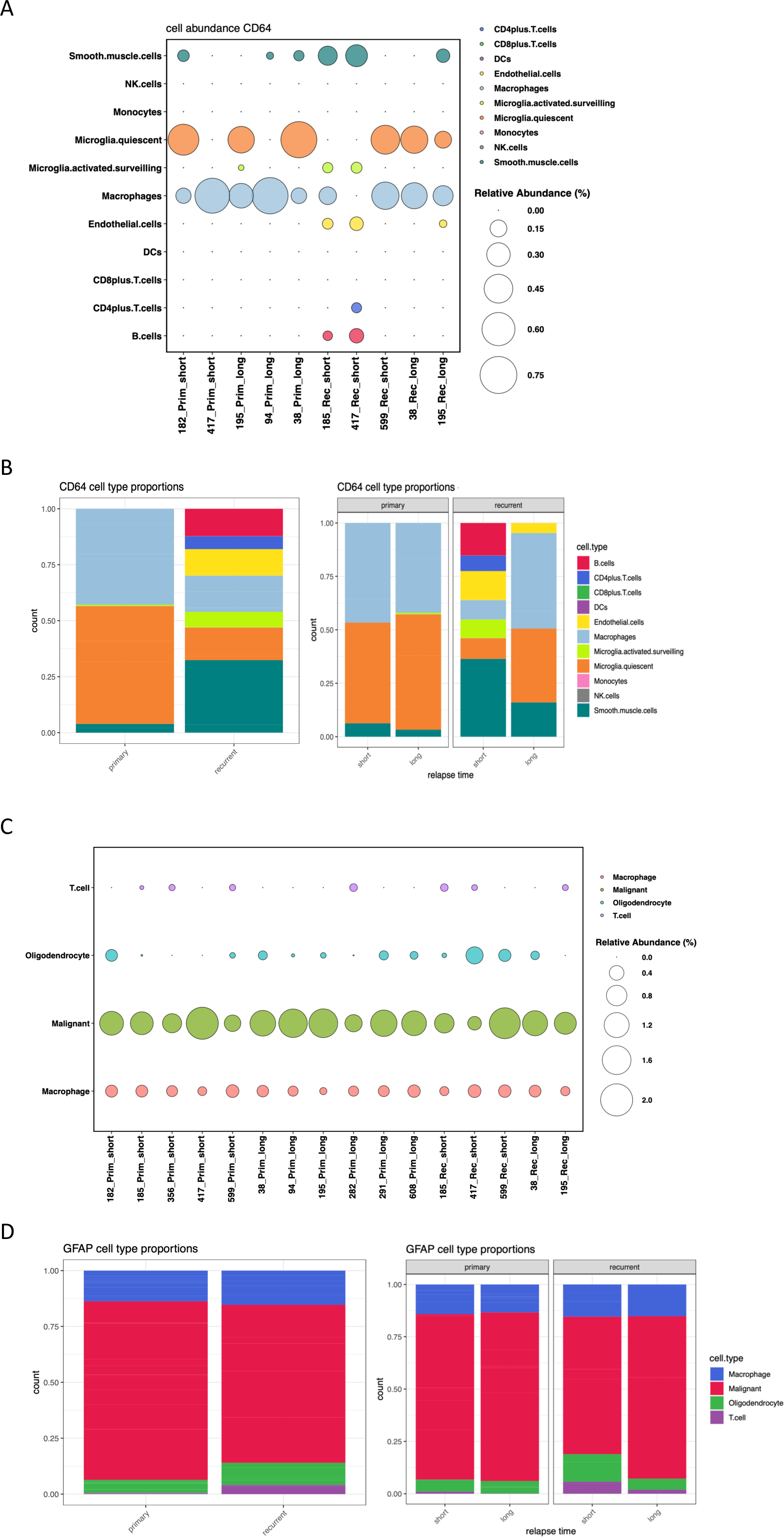
Cellular deconvolution of CD64+ and GFAP+ cell populations revealed subtle changes in microglia subtypes in short relapsing rGBM. **A)** Cell type frequency comparisons between pGBM and rGBM in CD64+ population, subdivided according to short and long TTR. Cell types are labeled and colored based on known cell annotations (using the atlas described by Neftel et al.^50^), and sizes are based on their relative abundance (in %). **B**) Relative cell type abundance in CD64+ population in pGBM and rGBM samples (left chart) and in the rGBM samples grouped according to short or long TTR (right chart). **B)** Cell type frequency comparisons between pGBM and rGBM in GFAP+ populations. Cell types are labeled and colored based on known cell annotations and sizes are based on their relative abundance (in %). **D**) Relative cell type abundance in primary and relapsing GBM samples (left) and in the rGBM samples grouped according to short or long times to relapse (right).

The GFAP-expressing cohort consisted mainly of neoplastic cells, as expected, but also neurons, oligodendrocytes and oligodendrocyte precursor cells (Fig. 5C). When comparing the relative proportion of these cell types in pGBM and rGBM, only minor differences were observed, such as a slight enhancement of the proportion of oligodendrocytes in some rGBM samples (Fig. 5D and **Fig. S4B**).

## Discussion

In this study, patient-matched pGBM and rGBM samples were compared by multi-omics to investigate mechanisms involved in tumor relapse and therapy resistance. We focused on DEGs with a strong association between their expression levels and TTR, to identify potentially targetable markers of quick relapsing tumors.

### Neuro-regenerative functions are activated in rGBM

Our analysis confirmed synapse formation/signaling, myelination and oligodendrocyte differentiation as the main enriched functions in rGBM. Oligodendrocyte precursor cells (OPCs) migrate rapidly to injured sites and occupy regions of traumatic brain injury within 1 day^19^, and may play a role in the development of rGBM, by causing GBM cells to acquire stem cell profiles and chemo-radioresistance^20^. The cellular characteristics of GBM cells resembling stages of oligodendrocyte differentiation^21^ may be exacerbated during recurrence. An increase of the OPC-like population in rGBM was observed also in our spatial transcriptomic analysis further reinforcing this concept. Enrichment of synapse signaling and myelination pathways in rGBM have been described earlier^26^ and suggest regenerative processes of the CNS at the site of recurrence that may facilitate the integration of tumor cells into the TME. Moreover, neurons have been described as important components of the TME regulating malignant growth in an activity-dependent manner^22^.

### Fcγ receptors, complement components and inflammatory mediators influence progression of rGBM

Recurrent tumors that evaded SoC treatment may *de novo* express therapy resistance genes that inversely correlate with TTR. Indeed, we identified 23 genes that were strongly negatively correlated with TTR in our study.Various genes with prognostic value contributing to GBM radioresistance were identified earlier including *FCGR1A* (*CD64*)^23^. FCGR1A is a high-affinity Fcγ receptor expressed on the surface of myelomonocytic and dendritic cells, and may be a prognostic marker related to immune infiltration levels in diverse cancers. Human microglia express FCGR1, FCGR2A, FCGR2B, and FCGR3A albeit at very low levels under normal conditions. The expression is increased on microglia in the CNS of patients with neuroinflammatory diseases such as Alzheimer’s disease^24^ in which the FGCRs play a role in plaque clearance. In GBM, TAMs are considered important drivers of the local immunosuppressive TME involved in progression and resistance to immunomodulating therapeutic strategies^25^. This role is related to their M2 phenotype linked to pro-tumorigenic activity^26^. In contrast, TAMs expressing M1 phenotype secrete proinflammatory cytokines and are prone to eliminate tumor cells. Human GBM harbors a heterogeneous population of M1/M2 TAMs, with a high M1:M2 ratio associated with a better prognosis^27^. Our data, however, suggest that TAMs expressing high levels of FCGRs such as FCGR1 and FCGR3A in rGBM, identified as activated microglia, are associated with STTR.

The expression of *FCGR3A* was negatively correlated with TTR. Expression of FCGR3A is associated with immune cell infiltration, immune checkpoint and DNA mismatch repair genes expression in generalized carcinoma, affecting drug sensitivity, thus serving as a promising biomarker for cancer detection, prognosis, therapeutic design, and follow-up^28^.

Expression of *FCER1G* was also strongly associated with STTR. High, and glioma-grade dependent *FCER1G* levels are associated with immunotherapeutic responses, rendering it a promising immunotherapeutic target^29^.

Complement components C3, C1qA, C1qB and C1qC were highly expressed in rGBM and correlated inversely with TTR. Proteins from the complement system in the CNS are constrained to microglia, oligodendrocytes, astrocytes, and ependymal cells^30^. The complement system is a key player in CNS homeostasis, neurogenesis and regulation of synaptic pruning, as well as in GSC maintenance^31^. Moreover, complement activation promotes carcinogenesis and facilitates proliferation, angiogenesis, resistance to apoptosis, modulation of anti-tumor immunity and activation of invasion and migration, thus providing a tumor-facilitating environment^32^. The C1q family is involved in pathophysiological functions not dependent on complement activation such as neuronal survival and neurite outgrowth *in vitro* but also in the pathogenesis of neurodegenerative diseases such as AD^39^. In GBM, the presence of C1q does not correlate with leukocytic infiltration but is highly concentrated in TAM and necrotic debris^33^, thereby promoting immunosuppression and glioma cell proliferation. C3, in addition to promoting inflammation, directly affects survival, proliferation, migration and stemness of tumor cells^34^.

Interestingly, *TYROBP*, coding for the tyrosine kinase binding protein DAP12, has been shown to switch microglia states from homeostatic to disease-associated, activated microglia in Alzheimer’s disease^35^.

*LST1,* expressed mainly in degranulating macrophages, was also strongly associated with STTR. Interestingly, LST1 supports the formation of tunneling nanotubes (TNTs), structures that are used as routes for brain invasion, proliferation, and interconnection over long distances. Communication between cells through TNT helps tumors acquire resistance to SoC, and inhibition of TNT formation may be a useful novel treatment tool (reviewed in^36^).

Another gene strongly associated with STTR is *SRGN* (serglycin), an intracellular proteoglycan overexpressed in aggressive cancers. Roy et al.^37^ identified serglycin as a potential prognostic marker for glioma.

Abnormal expression of *SOCS1* and *SOCS3* in microglia and astrocytes is associated with poor prognosis of GBM^38^. Moreover, the activation of the JAK/STAT3/SOCS3 signaling pathway can promote the formation of the GBM inflammatory microenvironment. SOCS proteins also participate in the epigenetic regulation of GBM cells via methylation and participate in GBM resistance to chemotherapy^39^.

Interrogating the proteomic results, several proteins encoded by these genes were also associated with TTR, such as proteins from the C1q family, FCGR1/FCGRB, FCGEG, PTPRC and SHIP1. This later protein is a 5′ inositol phosphatase playing crucial roles in cancer signaling and immune response to cancer. SHIP inhibitors (SHIPi) have shown great potential as both chemotherapeutics and immunotherapeutics^40^.

### DNA methylation and other genes favoring LTTR

Genes with higher expression in LTTR rGBM may be associated with a less aggressive phenotype. Higher expression of DNA methyltransferases *DNMT1* and *DNMT3A* were observed in our LTTR patients. Indeed, DNA methylation is associated with TMZ sensitivity in glioma cells^41^. *SMARCA4,* another gene mediating DNA repair and methylation positively correlated with TTR. These observations support a favorable role of DNA methyltransferases in fine-tuning chemoresistance after GBM SOC treatment.

In addition, expression of *MFGE8, SFXN1* and *ABCF1* positively correlated with TTR. *MFGE8* promotes phagocytic removal of apoptotic cells leading to tolerogenic immune responses, and vascular endothelial growth factor (VEGF)-induced angiogenesis^42^. This phagocytic function may help delay tumor relapse. *SFXN1* affects mitochondrial function and iron transport and may also modulate tumor progression in gliomas, thus influencing survival^43^. *ABCF1*, a multidrug resistance gene, confers resistance of cancer cells to a range of anti-cancer agents^51^. The positive correlation between its expression and TTR was unpredicted and would need further investigation.

### Minor transcriptional changes in neoplastic and myeloid cell populations during disease progression

Dissecting the heterogeneity of the cell types present in pGBM and rGBM may be of critical importance to target cell populations in a clinical setting. Using spatial transcriptomics, we explored the transcriptomic changes taking place specifically in the neoplastic cell population and CD64^+^ myeloid cells during disease recurrence.

Gene expression in GFAP^+^ cells, composed mainly of cancer cells, were not significantly changed after SOC therapies. Our study did not highlight specific gene expression changes in GFAP^+^ tumor cells significantly influencing TTR. In addition, no change in the cell types composing GFAP^+^ population could be observed between pGBM and rGBM.

Concerning the CD64^+^ myeloid population, no significant differences in gene expression could be found between pGBM and rGBM. Only when comparing genes expression in rGBM samples with STTR and LTTR, two genes showed significantly elevated levels in samples with LTTR. One of these, *SFRP4*, coding for the Secreted frizzled-related protein 4, a Wnt antagonist, is known to reduce stemness and MMP-2 mediated invasion of glioma cells^44^, and to increase response to chemotherapeutic response of glioma stem cells^45^. The increased expression of *SFRP4* in rGBM with LTTR in our study reinforces a protective role of SFRP4 in GBM.

### Changes in the cell type and cell status during disease progression

The change in the composition of microglial status toward activated microglia in rGBM with STTR indicates that CD64^+^, M1-like microglia may contribute to radio and chemoresistance and recurrence of GBM.

Macrophages are known to express a range of receptors including Fc receptors, including CD64, upregulated under pro-inflammatory conditions^54^. The M1 pro-inflammatory and M2 anti-inflammatory macrophage populations both express CD64, albeit in low levels on M2 macrophages^46^. Surprisingly, M2-like macrophages were not identified in the CD64^+^ cell population in our GBM samples. The absence of CD64 expression by this phenotype may reflect the lack of pro-tumoral macrophages in our analysis, since radioresistant glioblastoma tumor microenvironments have been described to contain more M2 than M1 macrophages^47^. Interestingly, CD64-directed immunotherapeutic agents have emerged for the treatment of macrophage-mediated chronic inflammatory diseases^57^. This receptor may therefore represent a new potential therapeutic target for activated microglia in GBM patients.

Cells in the CD64^+^ population also showed expression of B cell markers in 2 out of 3 short relapsing rGBM samples. Interestingly, B cells have been reported to be part of the GBM immune landscape and are characterized by immunosuppressive activity toward activated CD8^+^ T cells, through overexpression of inhibitory molecules such as PD-L1 and CD155, and production of the immunosuppressive cytokines TGFβ and IL10^48^.

We noted an increase in the proportion of endothelial cells in the CD64+ cell population in rGBM samples. Since endothelial cells are not known to express CD64, this might originate from contamination by blood vessels due to strong adhesion of CD64+ monocytes to vascular structures after their activation^49^.

This study has several limitations. The small cohort of patient-paired GBM samples, and grouping our samples according to TTR further reduced the confidence of the findings. But, despite the small number of patients, we found strong correlations between genes/proteins expression and TTR. Correlation does not imply causation, but highlights a relationship between both factors. Therefore, it would be essential to validate this association in an *in vivo* model to clearly define the role of CD64 in GBM recurrence.

In conclusion, developing new drugs for treating GBM patients is challenging. Multi-omic analyses are one powerful tool to identify specific targets at various disease stages. Here we identified genes and proteins that are changed from initial diagnosis to recurrence. Due to the tremendous heterogeneity of GBM tumors, we focussed our transcriptomic analysis on immune and neuroinflammatory targets and concentrated our attention on molecular players favoring recurrence, that undeniably represent the preferred targets for new therapies. The observation that CD64 and C1q strongly influence TTR on gene as well as protein levels pinpoints them as potential new targets. Therefore, we propose CD64^+^ myeloid cells present in the GBM microenvironment as a new research center of attention in better understanding resistance processes and preventing patients from relapse.

## Funding

The Swiss Life Jubiläumsstiftung (1138 to T.S., 1196 to T.S.); Krebsliga beider Basel (5072-02-2020 to T.S. and G.H.).

## Acknowledgments

We thank Dr. Emmanuel Contassot (Department of Biomedicine, University of Basel, Basel), Dr. Katarzyna Buczak (Proteomics Core facility, Biozentrum, University of Basel, Basel), Dr. Ida AK Nilsson and Lars Selander (Department of Molecular Medicine and Surgery, Karolinska Institutet, Stockholm, Sweden) for providing reagents and important technical support. We acknowledge the Neurosurgery Department, University Hospital Basel, Basel, Switzerland for supplying with fresh tumor material removed during surgery. We also thank the Neuropathology Department, Institute of Pathology, University Hospital Basel, Basel, Switzerland for providing us with FFPE tumor material.

## Author contribution

T.S. and M.-F.R. conceived and designed the experiments. T.S., M.-F.R, T.A.M., M.M., C.D., A.G. performed experiments. T.S., M.-F.R, S.H., J.R. analyzed the data. T.S., M.-F.R, S.H. and G.H. wrote the manuscript. D.K., B.B. and P.S. provided technical support. All authors read and approved the manuscript.

## Declaration of interest

All other authors declare they have no competing interests.

## Notes

### Competing Interest Statement

The authors have declared no competing interest.

