## Supplementary methods and figures for "Multidimensional analysis of matched primary and recurrent glioblastoma identifies Fcγ receptors upregulation on microglia as a contributor of tumor recurrence"

**Supplementary Methods 1**

Peptide preparation and Liquid chromatography/mass spectrometry

Protein samples (approx. 20 mg tumor tissue) were grinded and lysed by sonication. After measuring the total protein concentration, proteins were reduced, alkylated, and subsequently tryptic digested, dried under vacuum, and stored at -20°C until further use.

Tandem mass tag (TMT) proteomics

Sample aliquots comprising 10 µg of flow through peptides were labeled with isobaric tandem mass tags (TMTpro 16-plex, Thermo Fisher Scientific). Peptides were resuspended in 10 µl labeling buffer (2 M urea, 0.2 M HEPES, pH 8.3) by sonication and 2.5 µL of each TMT reagent was added to the individual peptide samples followed by a 1 h incubation at 25°C shaking at 500 rpm. After quenching of the labelling reaction, all samples were pooled and the pH was increased to 12 to remove TMT labels linked to peptide hydroxyl groups. The reaction was stopped by acidification. Finally, peptide samples were desalted using a C18 reverse-phase spin columns (Macrospin, Harvard Apparatus) according to the manufacturer's instructions and dried under vacuum.

High performance liquid chromatography (HPLC) fractionation

TMT-labeled peptides were fractionated by high-pH reversed phase separation using a XBridge Peptide Ethylene Bridged Hybrid (BEH) C18 column (3,5 µm, 130 ångstrom (Å), 1 mm x 150 mm, Waters) on an Agilent 1260 Infinity HPLC system. Peptides were loaded in buffer A (20 mM ammonium formate in water, pH 10) and eluted using a two-step linear gradient from 2% to 10% in 5 minutes and then to 50% buffer B (20 mM ammonium formate in 90% acetonitrile, pH 10) over 55 minutes at a flow rate of 42 µl/min. Elution of peptides was monitored with an ultraviolet (UV) detector (215 nm, 254 nm) and a total of 36 fractions were collected, pooled into 12 fractions using a post-concatenation strategy and dried under vacuum.

LC-MS/MS analysis

Dried peptides were resuspended in 0.1% aqueous formic acid (Buffer A) and subjected to LC–MS/MS analysis using a Q Exactive HF Mass Spectrometer fitted with an EASY-nLC 1000 (both Thermo Fisher Scientific) at 60°C. Peptides were resolved using a RP-HPLC column (75µm × 30cm) with C18 resin (ReproSil-Pur C18–AQ, 1.9 µm resin; Dr. Maisch GmbH) at a flow rate of 0.2 $\mu Lmin^{-1}$. The following gradient was used for peptide separation: from 5% solvent B (80% acetonitrile, 0.1% formic acid in water) to 15% solvent B over 10 min, to 30% solvent B over 60 min, to 45 % solvent B over 20 min, to 95% solvent B over 2 min, followed by 18 min at 95% solvent B.

The mass spectrometer was operated in DDA mode with a total cycle time of approximately 1 s. Each MS1 scan was followed by high-collision-dissociation (HCD) of the 10 most abundant precursor ions with dynamic exclusion set to 30 seconds.

The acquired raw-files were analysed using the SpectroMine software (Biognosis AG, Schlieren, Switzerland). Spectra were searched against a human database consisting of 20742 protein sequences (downloaded from Uniprot on 20190307) and 392 commonly observed contaminants. Raw reporter ions intensities of protein group specific PSMs were exported and used for quantification.

For each TMTpro 16-plex experiment, raw PSMs intensities were summed within a protein group and normalized using quantiles method.

**Supplementary methods 2**

Spatial transcriptomic

Slide-mounted FFPE TMA was processed for antigen retrieval using a heat induced epitope retrieval protocol for 20 minutes followed by a 5-min wash with 1µg/mL proteinase K solution. TMA was next incubated overnight with GeoMx RNA detection probes containing-photocleavable oligos. Next, slides were stained using a mixture of conjugated antibodies for identification of tissue morphology: anti-GFAP-594 (Novus Biologicals, NBP2-33184DL594; 1:1000) and anti-CD64 (purified with Abcam purification kit (ab102784) and conjugated with AlexaFluor647 (ab269235); 1:50) for the labeling of tumor cells and myeloid cells, respectively. In addition, Syto83 was used for DNA labeling (ThermoFisher Scientific, USA, 1:25). Stained slides were loaded onto a GeoMx instrument and scanned. Custom masks were created to define ROIs of interest for UV illumination. Once ROIs were defined, each area of interest was exposed to 385 nm light, releasing the indexed oligonucleotides which were collected and deposited in a 96-well plate for subsequent processing. Sequencing libraries were generated by PCR from the oligos and sequenced using Illumina NovoSeq according to the manufacturer’s protocol.

**Figure S1**

Barplots of the most differentially expressed genes in rGBM samples with STTR (**A**) and LTTR (**B**).

**Figure S2**

Barplots (left) and correlation scatterplots (right) of the mostly differentially expressed proteins in STTR (**A**) and LTTR rGBM samples (**B**).

**Figure S3**

Representative histological section of the TMA containing patient-paired pGBM and rGBM FFPE tissue cores and 2 liver tissue cores (**A**), and corresponding sample names (**B**).

**Figure S4**

Cell type abundances in counts (left) and as proportions (right) of CD64+ cell population (**A**) and GFAP+ cell populations (**B**) in individual tumor samples.

**Supplementary Table legends**

**Table 1S**. Patients' characteristics including diagnostic and therapeutic data.

**Table 2S.** Significant DEGs between pGBM and rGBM

**Table 3S.** Significant DE proteins between pGBM and rGBM

**Table 4S.** DEGs between short and long relapsing rGBM samples

**Table 5S.** DE proteins between short and long relapsing rGBM samples

**Table 6S.** DEGs between pGBM and rGBM in CD64+ cell population

**Table 7S.** DEGs between short and long relapsing rGBM samples in the CD64+ cell population

**Table 8S.** DEGs between pGBM and rGBM in GFAP+ cell population

**Table 9S**. DEGs between short and long rGBM samples in the GFAP+ cell population

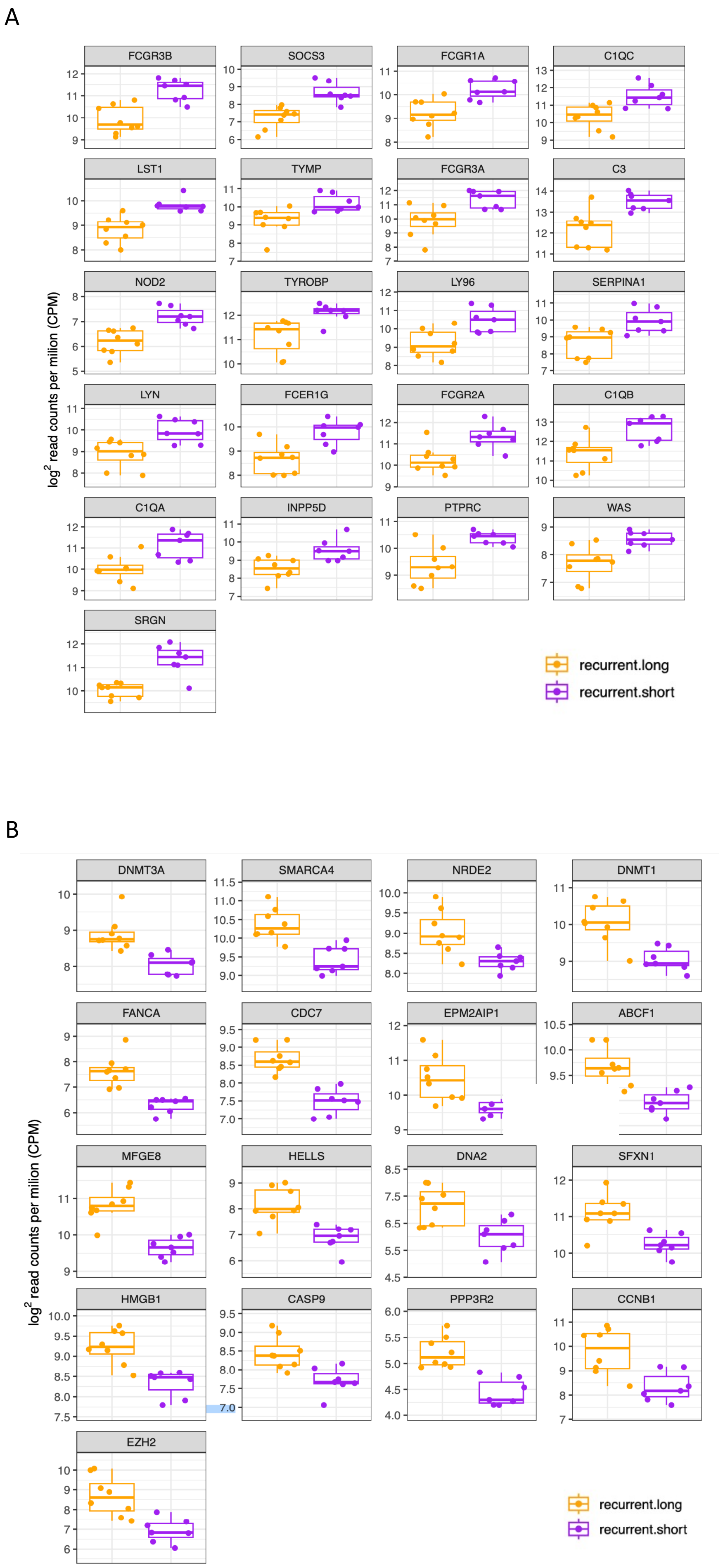


### Supplementary Figure 1

A

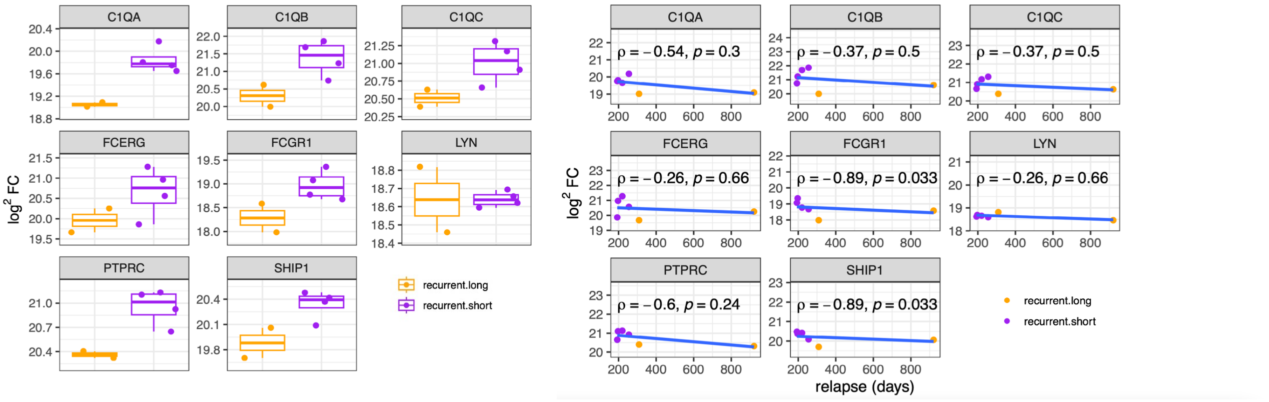


B

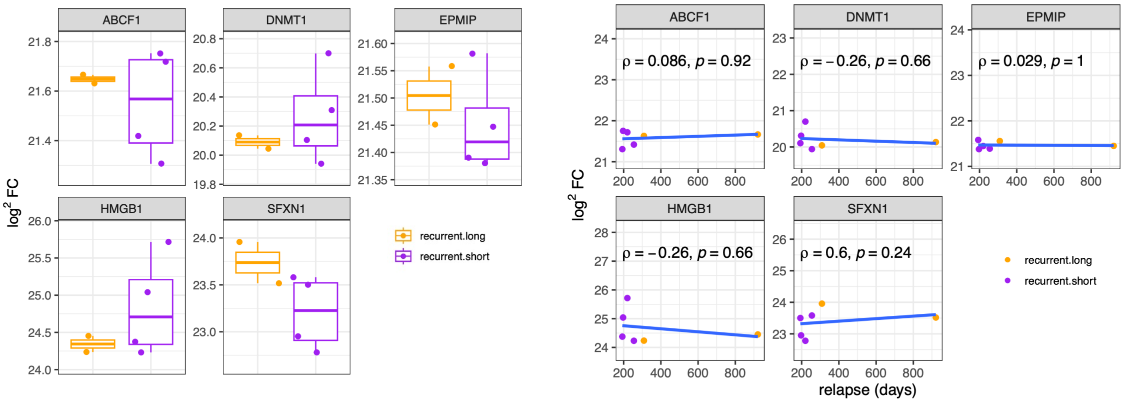


Supplementary Figure 2

A

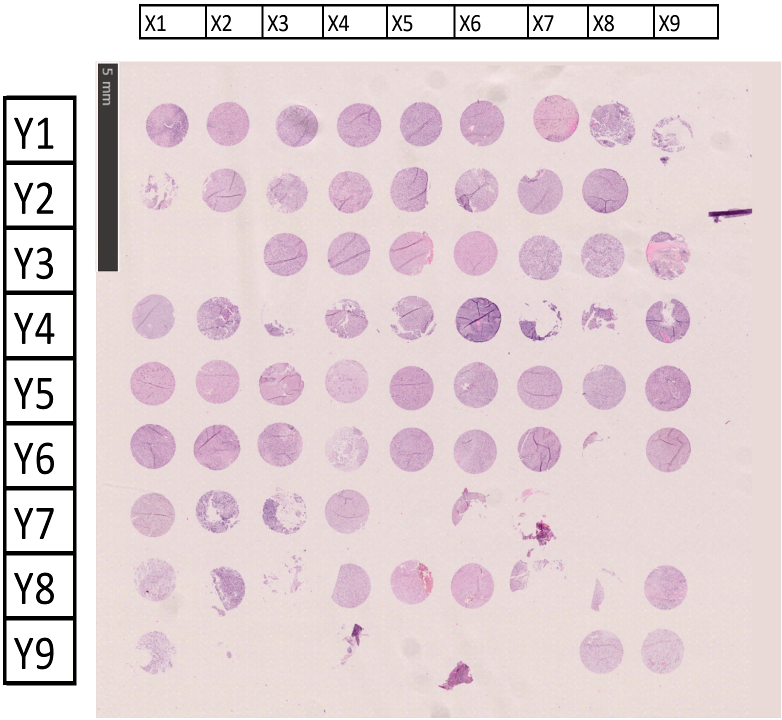


B

| | X1 | X2 | X3 | X4 | X5 | X6 | X7 | X8 | X9 |
|---|---|---|---|---|---|---|---|---|---|
| Y1 | liver | liver | 195P | 195P | 195P | 195P | 195P | 195P | |
| Y2 | | 356P | 356P | 356P | 356P | 282P | 282P | 282P | |
| Y3 | | | 182P | 182P | 182P | 182P | 185P | 185P | 185P |
| Y4 | 185P | 185R | | 185R | 185R | 417P | | | 417P |
| Y5 | 417R | 417R | 417R | 417R | 38P | 38P | 38P | 38P | 38R |
| Y6 | 38R | 38R | 38R | 94P | 94P | 94P | 94P | | 94R |
| Y7 | 94R | 291P | | 291P | | 291R | | | |
| Y8 | 599R | 599R | 599R | 599R | 599P | 599P | | | 608P |
| Y9 | 608P | | | 608R | | | | 182R | 182R |

Supplementary Figure 3

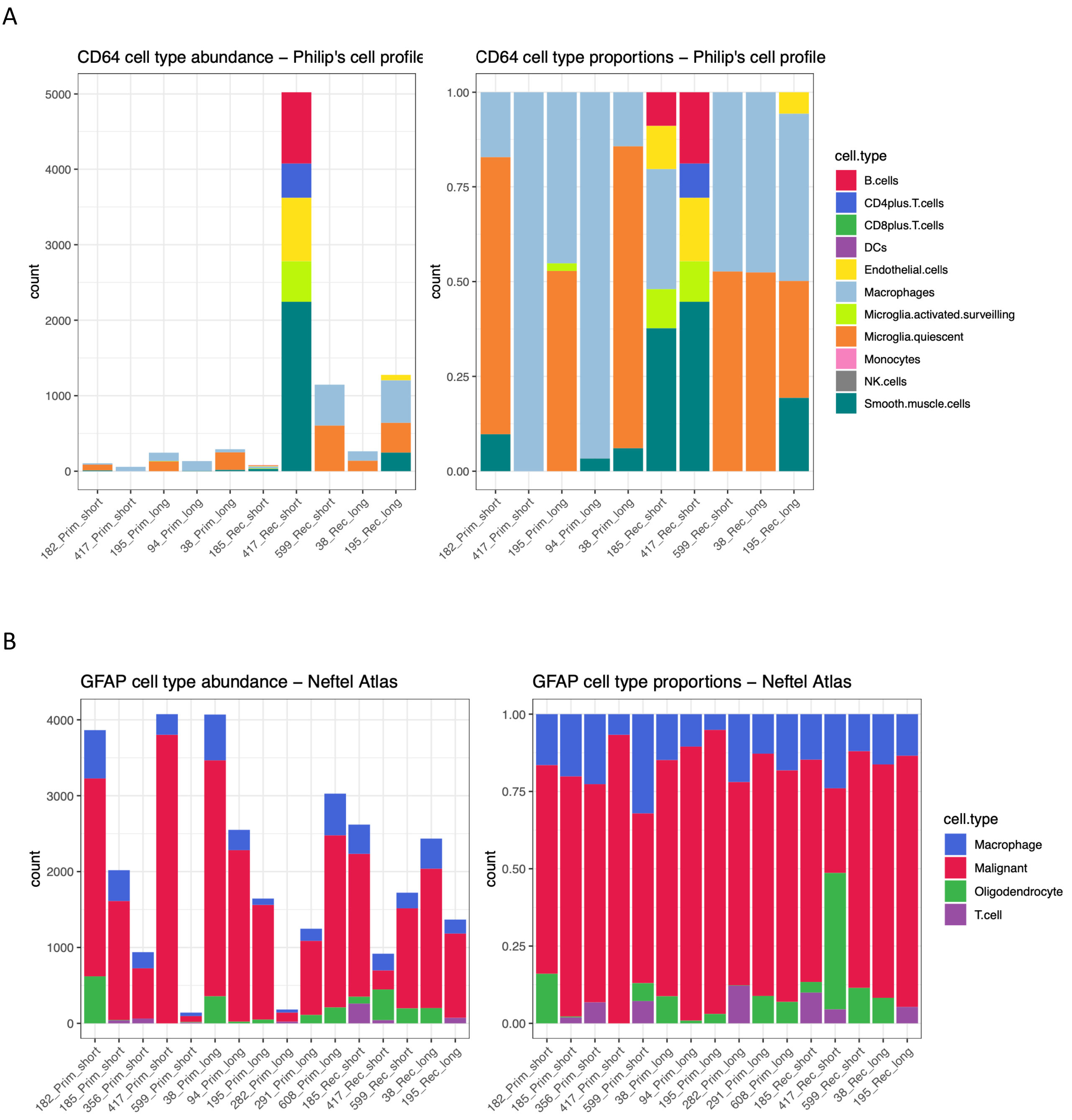


Supplementary Figure 4

Table S1

| BTB# | Birth Year | Age at diagnosis | Sex | Diagnosis | e of 1st surg | IDH status |
|---|---|---|---|---|---|---|
| 182 | 1946 | 65 | M | GBM IV | 07.09.2011 | wt |
| 195 | 1949 | 62 | F | GBM IV | 01.12.2011 | wt |
| 356 | 1941 | 73 | M | GBM IV | 16.07.2014 | wt |
| 417 | 1968 | 47 | M | GBM IV | 28.09.2015 | wt |
| 496 | 1953 | 63 | M | GBM IV | 29.11.2016 | wt |
| 185 | 1958 | 51 | F | GBM IV | 13.09.2011 | wt |
| 291 | 1974 | 39 | M | GBM IV | 12.07.2013 | wt |
| 532 | 1958 | 59 | M | GBM IV | 21.08.2017 | wt |
| 545 | 1953 | 64 | M | GBM IV | 11.01.2018 | wt |
| 38 | 1954 | 56 | M | GBM IV | 16.03.2010 | wt |
| 94 | 1946 | 64 | M | GBM IV | 11.10.2010 | wt |
| 560 | 1973 | 45 | F | GBM IV | 18.05.2018 | wt |
| 551 | 1967 | 51 | M | GBM IV | 20.03.2018 | wt |
| 595 | 1961 | 57 | M | GBM IV | 23.04.2019 | wt |
| 599 | 1956 | 63 | M | GBM IV | 08.05.2019 | wt |
| 608 | 1962 | 56 | M | GBM IV | 06.08.2019 | wt |
| 623 | 1965 | 52 | F | GBM IV | 18.12.2019 | wt |
| 282 | 1950 | 63 | M | GBM IV | 05.04.2013 | wt |

NA: not available
ND: nor determined
mes: mesenchymal

| MGMT-promotor methylation | Adjuvant Treatment | ate of 2nd surge |
|---|---|---|
| negative | CRT | 14.03.2012 |
| negative | CRT | 11.04.2013 |
| positive | CRT | 10.03.2015 |
| negative | CRT | 12.04.2016 |
| positive | CRT | 13.03.2018 |
| negative | CRT | 18.07.2012 |
| negative | CRT | 28.05.2015 |
| NA | CRT | 05.09.2018 |
| positive | CRT | 08.06.2018 |
| negative | CRT | 24.01.2011 |
| positive | CRT | 19.04.2013 |
| negative | CRT | 27.11.2018 |
| negative | CRT | 30.11.2018 |
| negative | CRT | 29.11.2019 |
| negative | CRT | 11.12.2019 |
| negative | CRT | 19.06.2020 |
| positive | CRT | 11.08.2020 |
| negative | CRT | 21.03.2014 |

| time to relapse (d) | short or long relapse | Date of death | al since 1st surg |
|---|---|---|---|
| 189 | short | 04.11.2012 | 424 |
| 497 | long | 01.11.2013 | 701 |
| 237 | short | 08.10.2016 | 815 |
| 197 | short | 10.05.2017 | 590 |
| 469 | long | 24.12.2019 | 1120 |
| 309 | long | 08.03.2013 | 542 |
| 685 | long | 16.07.2016 | 1100 |
| 380 | long | 01.04.2019 | 588 |
| 148 | short | 23.04.2019 | 467 |
| 314 | long | 27.05.2011 | 437 |
| 921 | long | 17.04.2014 | 1284 |
| 193 | short | 09.08.2019 | 448 |
| 255 | short | 11.07.2019 | 478 |
| 220 | short | 17.02.2020 | 300 |
| 217 | short | 10.02.2020 | 278 |
| 318 | long | alive | NA |
| 237 | short | 29.10.2020 | 316 |
| 350 | long | 04.01.2015 | 639 |

| Subtype | Transcriptomic | Proteomics | Spatial transcriptomic |
|---|---|---|---|
| ND | Yes | No | Yes |
| ND | Yes | No | Yes |
| ND | Yes | No | Yes |
| ND | Yes | Yes | Yes |
| Mes | Yes | No | No |
| ND | Yes | Yes | Yes |
| ND | Yes | No | Yes |
| Classical | Yes | No | No |
| Proneural | Yes | No | No |
| ND | Yes | No | Yes |
| ND | Yes | Yes | Yes |
| Classical | No | Yes | No |
| Classical | Yes | Yes | No |
| Proneural | Yes | Yes | No |
| Classical | Yes | No | No |
| Proneural | Yes | No | Yes |
| Mes | Yes | No | No |
| ND | No | No | Yes |

Table S2

| ID | logFC | P.Value | adj.P.Val |
|---|---|---|---|
| MOBP | 3.6610695 | 0.000038 | 0.0043919 |
| OPALIN | 3.2137159 | 0.000146 | 0.0064243 |
| SYN2 | 3.029232 | 0.0003187 | 0.008909 |
| NEFL | 3.015474 | 0.000916 | 0.0165641 |
| PLP1 | 3.0010334 | 0.0000589 | 0.0043919 |
| PACSIN1 | 2.8284626 | 0.0006728 | 0.0133781 |
| MOG | 2.6615933 | 0.0000943 | 0.0054131 |
| MAG | 2.6541426 | 0.000012 | 0.0036156 |
| NRGN | 2.5891733 | 0.0010841 | 0.0174441 |
| SLC17A7 | 2.4848483 | 0.0021308 | 0.0257674 |
| MAL | 2.3815942 | 0.0000481 | 0.0043919 |
| GJB1 | 2.3242862 | 0.0000599 | 0.0043919 |
| TMEM144 | 2.2754256 | 0.0000084 | 0.0036156 |
| RASGRF1 | 2.2702965 | 0.0000214 | 0.0041864 |
| MYRF | 2.2027901 | 0.0002421 | 0.0087748 |
| GRIN2A | 2.0775256 | 0.0002593 | 0.0087748 |
| TUBB4A | 1.9627976 | 0.0002293 | 0.0087748 |
| BCAS1 | 1.7757596 | 0.0021668 | 0.0257674 |
| FA2H | 1.7533498 | 0.0015089 | 0.0199179 |
| S1PR5 | 1.752579 | 0.000225 | 0.0087748 |
| BOK | 1.7451362 | 0.0002906 | 0.008909 |
| FCGR2B | 1.7300455 | 0.0003316 | 0.0089338 |
| SNCA | 1.7262228 | 0.0011427 | 0.0175845 |
| GRM3 | 1.7174971 | 0.0011305 | 0.0175845 |
| CYP7B1 | 1.6778633 | 0.000103 | 0.005439 |
| UGT8 | 1.6705542 | 0.0036778 | 0.0351787 |
| LYZ | 1.6202701 | 0.0003208 | 0.008909 |
| RBFOX3 | 1.5502365 | 0.003255 | 0.0331479 |
| NINJ2 | 1.5397771 | 0.0000991 | 0.005439 |
| MS4A4A | 1.5295684 | 0.0000222 | 0.0041864 |
| GPR62 | 1.4996866 | 0.0012613 | 0.0187074 |
| CDKN1C | 1.4339642 | 0.0007823 | 0.0147521 |
| DEPTOR | 1.355037 | 0.0003412 | 0.0090082 |
| LACC1 | 1.3514589 | 0.0009721 | 0.0171087 |
| MS4A6A | 1.3363749 | 0.0001317 | 0.0061198 |
| GOT1 | 1.3339448 | 0.0010323 | 0.0172479 |
| HLA.DPB1 | 1.3311232 | 0.0000433 | 0.0043919 |
| PLEKHB1 | 1.3115932 | 0.0009465 | 0.016884 |
| GZMA | 1.2715446 | 0.001352 | 0.0193327 |
| SLC6A1 | 1.2509928 | 0.0041551 | 0.0383545 |
| NWD1 | 1.2203329 | 0.0026507 | 0.0296515 |
| HLA.DPA1 | 1.1748478 | 0.0003848 | 0.0094064 |
| FPR3 | 1.1734583 | 0.0004008 | 0.0096189 |
| MRC1 | 1.1664528 | 0.0048054 | 0.0422875 |
| CHN2 | 1.1575189 | 0.0002675 | 0.0087748 |
| CTSS | 1.150681 | 0.0001345 | 0.0061198 |

| | | | |
|---|---|---|---|
| ADRA2A | 1.1264576 | 0.000038 | 0.0043919 |
| IGSF6 | 1.1202495 | 0.003073 | 0.0324111 |
| RAB6B | 1.1030419 | 0.0016619 | 0.0214753 |
| CD84 | 1.0822655 | 0.0002596 | 0.0087748 |
| HDAC11 | 1.0812568 | 0.0022443 | 0.0262163 |
| HLA.DRA | 1.0796346 | 0.0013774 | 0.0193327 |
| IFIT2 | 1.0774128 | 0.0051629 | 0.0425456 |
| DAB2 | 1.0676918 | 0.0000317 | 0.0043919 |
| TMC7 | 1.0676688 | 0.0005748 | 0.0123771 |
| IFIT1 | 1.0639317 | 0.0005814 | 0.0123771 |
| IFIT3 | 1.0520409 | 0.0048726 | 0.0425173 |
| HLA.DMB | 1.0350029 | 0.0003206 | 0.008909 |
| NQO1 | 1.0287253 | 0.0024636 | 0.0282775 |
| CMKLR1 | 1.0170103 | 0.0000082 | 0.0036156 |
| SIGLEC1 | 1.0090375 | 0.000478 | 0.0112196 |
| IL18 | 1.0042034 | 0.0000879 | 0.0052767 |
| CD68 | 1.0016684 | 0.0000573 | 0.0043919 |
| P2RY13 | 0.9962397 | 0.00108 | 0.0174441 |
| GSN | 0.9929005 | 0.0010178 | 0.017224 |
| C7 | 0.9791482 | 0.0018461 | 0.0227739 |
| SIRPA | 0.9764728 | 0.0000582 | 0.0043919 |
| LGMN | 0.9666401 | 0.0006556 | 0.0133781 |
| CD48 | 0.9482833 | 0.0032395 | 0.0331479 |
| HLA.DMA | 0.9367397 | 0.0013184 | 0.0191244 |
| SERPINF1 | 0.9297496 | 0.0000872 | 0.0052767 |
| FCGR1A | 0.9231869 | 0.0006857 | 0.0133781 |
| GPR34 | 0.9215566 | 0.0016402 | 0.0214366 |
| IRF8 | 0.9214562 | 0.0003723 | 0.0092725 |
| CD45RO | 0.918634 | 0.0012323 | 0.0186969 |
| MPEG1 | 0.9155204 | 0.002582 | 0.0291301 |
| PILRA | 0.9103417 | 0.0017986 | 0.0226104 |
| ITPK1 | 0.890311 | 0.0027993 | 0.0303858 |
| MEF2C | 0.88362 | 0.0051854 | 0.0425456 |
| SIGLEC8 | 0.8827137 | 0.0036261 | 0.0351677 |
| PLA1A | 0.8600279 | 0.0005576 | 0.0123771 |
| BIN1 | 0.8563437 | 0.0038501 | 0.0365197 |
| AK1 | 0.8256567 | 0.00365 | 0.0351677 |
| TMCC3 | 0.8133482 | 0.0045556 | 0.0411873 |
| PIK3R5 | 0.8083444 | 0.0039286 | 0.0365197 |
| IL2RG | 0.8006906 | 0.0049466 | 0.0425173 |
| MAP3K8 | 0.7998181 | 0.0016757 | 0.0214753 |
| HAVCR2 | 0.7962987 | 0.0014701 | 0.0196019 |
| ICOSLG | 0.7917089 | 0.004252 | 0.0389763 |
| MERTK | 0.7905594 | 0.0050486 | 0.0425456 |
| PRR5 | 0.7816111 | 0.0031332 | 0.0325658 |
| TGFBR2 | 0.7804756 | 0.0013171 | 0.0191244 |
| PRKACB | 0.7791308 | 0.0059592 | 0.0476738 |
| RGL1 | 0.7382527 | 0.0003557 | 0.0092068 |

| | | | |
|---|---|---|---|
| SLCO2B1 | 0.7320257 | 0.0009868 | 0.0171396 |
| BCL2 | 0.670168 | 0.0010629 | 0.0174441 |
| LAIR1 | 0.6644222 | 0.0020461 | 0.0250081 |
| PPARG | 0.6642208 | 0.002154 | 0.0257674 |
| TREM2 | 0.661085 | 0.0047524 | 0.0422875 |
| TLR7 | 0.6363736 | 0.0030704 | 0.0324111 |
| GIMAP4 | 0.6098413 | 0.0008668 | 0.0158922 |
| CLEC7A | 0.6005086 | 0.0051678 | 0.0425456 |
| CD47 | 0.5509627 | 0.0043515 | 0.0396135 |
| PIK3CG | 0.5105123 | 0.0047841 | 0.0422875 |
| GIMAP6 | 0.4953028 | 0.0051893 | 0.0425456 |

Table S2

| ID | logFC | P.Value | adj.P.Val |
|---|---|---|---|
| VEGFA | -2.6273842 | 0.0001317 | 0.0061198 |
| SOX4 | -1.6060099 | 0.0017467 | 0.0221699 |
| SOX11 | -1.5418043 | 0.0027656 | 0.0303858 |
| EZH2 | -1.4208313 | 0.0010147 | 0.017224 |
| RRM2 | -1.4174666 | 0.0022159 | 0.0261159 |
| MMP9 | -1.4137131 | 0.0013844 | 0.0193327 |
| TMEM100 | -1.3636397 | 0.0059153 | 0.0476106 |
| UBE2C | -1.336422 | 0.0056186 | 0.0455003 |
| IL1RAP | -1.3177288 | 0.0070453 | 0.053447 |
| LMNB1 | -1.289665 | 0.002541 | 0.0289149 |
| BRIP1 | -1.2114671 | 0.0002577 | 0.0087748 |
| JAG1 | -1.2003487 | 0.0012589 | 0.0187074 |
| SOX2 | -1.1952179 | 0.000109 | 0.0055342 |
| TNFRSF12A | -1.1778646 | 0.0008502 | 0.0158064 |
| LIF | -1.1674342 | 0.003923 | 0.0365197 |
| VCAN | -1.1638977 | 0.0005008 | 0.0113972 |
| NOTCH1 | -1.1606587 | 0.0000879 | 0.0052767 |
| HEY1 | -1.1542965 | 0.0000435 | 0.0043919 |
| DOT1L | -1.1380127 | 0.0023577 | 0.0272993 |
| ANLN | -1.1289066 | 0.0014353 | 0.0193327 |
| CCND2 | -1.0855142 | 0.0027254 | 0.0302313 |
| EIF4EBP1 | -1.0759933 | 0.0003676 | 0.0092725 |
| TCF3 | -1.0628581 | 0.0000477 | 0.0043919 |
| NDUFA4L2 | -1.0444024 | 0.0002726 | 0.0087748 |
| POLE | -1.0377706 | 0.000324 | 0.008909 |
| CDK2 | -1.0122706 | 0.0033352 | 0.0336068 |
| LOXL2 | -1.0011379 | 0.0000137 | 0.0036156 |
| CHEK1 | -0.9837248 | 0.003459 | 0.0343296 |
| TYMS | -0.9694236 | 0.006372 | 0.0500659 |
| MCM2 | -0.9670817 | 0.0014001 | 0.0193327 |
| CDH11 | -0.9644845 | 0.0000114 | 0.0036156 |
| DNMT1 | -0.9540615 | 0.0002876 | 0.008909 |
| CD276 | -0.9527916 | 0.0002719 | 0.0087748 |
| SLC16A1 | -0.9209813 | 0.0006433 | 0.0133781 |
| BMP2 | -0.8980379 | 0.0004845 | 0.0112196 |
| BRD4 | -0.8864813 | 0.0030098 | 0.0322998 |
| BLM | -0.876337 | 0.0032646 | 0.0331479 |
| DLL4 | -0.8714463 | 0.0006892 | 0.0133781 |
| SLC7A5 | -0.8504558 | 0.0014324 | 0.0193327 |
| PDGFRB | -0.8468346 | 0.0061247 | 0.0487028 |
| BARD1 | -0.8456906 | 0.0005765 | 0.0123771 |
| BATF3 | -0.8365931 | 0.0000664 | 0.0046109 |
| WDR76 | -0.8175665 | 0.0036285 | 0.0351677 |
| VIM | -0.7957169 | 0.0049604 | 0.0425173 |
| CHEK2 | -0.7810276 | 0.0001973 | 0.0084018 |
| MCM6 | -0.7736048 | 0.0011457 | 0.0175845 |

| | | | |
|---|---|---|---|
| STC1 | -0.7599104 | 0.0033766 | 0.0337661 |
| CDC20 | -0.7396007 | 0.005103 | 0.0425456 |
| KDM3A | -0.7026551 | 0.0039056 | 0.0365197 |
| RPS2 | -0.7002605 | 0.0014112 | 0.0193327 |
| SMAD5 | -0.6950034 | 0.0002649 | 0.0087748 |
| TFRC | -0.6846632 | 0.0050325 | 0.0425456 |
| SETD1B | -0.6810885 | 0.0069586 | 0.053447 |
| SMC1A | -0.6091249 | 0.0007631 | 0.0145991 |
| CASP6 | -0.5827985 | 0.0018277 | 0.0227601 |
| BCL10 | -0.5449866 | 0.0003078 | 0.008909 |
| SETDB1 | -0.5359754 | 0.0006617 | 0.0133781 |
| TLR9 | -0.533844 | 0.0049496 | 0.0425173 |
| OGG1 | -0.5330204 | 0.0028084 | 0.0303858 |
| NRAS | -0.5315765 | 0.0046228 | 0.0415107 |
| RALA | -0.5254246 | 0.006969 | 0.053447 |
| RELA | -0.5111642 | 0.0010969 | 0.0174441 |
| TWF1 | -0.5012204 | 0.0030938 | 0.0324111 |
| CALR | -0.4996751 | 0.0063064 | 0.049847 |
| MLH1 | -0.4648316 | 0.0055606 | 0.0453086 |
| HUS1 | -0.4463035 | 0.0035493 | 0.0349632 |

Table S3

| ID | FC | P.Value | adj.P.Val |
|---|---|---|---|
| NEUG | 1.9910878 | 0.0002401 | 0.0108114 |
| TPPP | 1.865966 | 0.0000833 | 0.0098936 |
| CPLX2 | 1.7639059 | 0.0005697 | 0.0121069 |
| ADCY1 | 1.7261731 | 0.0038158 | 0.022895 |
| SYPH | 1.6756184 | 0.0004099 | 0.0114048 |
| VAMP1 | 1.6442547 | 0.0002789 | 0.011047 |
| STXB1 | 1.6407932 | 0.0004238 | 0.0114048 |
| SV2A | 1.6178553 | 0.0015215 | 0.0155249 |
| AT1B1 | 1.6094698 | 0.0003708 | 0.0114048 |
| SYUA | 1.6088477 | 0.0004448 | 0.0114605 |
| KCC2A | 1.6030719 | 0.0005416 | 0.0119152 |
| PACN1 | 1.6012639 | 0.0003068 | 0.0112458 |
| STX1B | 1.5739163 | 0.000382 | 0.0114048 |
| AT1A3 | 1.5631302 | 0.0005753 | 0.0121069 |
| SYUB | 1.5376025 | 0.0004112 | 0.0114048 |
| KCRU | 1.5126472 | 0.0003366 | 0.0112859 |
| AINX | 1.504588 | 0.0001452 | 0.0099188 |
| SYN1 | 1.5016716 | 0.0012144 | 0.0141972 |
| IGSF8 | 1.4993491 | 0.0002244 | 0.0104486 |
| SNAB | 1.4955866 | 0.0001221 | 0.0098936 |
| FABPH | 1.466362 | 0.000012 | 0.0098936 |
| CADM2 | 1.4599182 | 0.0018706 | 0.0160152 |
| DYN1 | 1.4586292 | 0.000277 | 0.011047 |
| VISL1 | 1.4574119 | 0.0001994 | 0.0100934 |
| BASP1 | 1.4477715 | 0.0017924 | 0.0157164 |
| PHF24 | 1.4449206 | 0.0003135 | 0.0112458 |
| SHPS1 | 1.4445422 | 0.0005599 | 0.0120631 |
| NFM | 1.4390092 | 0.0007876 | 0.0125466 |
| SYT1 | 1.4356262 | 0.0015509 | 0.0156055 |
| TBA8 | 1.4340395 | 0.0003829 | 0.0114048 |
| SNP25 | 1.4331029 | 0.0002426 | 0.0108114 |
| S7A14 | 1.4312225 | 0.0119457 | 0.0450475 |
| ATIF1 | 1.4283551 | 0.0001023 | 0.0098936 |
| VATG2 | 1.4277655 | 0.0001487 | 0.0099188 |
| AATC | 1.4255934 | 0.0000547 | 0.0098936 |
| RAB3A | 1.4234487 | 0.003017 | 0.0200511 |
| PSD3 | 1.4132075 | 0.0005622 | 0.0120631 |
| SH3G2 | 1.4106049 | 0.0001901 | 0.0100934 |
| HPCL4 | 1.4058014 | 0.0000725 | 0.0098936 |
| SYN2 | 1.4050499 | 0.0009736 | 0.0134835 |
| VGLU1 | 1.4020301 | 0.0024694 | 0.0179157 |
| SH3G3 | 1.3900765 | 0.0000456 | 0.0098936 |
| AMPH | 1.3838843 | 0.0004206 | 0.0114048 |
| ENOG | 1.3831027 | 0.0001588 | 0.0099746 |
| S12A5 | 1.3810853 | 0.0001627 | 0.0099806 |
| ICAM5 | 1.3757709 | 0.0026706 | 0.0185659 |

| | | | |
|---|---|---|---|
| CRYM | 1.3645831 | 0.0001087 | 0.0098936 |
| PLM | 1.3616939 | 0.0017105 | 0.0157054 |
| NCDN | 1.3579646 | 0.0003266 | 0.0112859 |
| SYUG | 1.3445051 | 0.0000771 | 0.0098936 |
| GNAO | 1.342923 | 0.0018328 | 0.015866 |
| KCD16 | 1.3418229 | 0.0003044 | 0.0112458 |
| DLGP3 | 1.3414422 | 0.0002612 | 0.0109942 |
| NEGR1 | 1.34013 | 0.0012883 | 0.0144765 |
| KCC2B | 1.3352225 | 0.0004108 | 0.0114048 |
| VPP1 | 1.3343233 | 0.0015533 | 0.0156055 |
| CALB2 | 1.33328 | 0.0001089 | 0.0098936 |
| CPLX1 | 1.3328027 | 0.0000567 | 0.0098936 |
| PHAR2 | 1.3263727 | 0.000547 | 0.0119477 |
| SYN3 | 1.325525 | 0.0019136 | 0.0161188 |
| HXK1 | 1.3243788 | 0.0006436 | 0.012285 |
| CCSAP | 1.3227356 | 0.0002875 | 0.0110797 |
| PKHA1 | 1.3179061 | 0.0002047 | 0.0100934 |
| NOE1 | 1.3133265 | 0.0010423 | 0.0136207 |
| TAU | 1.3132661 | 0.0001668 | 0.0099806 |
| VA0D1 | 1.31051 | 0.0026543 | 0.0185266 |
| DLG4 | 1.3022871 | 0.0004977 | 0.0118593 |
| ACBD7 | 1.3000286 | 0.0006622 | 0.0123201 |
| DEMA | 1.2977872 | 0.0001055 | 0.0098936 |
| RP3A | 1.2963419 | 0.0034955 | 0.0218931 |
| CPNE6 | 1.2939161 | 0.0001524 | 0.0099188 |
| SH3L2 | 1.2909198 | 0.000049 | 0.0098936 |
| PALM | 1.286484 | 0.0005244 | 0.0118593 |
| BIN1 | 1.2846437 | 0.0000451 | 0.0098936 |
| IQEC3 | 1.284021 | 0.0002571 | 0.0109942 |
| HCN1 | 1.282533 | 0.0001525 | 0.0099188 |
| CAMKV | 1.2820069 | 0.0013638 | 0.0147832 |
| GUAD | 1.2778342 | 0.0000516 | 0.0098936 |
| GPC5 | 1.2762349 | 0.0083182 | 0.0358365 |
| CN37 | 1.2740712 | 0.0083234 | 0.0358365 |
| TBB2A | 1.2728736 | 0.0011854 | 0.0140408 |
| NELL2 | 1.2672183 | 0.0003055 | 0.0112458 |
| KPCG | 1.2666421 | 0.0014887 | 0.0154658 |
| HPRT | 1.2656306 | 0.0000692 | 0.0098936 |
| MDHC | 1.2640906 | 0.0000708 | 0.0098936 |
| SV2B | 1.2624084 | 0.002101 | 0.0168018 |
| WASF1 | 1.2580948 | 0.0001461 | 0.0099188 |
| CX6B1 | 1.2547792 | 0.0007026 | 0.0123757 |
| LFG2 | 1.2544637 | 0.0024986 | 0.018019 |
| K0513 | 1.247611 | 0.0007676 | 0.0124986 |
| MAL2 | 1.2427525 | 0.0057203 | 0.0286236 |
| ZNT3 | 1.2427212 | 0.0016568 | 0.0156495 |
| PP2BA | 1.2416997 | 0.000559 | 0.0120631 |
| NPTX1 | 1.2397798 | 0.0013207 | 0.0146516 |

| | | | |
|---|---|---|---|
| HPCA | 1.2395492 | 0.0024074 | 0.017854 |
| SNG1 | 1.23508 | 0.001913 | 0.0161188 |
| KAD5 | 1.2292364 | 0.0000447 | 0.0098936 |
| NCKX2 | 1.2285355 | 0.0041234 | 0.0238581 |
| KNDC1 | 1.2280713 | 0.0007816 | 0.0125466 |
| AT2B3 | 1.2264827 | 0.0000127 | 0.0098936 |
| NCS1 | 1.2256648 | 0.0052227 | 0.0271769 |
| GNAZ | 1.2234158 | 0.0007118 | 0.0124393 |
| UCHL1 | 1.2187892 | 0.00008 | 0.0098936 |
| GDAP1 | 1.2143861 | 0.0017145 | 0.0157054 |
| SCG1 | 1.2097241 | 0.0074743 | 0.0337118 |
| MAP6 | 1.2094834 | 0.0000677 | 0.0098936 |
| PRRT3 | 1.2090952 | 0.0026092 | 0.0184094 |
| PHYIP | 1.2074881 | 0.0004148 | 0.0114048 |
| OPALI | 1.2050391 | 0.0124261 | 0.0460481 |
| NFL | 1.2013145 | 0.0019005 | 0.0160604 |
| STX1A | 1.1996954 | 0.0021449 | 0.0169702 |
| NAC2 | 1.1993013 | 0.0030444 | 0.0201074 |
| RAB3C | 1.1942917 | 0.0022183 | 0.017263 |
| RIMS1 | 1.1942147 | 0.0001924 | 0.0100934 |
| SCAM5 | 1.1830957 | 0.0001314 | 0.0098936 |
| SYGP1 | 1.1828939 | 0.0006796 | 0.0123592 |
| NDUS6 | 1.1764415 | 0.0001843 | 0.0100934 |
| RASL1 | 1.1747303 | 0.0000738 | 0.0098936 |
| AUXI | 1.1737522 | 0.0003181 | 0.0112458 |
| NMDZ1 | 1.172985 | 0.0003954 | 0.0114048 |
| PP2BB | 1.1721534 | 0.0004654 | 0.0115541 |
| AP180 | 1.1684826 | 0.0001222 | 0.0098936 |
| CEND | 1.1677884 | 0.0041142 | 0.0238391 |
| NPTN | 1.1657654 | 0.0019913 | 0.0164544 |
| MARE3 | 1.1656782 | 0.0004659 | 0.0115541 |
| AT2B2 | 1.1650386 | 0.0011836 | 0.0140408 |
| OPCM | 1.1634382 | 0.0007685 | 0.0124986 |
| NPTXR | 1.1610119 | 0.0042597 | 0.0243263 |
| SODC | 1.15707 | 0.0009713 | 0.0134835 |
| DCE2 | 1.1568135 | 0.0001081 | 0.0098936 |
| IQEC2 | 1.1567046 | 0.0002134 | 0.0103003 |
| VAMP3 | 1.1520715 | 0.0050712 | 0.0266857 |
| SYT12 | 1.150749 | 0.0010978 | 0.01388 |
| NSF | 1.1452993 | 0.000242 | 0.0108114 |
| NFH | 1.1452247 | 0.0006353 | 0.0122235 |
| GABR1 | 1.1440713 | 0.0005072 | 0.0118593 |
| TPRGL | 1.1435487 | 0.0001645 | 0.0099806 |
| TAGL3 | 1.140949 | 0.0009879 | 0.0134835 |
| F241B | 1.1363284 | 0.0021381 | 0.016968 |
| FBX2 | 1.1330777 | 0.000023 | 0.0098936 |
| PPR1B | 1.1324613 | 0.0006243 | 0.0121479 |
| NDRG4 | 1.1281526 | 0.0003351 | 0.0112859 |

| | | | |
|---|---|---|---|
| KAD1 | 1.1280958 | 0.0006963 | 0.0123592 |
| SCN2B | 1.1252747 | 0.001086 | 0.013861 |
| TBA4A | 1.1231333 | 0.002208 | 0.0172086 |
| CADM3 | 1.120614 | 0.0001085 | 0.0098936 |
| PRRT2 | 1.1165733 | 0.0027895 | 0.0191879 |
| KCC2G | 1.1162109 | 0.0003811 | 0.0114048 |
| CACB3 | 1.1162089 | 0.0003214 | 0.0112458 |
| GHC1 | 1.1147611 | 0.0013375 | 0.014682 |
| AT8A1 | 1.1147483 | 0.0010771 | 0.0138149 |
| MA7D2 | 1.1137944 | 0.0001465 | 0.0099188 |
| QCR6 | 1.1107189 | 0.0043561 | 0.0246016 |
| OGDHL | 1.1099928 | 0.0008646 | 0.0128902 |
| SEPT5 | 1.1050655 | 0.0023975 | 0.017854 |
| SNG3 | 1.1045241 | 0.001576 | 0.0156366 |
| SHSA7 | 1.1018989 | 0.0026208 | 0.0184567 |
| ATP5E | 1.1015814 | 0.0002914 | 0.0110894 |
| CAPS1 | 1.1004071 | 0.0004036 | 0.0114048 |
| PP14C | 1.0991067 | 0.0030512 | 0.0201126 |
| CANB1 | 1.0989957 | 0.0001539 | 0.0099188 |
| VAMP2 | 1.0983778 | 0.0004828 | 0.0117635 |
| CNTN1 | 1.0954152 | 0.0022928 | 0.0174581 |
| SCG2 | 1.0948793 | 0.0011837 | 0.0140408 |
| BSN | 1.0933706 | 0.0012245 | 0.0142255 |
| SNTA1 | 1.0866236 | 0.0005916 | 0.0121069 |
| EAA2 | 1.0864191 | 0.002951 | 0.0198021 |
| RYR2 | 1.0857683 | 0.0013 | 0.0145148 |
| GBRL1 | 1.0849544 | 0.0016658 | 0.0157032 |
| MOG | 1.0843681 | 0.0048671 | 0.0261351 |
| CISD1 | 1.0832125 | 0.0004462 | 0.0114605 |
| TIAM2 | 1.0826468 | 0.0011696 | 0.0140408 |
| DLGP1 | 1.0814688 | 0.0006036 | 0.0121069 |
| L1CAM | 1.0806637 | 0.0013518 | 0.0147766 |
| OXR1 | 1.0795884 | 0.000409 | 0.0114048 |
| CA2D1 | 1.0793069 | 0.001552 | 0.0156055 |
| LY6H | 1.0787796 | 0.0061738 | 0.0300282 |
| HS12A | 1.0781499 | 0.000315 | 0.0112458 |
| ST32C | 1.0772693 | 0.0008261 | 0.0127211 |
| CCKN | 1.0770358 | 0.0018341 | 0.015866 |
| GLNA | 1.0699005 | 0.0106429 | 0.0420076 |
| PSD1 | 1.0669744 | 0.0064937 | 0.0309489 |
| AT1A2 | 1.0649034 | 0.0016717 | 0.0157032 |
| CLCB | 1.0648611 | 0.0008838 | 0.012953 |
| LIPA3 | 1.0642685 | 0.0001009 | 0.0098936 |
| ANK2 | 1.0637407 | 0.0004237 | 0.0114048 |
| VATA | 1.0570862 | 0.000503 | 0.0118593 |
| E41L3 | 1.057014 | 0.0003107 | 0.0112458 |
| KKCC1 | 1.0540658 | 0.0002075 | 0.0101126 |
| IDH3A | 1.0516523 | 0.0002605 | 0.0109942 |

| | | | |
|---|---|---|---|
| SHLB2 | 1.0515757 | 0.0001359 | 0.0099188 |
| MIC25 | 1.0506103 | 0.0004017 | 0.0114048 |
| TPD53 | 1.0505138 | 0.0001385 | 0.0099188 |
| MAP1A | 1.0495197 | 0.0001884 | 0.0100934 |
| SH2D5 | 1.046257 | 0.0009836 | 0.0134835 |
| E41L1 | 1.0454923 | 0.0012066 | 0.0141376 |
| PLCB1 | 1.0451957 | 0.0006889 | 0.0123592 |
| SYNJ1 | 1.0444077 | 0.0000705 | 0.0098936 |
| COX5A | 1.0422743 | 0.0018358 | 0.015866 |
| DNJC5 | 1.0399055 | 0.0011728 | 0.0140408 |
| ADT1 | 1.0391822 | 0.0017657 | 0.0157164 |
| NDUA2 | 1.0359221 | 0.0002468 | 0.0109053 |
| PCLO | 1.0354686 | 0.0004254 | 0.0114048 |
| VATH | 1.0345561 | 0.0003789 | 0.0114048 |
| NPAL3 | 1.0344735 | 0.0017746 | 0.0157164 |
| K154L | 1.0338954 | 0.0006736 | 0.0123592 |
| DC1I1 | 1.0333413 | 0.0000733 | 0.0098936 |
| PHAR1 | 1.0308945 | 0.0000938 | 0.0098936 |
| AT2B1 | 1.0260159 | 0.0064319 | 0.0307291 |
| X7B2 | 1.0236083 | 0.0000985 | 0.0098936 |
| GBRA1 | 1.0230593 | 0.0035023 | 0.0218931 |
| NFASC | 1.022682 | 0.009886 | 0.0401035 |
| GBG7 | 1.0205494 | 0.0005792 | 0.0121069 |
| PPR1A | 1.0202505 | 0.0036912 | 0.0226423 |
| AATM | 1.0199898 | 0.0014488 | 0.015292 |
| GNAI1 | 1.0183572 | 0.0033799 | 0.021439 |
| MLF2 | 1.0171966 | 0.0029889 | 0.0199028 |
| CARL2 | 1.0143719 | 0.0004372 | 0.0114564 |
| ACON | 1.013251 | 0.0007782 | 0.0125466 |
| VATE1 | 1.0119358 | 0.0005851 | 0.0121069 |
| VATB2 | 1.0081705 | 0.0004448 | 0.0114605 |
| SHAN3 | 1.0059808 | 0.0022477 | 0.0172874 |
| PEBP1 | 1.0027468 | 0.0016383 | 0.0156495 |
| PFKAP | 1.000489 | 0.0000607 | 0.0098936 |
| CC177 | 1.0004527 | 0.0032049 | 0.0206555 |
| ANS1B | 0.9976989 | 0.00109 | 0.013861 |
| TBB4A | 0.9964404 | 0.0003335 | 0.0112859 |
| GLSK | 0.9961289 | 0.0004167 | 0.0114048 |
| NECA1 | 0.99568 | 0.0021552 | 0.0170261 |
| RHOB | 0.9954996 | 0.0024223 | 0.0178895 |
| PYGM | 0.9950716 | 0.000832 | 0.0127211 |
| KPCE | 0.9943166 | 0.0002224 | 0.0104472 |
| SPTN1 | 0.9939631 | 0.0005726 | 0.0121069 |
| SEPT3 | 0.9892539 | 0.0012232 | 0.0142255 |
| K1107 | 0.986688 | 0.0007982 | 0.0126095 |
| PTPRN | 0.9865217 | 0.000914 | 0.0130695 |
| PROF2 | 0.9842516 | 0.0002016 | 0.0100934 |
| SKP1 | 0.9841777 | 0.0004647 | 0.0115541 |

| | | | |
|---|---|---|---|
| STXB6 | 0.9821131 | 0.0011753 | 0.0140408 |
| CMGA | 0.9811186 | 0.0006334 | 0.0122235 |
| NMDE1 | 0.9807583 | 0.0053274 | 0.0275022 |
| ATP5I | 0.9799272 | 0.0002052 | 0.0100934 |
| AJM1 | 0.9795841 | 0.0017276 | 0.0157054 |
| VATC1 | 0.9784576 | 0.0002662 | 0.011047 |
| X1433Z | 0.9776854 | 0.0007693 | 0.0124986 |
| THAP4 | 0.9776824 | 0.0017743 | 0.0157164 |
| KCAB2 | 0.9768165 | 0.0042325 | 0.0242772 |
| RGS7 | 0.9757091 | 0.0007557 | 0.0124986 |
| S6A17 | 0.9744072 | 0.0004505 | 0.0115146 |
| K121L | 0.9743675 | 0.0006479 | 0.012285 |
| KCRB | 0.9691148 | 0.0089934 | 0.037694 |
| ACTN2 | 0.9649222 | 0.0005213 | 0.0118593 |
| SCOT1 | 0.9642964 | 0.0005201 | 0.0118593 |
| BACH | 0.9600016 | 0.0009091 | 0.0130695 |
| NDUS4 | 0.9597339 | 0.0001095 | 0.0098936 |
| SNPH | 0.9594162 | 0.0010305 | 0.0136032 |
| ADDB | 0.9544914 | 0.0003398 | 0.0112859 |
| DHPR | 0.9507156 | 0.00022 | 0.0104472 |
| F126B | 0.9497206 | 0.000106 | 0.0098936 |
| RASH | 0.9482627 | 0.0023419 | 0.0176965 |
| FRIH | 0.9476079 | 0.000156 | 0.0099188 |
| PDE1B | 0.9473855 | 0.0012419 | 0.0142673 |
| FRIL | 0.9473594 | 0.0004853 | 0.0117685 |
| SCN2A | 0.9454656 | 0.0005188 | 0.0118593 |
| RPGP2 | 0.9450023 | 0.0011199 | 0.0139693 |
| NDUA7 | 0.9427693 | 0.0015894 | 0.0156495 |
| IDH3B | 0.9410047 | 0.0003869 | 0.0114048 |
| ATPA | 0.9382704 | 0.0018876 | 0.0160604 |
| NDUS8 | 0.937151 | 0.0021852 | 0.0171048 |
| ATP5H | 0.9367078 | 0.0010784 | 0.0138149 |
| HPLN2 | 0.9356419 | 0.0047743 | 0.0258495 |
| KPCB | 0.9332323 | 0.0019854 | 0.0164317 |
| C170B | 0.932042 | 0.0001128 | 0.0098936 |
| DPP10 | 0.9315919 | 0.0008777 | 0.0129281 |
| MLP3A | 0.9310497 | 0.0001421 | 0.0099188 |
| SRCN1 | 0.9289365 | 0.0000838 | 0.0098936 |
| DYN3 | 0.9254702 | 0.003416 | 0.0216152 |
| SUCB1 | 0.9234256 | 0.0007882 | 0.0125466 |
| PCSK1 | 0.9224701 | 0.002084 | 0.0167685 |
| TPD52 | 0.9222163 | 0.0000372 | 0.0098936 |
| HOME1 | 0.9210796 | 0.0009124 | 0.0130695 |
| GNB5 | 0.9203327 | 0.0004078 | 0.0114048 |
| SGIP1 | 0.9197746 | 0.0004322 | 0.0114048 |
| RIMS2 | 0.9179168 | 0.0032827 | 0.021027 |
| IDH3G | 0.9176835 | 0.000588 | 0.0121069 |
| GBB1 | 0.9157471 | 0.0024141 | 0.017854 |

| | | | |
|---|---|---|---|
| DPYL2 | 0.9147115 | 0.0007385 | 0.0124668 |
| SKT | 0.9133162 | 0.0000893 | 0.0098936 |
| VATD | 0.9120748 | 0.0005915 | 0.0121069 |
| GRM2 | 0.9111606 | 0.0030978 | 0.0202842 |
| FBX41 | 0.9066178 | 0.0026262 | 0.0184567 |
| NECP1 | 0.904761 | 0.0000431 | 0.0098936 |
| PLXA4 | 0.9037088 | 0.0016247 | 0.0156495 |
| CSRP1 | 0.900612 | 0.0024422 | 0.0179029 |
| NDUV2 | 0.8992664 | 0.0017494 | 0.0157054 |
| X1433G | 0.8962301 | 0.0037517 | 0.0227459 |
| SYNPO | 0.8956577 | 0.000716 | 0.0124393 |
| CYC | 0.8948635 | 0.0106925 | 0.0421076 |
| PNM8B | 0.8943086 | 0.007079 | 0.0327508 |
| VIAAT | 0.8924725 | 0.0043998 | 0.0246664 |
| ATPG | 0.8921995 | 0.0015709 | 0.0156366 |
| LZTS3 | 0.8914161 | 0.0025621 | 0.0182249 |
| AUHM | 0.8905616 | 0.0000846 | 0.0098936 |
| KCTD8 | 0.8904881 | 0.0121378 | 0.0455627 |
| ABLM2 | 0.890393 | 0.0015389 | 0.0155807 |
| LGI1 | 0.8891263 | 0.0006663 | 0.0123201 |
| PRIO | 0.8888888 | 0.0014879 | 0.0154658 |
| MP2K1 | 0.8877517 | 0.000832 | 0.0127211 |
| TMOD2 | 0.8877267 | 0.0009626 | 0.0134835 |
| DLG2 | 0.8853913 | 0.0001191 | 0.0098936 |
| ATPD | 0.8845332 | 0.0000924 | 0.0098936 |
| CPEB3 | 0.8844008 | 0.0003495 | 0.0113186 |
| ANK3 | 0.8839516 | 0.0003824 | 0.0114048 |
| GGA3 | 0.8809217 | 0.0049901 | 0.0265764 |
| GRM5 | 0.8804388 | 0.0008659 | 0.0128902 |
| NDUB5 | 0.8794745 | 0.0118139 | 0.0447984 |
| EF1A2 | 0.879449 | 0.0050771 | 0.0266857 |
| PURA | 0.8786608 | 0.0005271 | 0.0118593 |
| CA2D3 | 0.8784994 | 0.0006444 | 0.012285 |
| ADA23 | 0.8778992 | 0.0026336 | 0.0184752 |
| KCC4 | 0.8778297 | 0.0048742 | 0.0261398 |
| F171B | 0.8760294 | 0.0004659 | 0.0115541 |
| AT5F1 | 0.8750042 | 0.00082 | 0.0127211 |
| NDUA4 | 0.8736579 | 0.0044324 | 0.0247035 |
| SPTB2 | 0.8729893 | 0.0016326 | 0.0156495 |
| SEPT4 | 0.8728589 | 0.000528 | 0.0118593 |
| SBP1 | 0.8727583 | 0.0000376 | 0.0098936 |
| CNRP1 | 0.8694264 | 0.0035061 | 0.0218931 |
| VATF | 0.8669498 | 0.0002042 | 0.0100934 |
| BRSK2 | 0.8666323 | 0.0005995 | 0.0121069 |
| MP2K4 | 0.8657671 | 0.0006907 | 0.0123592 |
| C2C2L | 0.865677 | 0.0007539 | 0.0124986 |
| LEGL | 0.8654399 | 0.0008462 | 0.0128304 |
| NRX3A | 0.8649989 | 0.0039484 | 0.0233601 |

| | | | |
|---|---|---|---|
| RPGF4 | 0.8648829 | 0.007567 | 0.033945 |
| C2C4C | 0.864677 | 0.0069387 | 0.03244 |
| CC50A | 0.8640182 | 0.0128943 | 0.0470225 |
| ST4A1 | 0.8624919 | 0.0004611 | 0.0115541 |
| AAK1 | 0.859422 | 0.0001779 | 0.0100934 |
| PIN1 | 0.8594031 | 0.0003862 | 0.0114048 |
| ECHM | 0.8566806 | 0.0003185 | 0.0112458 |
| ATPB | 0.8560125 | 0.011117 | 0.043198 |
| PKHA6 | 0.8556373 | 0.0000483 | 0.0098936 |
| NDUA5 | 0.8550192 | 0.0036884 | 0.0226423 |
| GABR2 | 0.8518207 | 0.0028553 | 0.0194866 |
| KCC1D | 0.8515268 | 0.003068 | 0.0201697 |
| PLPP | 0.8511182 | 0.000502 | 0.0118593 |
| ARRB1 | 0.8504614 | 0.0000541 | 0.0098936 |
| QCR8 | 0.8499876 | 0.001917 | 0.0161216 |
| X1433F | 0.8490225 | 0.0004187 | 0.0114048 |
| JAM3 | 0.8476437 | 0.0022964 | 0.0174581 |
| IQEC1 | 0.8461879 | 0.0006546 | 0.0123201 |
| BAIP2 | 0.8451677 | 0.0006237 | 0.0121479 |
| TPM1 | 0.8430349 | 0.0007424 | 0.0124668 |
| BRSK1 | 0.8347355 | 0.0002752 | 0.011047 |
| RTN3 | 0.8345878 | 0.0008053 | 0.0126095 |
| CALM1.CALM | 0.8338011 | 0.0029046 | 0.0196168 |
| PHAR3 | 0.832923 | 0.0015805 | 0.0156367 |
| QCR7 | 0.8316366 | 0.0018052 | 0.0157397 |
| ADCY5 | 0.831009 | 0.0075944 | 0.033989 |
| LYRM9 | 0.8310042 | 0.0016472 | 0.0156495 |
| RAP2A | 0.8306041 | 0.0019685 | 0.0163878 |
| PTN5 | 0.8298735 | 0.0016276 | 0.0156495 |
| NDUA8 | 0.8269274 | 0.0050182 | 0.0265873 |
| NAC1 | 0.8214034 | 0.001748 | 0.0157054 |
| PDE2A | 0.8210025 | 0.0045959 | 0.0253041 |
| HCN2 | 0.820476 | 0.0007463 | 0.0124726 |
| NDUV1 | 0.8201436 | 0.0006051 | 0.0121069 |
| PDE1A | 0.8194295 | 0.0019726 | 0.0163878 |
| GRM3 | 0.8183871 | 0.0003831 | 0.0114048 |
| NMDE2 | 0.8182691 | 0.002716 | 0.0187564 |
| SYT7 | 0.81822 | 0.0047138 | 0.0256774 |
| LIPA2 | 0.8178606 | 0.0005334 | 0.0118593 |
| MARE2 | 0.8159377 | 0.0014323 | 0.0151787 |
| SPTN2 | 0.8149685 | 0.0010059 | 0.0135177 |
| AVL9 | 0.8146313 | 0.0016986 | 0.0157054 |
| PLCX3 | 0.8132032 | 0.0072608 | 0.0331797 |
| ERC2 | 0.8127836 | 0.0014123 | 0.0149974 |
| ODPA | 0.8123569 | 0.0017385 | 0.0157054 |
| X68MP | 0.8122343 | 0.0017269 | 0.0157054 |
| MINK1 | 0.8113488 | 0.0008465 | 0.0128304 |
| PAK1 | 0.8110492 | 0.0013609 | 0.0147828 |

| | | | |
|---|---|---|---|
| CAPS2 | 0.8084522 | 0.0058955 | 0.0291413 |
| LSAMP | 0.8078501 | 0.0083499 | 0.0358689 |
| NCAM1 | 0.8075232 | 0.005301 | 0.0274474 |
| AL1A1 | 0.8058069 | 0.0003465 | 0.0113186 |
| TINAL | 0.8045947 | 0.0002735 | 0.011047 |
| DNJA4 | 0.8031382 | 0.0001734 | 0.010048 |
| AP2A1 | 0.8008454 | 0.0042851 | 0.0243651 |
| SEPT6 | 0.8001834 | 0.0016966 | 0.0157054 |
| GRIN1 | 0.8000142 | 0.0055298 | 0.0279927 |
| CNTP1 | 0.7985975 | 0.001152 | 0.0140023 |
| CYFP2 | 0.7977529 | 0.0011329 | 0.0140023 |
| MDHM | 0.7958916 | 0.0053468 | 0.0275305 |
| MAP7 | 0.7951335 | 0.0115342 | 0.0443507 |
| SCG3 | 0.7945961 | 0.0059707 | 0.0293969 |
| PGFRB | 0.7944989 | 0.0117867 | 0.0447929 |
| ODPB | 0.7903269 | 0.001455 | 0.0153263 |
| CNTP2 | 0.7885857 | 0.0017944 | 0.0157164 |
| NAPEP | 0.7873436 | 0.0008399 | 0.0128055 |
| DLG3 | 0.7868252 | 0.0000346 | 0.0098936 |
| AN13D | 0.7865506 | 0.001869 | 0.0160152 |
| MK10 | 0.7845446 | 0.0001888 | 0.0100934 |
| NDUS1 | 0.7832253 | 0.0043243 | 0.0245075 |
| AP2M1 | 0.7830343 | 0.0016557 | 0.0156495 |
| EHD3 | 0.7825019 | 0.0023855 | 0.0178452 |
| DCD | 0.7821681 | 0.0005254 | 0.0118593 |
| MAST3 | 0.7814261 | 0.0082367 | 0.0356398 |
| TBB6 | 0.7809021 | 0.0007403 | 0.0124668 |
| ANLN | 0.7805998 | 0.0060085 | 0.0294432 |
| UCRI | 0.7804654 | 0.0020358 | 0.0167092 |
| ADCY2 | 0.7783977 | 0.0035933 | 0.0221986 |
| NTRK3 | 0.7779166 | 0.0029158 | 0.0196168 |
| I5P1 | 0.7776177 | 0.0020247 | 0.0167035 |
| ACPM | 0.7773144 | 0.0010898 | 0.013861 |
| KAP3 | 0.7773122 | 0.002411 | 0.017854 |
| ATPO | 0.7755178 | 0.004535 | 0.0251012 |
| NDEL1 | 0.7729055 | 0.0070989 | 0.0328018 |
| DNJB4 | 0.7726336 | 0.0017527 | 0.0157054 |
| COX7R | 0.7717336 | 0.0074519 | 0.0337017 |
| CAMP3 | 0.7702495 | 0.000923 | 0.0130777 |
| APLP1 | 0.7689718 | 0.0014107 | 0.0149974 |
| ADDA | 0.7679066 | 0.001377 | 0.0148345 |
| VAS1 | 0.765868 | 0.0122617 | 0.0456661 |
| DDAH1 | 0.7645446 | 0.0018575 | 0.0159816 |
| PKHA7 | 0.7642087 | 0.0011508 | 0.0140023 |
| X1433S | 0.7641752 | 0.0007 | 0.0123714 |
| CB39L | 0.7641598 | 0.000983 | 0.0134835 |
| TTBK1 | 0.7634821 | 0.0012793 | 0.0144581 |
| MROH1 | 0.7633873 | 0.0000297 | 0.0098936 |

| | | | |
|---|---|---|---|
| GFRA2 | 0.7633299 | 0.0023736 | 0.0178071 |
| WDR37 | 0.7629095 | 0.0003539 | 0.0113391 |
| SGTB | 0.7622302 | 0.001109 | 0.0139146 |
| MAON | 0.7605872 | 0.0007405 | 0.0124668 |
| DYL2 | 0.7597457 | 0.0044095 | 0.0246664 |
| CYBR1 | 0.7580854 | 0.000442 | 0.0114605 |
| KBTBB | 0.7539201 | 0.0006244 | 0.0121479 |
| CCD92 | 0.7534294 | 0.0010292 | 0.0136032 |
| NCEH1 | 0.7512819 | 0.0058754 | 0.0290927 |
| ODPX | 0.7510477 | 0.0012507 | 0.0143009 |
| DNM1L | 0.7498608 | 0.0020416 | 0.0167092 |
| CLH1 | 0.7495862 | 0.008079 | 0.0352061 |
| MA6D1 | 0.748518 | 0.0025085 | 0.018049 |
| CTRO | 0.7484012 | 0.0010782 | 0.0138149 |
| DLGP2 | 0.7483962 | 0.013731 | 0.049205 |
| PPM1H | 0.7481984 | 0.0023816 | 0.0178413 |
| RGRF2 | 0.7478706 | 0.0133318 | 0.0481052 |
| NOS1 | 0.7463978 | 0.0002774 | 0.011047 |
| WDR7 | 0.7453775 | 0.0002048 | 0.0100934 |
| PGAM1 | 0.7437242 | 0.001225 | 0.0142255 |
| LGI3 | 0.7433226 | 0.0051214 | 0.0268105 |
| MYO5A | 0.7430524 | 0.0039134 | 0.0232512 |
| RGS6 | 0.7430414 | 0.0054783 | 0.0278937 |
| AP2B1 | 0.7420792 | 0.0026469 | 0.0184999 |
| DOCK3 | 0.7417034 | 0.0013478 | 0.0147636 |
| IP3KA | 0.740626 | 0.0012923 | 0.01449 |
| RT36 | 0.7396869 | 0.0008869 | 0.012953 |
| COX5B | 0.7391766 | 0.0016543 | 0.0156495 |
| FSD1L | 0.7374436 | 0.0002003 | 0.0100934 |
| CCDC6 | 0.7369532 | 0.0001822 | 0.0100934 |
| ATP5L | 0.7358529 | 0.0060515 | 0.029571 |
| PI42B | 0.7354673 | 0.0017591 | 0.0157054 |
| STXB5 | 0.734648 | 0.0026266 | 0.0184567 |
| SHAN1 | 0.7328589 | 0.0056353 | 0.028307 |
| SCAI | 0.731479 | 0.0012578 | 0.0143507 |
| SNAG | 0.7303737 | 0.0013329 | 0.0146615 |
| NHRF1 | 0.7294927 | 0.0092091 | 0.0381081 |
| MBLC2 | 0.7290297 | 0.0037091 | 0.022649 |
| PPM1E | 0.7267657 | 0.0066365 | 0.0314601 |
| MAOX | 0.726341 | 0.000615 | 0.0121472 |
| QCR1 | 0.7246761 | 0.0080986 | 0.0352176 |
| NDUAA | 0.7226385 | 0.0066372 | 0.0314601 |
| CSKI1 | 0.7219489 | 0.0065975 | 0.0313575 |
| GAS7 | 0.7210261 | 0.0008697 | 0.0128902 |
| AKAP5 | 0.7208438 | 0.0024777 | 0.017918 |
| DLG1 | 0.7190725 | 0.0041149 | 0.0238391 |
| NEBL | 0.7188816 | 0.0007377 | 0.0124668 |
| NTRI | 0.7187309 | 0.0037695 | 0.0227481 |

| | | | |
|---|---|---|---|
| FAAH1 | 0.7178684 | 0.0012423 | 0.0142673 |
| FAK2 | 0.7171088 | 0.0047706 | 0.0258495 |
| COA6 | 0.7166842 | 0.0042727 | 0.0243435 |
| EPN1 | 0.7163927 | 0.0004168 | 0.0114048 |
| TRIM2 | 0.714453 | 0.0002615 | 0.0109942 |
| UBP31 | 0.7138594 | 0.0049303 | 0.0263658 |
| AT1B2 | 0.7115592 | 0.0058269 | 0.0289621 |
| CISD3 | 0.7110745 | 0.0017572 | 0.0157054 |
| MARK4 | 0.7106553 | 0.0039482 | 0.0233601 |
| MYCT | 0.7103782 | 0.0025586 | 0.0182249 |
| ATLA1 | 0.7060504 | 0.0022872 | 0.0174581 |
| RPGF2 | 0.7054864 | 0.0009108 | 0.0130695 |
| NDRG2 | 0.7034972 | 0.0022579 | 0.0172874 |
| RPGP1 | 0.7032554 | 0.0020903 | 0.0167933 |
| NCOA7 | 0.7022432 | 0.0020707 | 0.0167132 |
| ODP2 | 0.7019799 | 0.0015285 | 0.0155489 |
| CAD13 | 0.7019541 | 0.007199 | 0.0330128 |
| ACYP2 | 0.6990266 | 0.0066724 | 0.0315596 |
| PLXA2 | 0.6988531 | 0.0018705 | 0.0160152 |
| CNKR2 | 0.6973996 | 0.0044091 | 0.0246664 |
| ICA69 | 0.695424 | 0.0007436 | 0.0124668 |
| NDUS3 | 0.6945842 | 0.0051505 | 0.0268814 |
| OPA1 | 0.6941916 | 0.0051988 | 0.0270797 |
| X1433B | 0.6939682 | 0.0042255 | 0.0242643 |
| LYSM2 | 0.6916813 | 0.0009847 | 0.0134835 |
| MMPOS | 0.6907993 | 0.0027434 | 0.0189207 |
| OMGP | 0.6882314 | 0.0133027 | 0.0480666 |
| DMWD | 0.6879564 | 0.0000882 | 0.0098936 |
| HPCL1 | 0.6868601 | 0.0074096 | 0.0336092 |
| NDUC2 | 0.6857262 | 0.0058739 | 0.0290927 |
| RUN3A | 0.6857102 | 0.0112592 | 0.0436471 |
| CAD20 | 0.6844481 | 0.0086896 | 0.036881 |
| CYGB | 0.6822874 | 0.008927 | 0.0375103 |
| ASSY | 0.6819123 | 0.0002868 | 0.0110797 |
| LHPP | 0.6815056 | 0.0028019 | 0.0192228 |
| TM1L2 | 0.6811416 | 0.0010043 | 0.0135177 |
| NDUB7 | 0.6807688 | 0.0114209 | 0.0440775 |
| RGS17 | 0.6796774 | 0.0116567 | 0.0445298 |
| NDUB3 | 0.6791358 | 0.0054441 | 0.0278016 |
| AGAP2 | 0.6790355 | 0.0040402 | 0.0236107 |
| BORG4 | 0.6782015 | 0.0121611 | 0.0455829 |
| FN3K | 0.6770442 | 0.0016336 | 0.0156495 |
| EAA1 | 0.6769665 | 0.0132667 | 0.04797 |
| TB22A | 0.6757114 | 0.0060981 | 0.0297434 |
| AP2A2 | 0.6755119 | 0.0042959 | 0.0243996 |
| TTPAL | 0.6748219 | 0.0070499 | 0.032703 |
| S4A4 | 0.6747559 | 0.0017502 | 0.0157054 |
| DPP6 | 0.6728426 | 0.0044506 | 0.0247393 |

| | | | |
|---|---|---|---|
| HS74L | 0.672315 | 0.0004776 | 0.0116909 |
| IGS21 | 0.672097 | 0.0072947 | 0.0332762 |
| MTURN | 0.6693952 | 0.0008726 | 0.0128902 |
| CLCA | 0.6684068 | 0.0023333 | 0.017657 |
| LIGO1 | 0.6646797 | 0.0010539 | 0.0137079 |
| PIMT | 0.6642739 | 0.0021669 | 0.0170918 |
| ARL15 | 0.6642601 | 0.0062393 | 0.030097 |
| CLVS2 | 0.6637536 | 0.00107 | 0.0138149 |
| TBB4B | 0.6634938 | 0.0040642 | 0.0237033 |
| PRDX3 | 0.6629065 | 0.0009121 | 0.0130695 |
| S4A10 | 0.6628022 | 0.0101866 | 0.0408246 |
| COX7C | 0.6614354 | 0.0070256 | 0.0326702 |
| ARP5L | 0.6586558 | 0.0023505 | 0.0177362 |
| GSTM3 | 0.6544139 | 0.0052398 | 0.0272092 |
| LMO7 | 0.6525784 | 0.002459 | 0.0179029 |
| DLDH | 0.6511547 | 0.0015707 | 0.0156366 |
| ADA22 | 0.6499122 | 0.0030207 | 0.0200511 |
| CA2D2 | 0.64836 | 0.0045697 | 0.0252401 |
| HEM4 | 0.6482688 | 0.003729 | 0.0226985 |
| ATPK | 0.6481379 | 0.0062478 | 0.0301075 |
| GD1L1 | 0.6458892 | 0.0109118 | 0.0426494 |
| KKCC2 | 0.6454199 | 0.0080824 | 0.0352061 |
| WNK2 | 0.6435491 | 0.0104871 | 0.0415909 |
| COX7B | 0.6434133 | 0.0116641 | 0.0445298 |
| NB5R2 | 0.6430535 | 0.0020566 | 0.0167092 |
| RTN4 | 0.6429617 | 0.0038604 | 0.0230826 |
| PLCL1 | 0.6429093 | 0.0011389 | 0.0140023 |
| ISCU | 0.6426847 | 0.0007523 | 0.0124986 |
| WDR47 | 0.6425781 | 0.0013309 | 0.0146615 |
| MTMR5 | 0.6425633 | 0.001377 | 0.0148345 |
| CY1 | 0.641411 | 0.0087998 | 0.0372723 |
| SEM4D | 0.6412109 | 0.0084893 | 0.0362812 |
| BDH | 0.639983 | 0.001314 | 0.0146083 |
| NEC1 | 0.6398604 | 0.0056918 | 0.0285358 |
| UN13A | 0.6395861 | 0.0015178 | 0.0155249 |
| WIPF3 | 0.6377439 | 0.0098385 | 0.0400453 |
| EDIL3 | 0.6368136 | 0.0003037 | 0.0112458 |
| PICK1 | 0.6359171 | 0.0013287 | 0.0146615 |
| CAAP1 | 0.6355443 | 0.0018967 | 0.0160604 |
| PGBD5 | 0.6339782 | 0.0019518 | 0.0163353 |
| ZNRF2 | 0.6333122 | 0.0069965 | 0.0325711 |
| ABHDA | 0.6326479 | 0.0001555 | 0.0099188 |
| CADM1 | 0.6325163 | 0.0040056 | 0.0235456 |
| ACSL6 | 0.6315692 | 0.0008309 | 0.0127211 |
| ALDH2 | 0.6312318 | 0.0028621 | 0.0195072 |
| ATP5J | 0.6295317 | 0.007183 | 0.0329686 |
| TBA1A | 0.6292821 | 0.0080976 | 0.0352176 |
| SSDH | 0.6283897 | 0.0057647 | 0.028763 |

| | | | |
|---|---|---|---|
| PEX5R | 0.6283372 | 0.0038795 | 0.0231703 |
| LANC1 | 0.6283142 | 0.0046348 | 0.0254376 |
| NGEF | 0.6274679 | 0.0021797 | 0.0170918 |
| NDUBA | 0.6271725 | 0.0100998 | 0.0405704 |
| CPNE4 | 0.6269153 | 0.01094 | 0.0427276 |
| CYLD | 0.6263342 | 0.000376 | 0.0114048 |
| RAB4B | 0.6262333 | 0.0030466 | 0.0201074 |
| KAP1 | 0.6259607 | 0.0054835 | 0.0278937 |
| DAAM1 | 0.6255962 | 0.0053487 | 0.0275305 |
| CAP2 | 0.6253311 | 0.0022563 | 0.0172874 |
| DNJB2 | 0.6241647 | 0.0024293 | 0.0179029 |
| NDUS7 | 0.6236719 | 0.0076626 | 0.0341369 |
| STMN2 | 0.6207153 | 0.0037835 | 0.0228057 |
| GP158 | 0.6202006 | 0.0114666 | 0.044221 |
| ZC21A | 0.620094 | 0.0009179 | 0.0130757 |
| NDUAD | 0.6197471 | 0.0061675 | 0.0300254 |
| GSK3A | 0.6188947 | 0.0068269 | 0.0320971 |
| CRTC1 | 0.6186175 | 0.0015298 | 0.0155489 |
| WFS1 | 0.6173257 | 0.000109 | 0.0098936 |
| PTGDS | 0.6163381 | 0.008231 | 0.0356398 |
| ODO1 | 0.6143346 | 0.0076669 | 0.0341369 |
| EPHA4 | 0.6142914 | 0.0022575 | 0.0172874 |
| DTD1 | 0.6138345 | 0.0014099 | 0.0149974 |
| PAK3 | 0.6134382 | 0.0034361 | 0.0217162 |
| QCR2 | 0.6091872 | 0.0063658 | 0.0305068 |
| NED4L | 0.6079002 | 0.0025353 | 0.0181581 |
| TIM29 | 0.6073602 | 0.0021159 | 0.0168688 |
| PALM2 | 0.6064661 | 0.0082811 | 0.0357136 |
| DDHD2 | 0.6060696 | 0.0014918 | 0.0154658 |
| NDUA3 | 0.6045732 | 0.0094301 | 0.0388174 |
| NUMBL | 0.6024108 | 0.0012321 | 0.0142673 |
| PI51C | 0.5971502 | 0.0024064 | 0.017854 |
| MIC26 | 0.5968886 | 0.0037496 | 0.0227459 |
| ISCA1 | 0.5967006 | 0.0028908 | 0.0196073 |
| DJC30 | 0.5961431 | 0.0026674 | 0.0185659 |
| DCE1 | 0.5949794 | 0.0017297 | 0.0157054 |
| SCAM1 | 0.5943359 | 0.0048415 | 0.0260972 |
| OPTN | 0.591752 | 0.0006747 | 0.0123592 |
| GDS1 | 0.5885889 | 0.0012811 | 0.0144581 |
| NDUAC | 0.5866502 | 0.0129102 | 0.0470396 |
| NDUA6 | 0.5866475 | 0.0041086 | 0.0238391 |
| PCDH9 | 0.5863778 | 0.0024473 | 0.0179029 |
| PACS1 | 0.5858382 | 0.0032238 | 0.0207518 |
| ADDG | 0.5848681 | 0.0093904 | 0.0387355 |
| T22D2 | 0.5847529 | 0.0037011 | 0.022649 |
| ELAV2 | 0.5836916 | 0.0057817 | 0.0288202 |
| SE6L2 | 0.5827936 | 0.0025168 | 0.0180506 |
| ACTY | 0.5827434 | 0.0008529 | 0.0128895 |

| | | | |
|---|---|---|---|
| PGM2L | 0.5811249 | 0.001717 | 0.0157054 |
| OGT1 | 0.5799179 | 0.0017934 | 0.0157164 |
| DIRA2 | 0.5764043 | 0.0079594 | 0.0348744 |
| DKK3 | 0.5762962 | 0.013775 | 0.0492951 |
| RASK | 0.5739795 | 0.0116748 | 0.0445298 |
| INP4A | 0.5732741 | 0.0026363 | 0.0184752 |
| RAE1 | 0.5720007 | 0.0003345 | 0.0112859 |
| CPNE5 | 0.5712159 | 0.0062262 | 0.030097 |
| F16B1 | 0.568069 | 0.0043645 | 0.0246016 |
| COQ3 | 0.5659828 | 0.0011594 | 0.0140023 |
| TBCD1 | 0.565696 | 0.0117665 | 0.044777 |
| PHP14 | 0.5650845 | 0.0096672 | 0.0395332 |
| TBB3 | 0.5633164 | 0.0046867 | 0.025615 |
| PGES2 | 0.5594736 | 0.0035056 | 0.0218931 |
| PACN2 | 0.5593255 | 0.0072564 | 0.0331797 |
| NDUS2 | 0.5591205 | 0.0099645 | 0.0402752 |
| LRC57 | 0.5582082 | 0.0093857 | 0.0387355 |
| DGKB | 0.5579769 | 0.0092395 | 0.0382037 |
| DNJB6_CON. | 0.5578901 | 0.0014812 | 0.0154658 |
| PITH1 | 0.5574276 | 0.0025467 | 0.0181894 |
| AMOT | 0.5568272 | 0.0083522 | 0.0358689 |
| SCO1 | 0.5566369 | 0.0073209 | 0.0333191 |
| ADT2 | 0.556192 | 0.0136225 | 0.0489241 |
| PTPRD | 0.5542022 | 0.0003194 | 0.0112458 |
| GAL3B.GAL3 | 0.551212 | 0.0013932 | 0.0149305 |
| TANC2 | 0.5511546 | 0.0062 | 0.030097 |
| RHG32 | 0.5509591 | 0.000579 | 0.0121069 |
| PTPRS | 0.5508476 | 0.0013707 | 0.0148279 |
| MECR | 0.5487343 | 0.0015209 | 0.0155249 |
| SPTN4 | 0.5482421 | 0.0034981 | 0.0218931 |
| KT3K | 0.5446824 | 0.0030722 | 0.0201697 |
| ZDHC5 | 0.5445858 | 0.0016099 | 0.0156495 |
| FAHD1 | 0.5423815 | 0.0101991 | 0.0408272 |
| DGLA | 0.5422444 | 0.0077137 | 0.0342585 |
| TEX2 | 0.5403908 | 0.002474 | 0.0179157 |
| PPM1K | 0.5389349 | 0.0070316 | 0.0326702 |
| OSCP1 | 0.5373518 | 0.0031249 | 0.0203011 |
| CATD | 0.5369586 | 0.0095656 | 0.0392466 |
| ASGL1 | 0.5368102 | 0.0016198 | 0.0156495 |
| GPD1L | 0.5366021 | 0.0086772 | 0.0368726 |
| SEPT7 | 0.5326089 | 0.0104248 | 0.0414607 |
| MTMR2 | 0.5292824 | 0.005072 | 0.0266857 |
| SHOT1 | 0.5291522 | 0.0093396 | 0.038587 |
| GPX4 | 0.5286217 | 0.003714 | 0.022649 |
| ASTN1 | 0.5277183 | 0.0045688 | 0.0252401 |
| F262 | 0.5236278 | 0.0019738 | 0.0163878 |
| APBB1 | 0.5233835 | 0.0066544 | 0.031513 |
| GIT1 | 0.5228861 | 0.0013905 | 0.0149305 |

| | | | |
|---|---|---|---|
| KS6A2 | 0.5222551 | 0.0020996 | 0.0168018 |
| GBRL2 | 0.5217753 | 0.007164 | 0.0329392 |
| GBB2 | 0.5215068 | 0.0014772 | 0.0154658 |
| GMPR1 | 0.5181299 | 0.0061122 | 0.029784 |
| NDUB9 | 0.5177305 | 0.0056232 | 0.0282733 |
| AOFA | 0.5175889 | 0.0044729 | 0.0248369 |
| SEPT11 | 0.5166323 | 0.0068458 | 0.0321566 |
| KATL1 | 0.5159528 | 0.0033501 | 0.021302 |
| GBG4 | 0.5159467 | 0.0088538 | 0.0373796 |
| MPP2 | 0.515903 | 0.0022396 | 0.0172874 |
| MFR1L | 0.5154889 | 0.0075116 | 0.033851 |
| AK1C1 | 0.5140242 | 0.0103185 | 0.0411948 |
| MYO1D | 0.5139981 | 0.0111268 | 0.043198 |
| ARFG1 | 0.5138338 | 0.0090736 | 0.0378192 |
| CRAC1 | 0.5137542 | 0.0082544 | 0.0356572 |
| OTUB1 | 0.512488 | 0.0037091 | 0.022649 |
| K1671 | 0.5118979 | 0.0052446 | 0.0272092 |
| ABR | 0.5105062 | 0.0017013 | 0.0157054 |
| EFHD2 | 0.5102343 | 0.0002573 | 0.0109942 |
| ARMC1 | 0.5096903 | 0.01218 | 0.0455896 |
| TRIM3 | 0.509668 | 0.0022454 | 0.0172874 |
| HEM2 | 0.5095992 | 0.0019571 | 0.0163529 |
| KIF1A | 0.5095344 | 0.0125561 | 0.0463979 |
| BABA1 | 0.5094212 | 0.0078433 | 0.0345477 |
| CHP3 | 0.5088843 | 0.0076249 | 0.0340438 |
| DPYL1 | 0.5086692 | 0.0140301 | 0.0499584 |
| ABLM1 | 0.5072803 | 0.0011558 | 0.0140023 |
| CISY | 0.5064483 | 0.0127236 | 0.0468177 |
| PI4KA | 0.5044189 | 0.0123219 | 0.0457594 |
| AP2S1 | 0.5020966 | 0.0122448 | 0.0456661 |
| CPTP | 0.5017002 | 0.0119698 | 0.0450475 |
| STAM1 | 0.5014464 | 0.0018806 | 0.0160561 |
| TIM9 | 0.500266 | 0.0128733 | 0.0470048 |
| ADAP1 | 0.4995552 | 0.0118935 | 0.0449741 |
| NPS3A | 0.498804 | 0.0021265 | 0.0169015 |
| SUCA | 0.4983889 | 0.0078437 | 0.0345477 |
| DPYL4 | 0.498312 | 0.0040289 | 0.0236107 |
| GPC5B | 0.4982304 | 0.0130177 | 0.0473321 |
| THNS1 | 0.4980887 | 0.0019629 | 0.0163756 |
| NRX2A | 0.4968987 | 0.0043879 | 0.0246664 |
| PHYD1 | 0.4942054 | 0.0046995 | 0.0256307 |
| TOLIP | 0.4938374 | 0.0067499 | 0.0318786 |
| PI42A | 0.4914972 | 0.0059955 | 0.0294432 |
| PPCEL | 0.4895274 | 0.0053793 | 0.0275992 |
| PFKAM | 0.4875399 | 0.0019813 | 0.0164234 |
| SRBS2 | 0.4861865 | 0.0037608 | 0.0227481 |
| FRDA | 0.485755 | 0.0088803 | 0.0374307 |
| ACS2L | 0.4852199 | 0.0017087 | 0.0157054 |

| | | | |
|---|---|---|---|
| ABI1 | 0.4794519 | 0.0062251 | 0.030097 |
| TMEM9 | 0.4790207 | 0.0080538 | 0.0351694 |
| MTCH1 | 0.4776384 | 0.0071171 | 0.0328145 |
| NUDT3 | 0.4775479 | 0.0022316 | 0.0172874 |
| NBEA | 0.4774185 | 0.0086184 | 0.0366526 |
| KC1D | 0.4764244 | 0.0016745 | 0.0157032 |
| NDRG3 | 0.4760717 | 0.0010278 | 0.0136032 |
| AGAP3 | 0.4759193 | 0.0121694 | 0.0455829 |
| TLN2 | 0.4758875 | 0.0054613 | 0.0278623 |
| PTPA | 0.4755612 | 0.0040345 | 0.0236107 |
| HINT1 | 0.4751439 | 0.0049012 | 0.0262368 |
| PHIPL | 0.4746143 | 0.0084378 | 0.0361501 |
| AMD | 0.4741642 | 0.0035616 | 0.0220286 |
| MARK3 | 0.4737398 | 0.0006096 | 0.0121307 |
| OCRL | 0.4732897 | 0.0018508 | 0.0159504 |
| NIPS1 | 0.4730977 | 0.0074707 | 0.0337118 |
| WDR13 | 0.4713713 | 0.0006874 | 0.0123592 |
| MTX2 | 0.4712174 | 0.0072162 | 0.0330624 |
| F1711 | 0.4707165 | 0.005367 | 0.0275973 |
| SYSM | 0.4663528 | 0.00352 | 0.0219117 |
| RFIP5 | 0.4662339 | 0.0071632 | 0.0329392 |
| ACTZ | 0.4651413 | 0.0011404 | 0.0140023 |
| ATP9A | 0.463735 | 0.0011874 | 0.0140408 |
| SL9A6 | 0.4634574 | 0.0020967 | 0.0168018 |
| MAST1 | 0.4634228 | 0.0097551 | 0.0398613 |
| PRUN1 | 0.4625415 | 0.0010298 | 0.0136032 |
| HINT3 | 0.4621436 | 0.0074526 | 0.0337017 |
| TBC24 | 0.4621265 | 0.0114858 | 0.0442295 |
| HS105 | 0.4610524 | 0.0050528 | 0.0266857 |
| MK03 | 0.4609997 | 0.0029075 | 0.0196168 |
| SIR5 | 0.4604599 | 0.002908 | 0.0196168 |
| GSTM2 | 0.4604332 | 0.0130488 | 0.047379 |
| X2ABB | 0.4585783 | 0.0039029 | 0.0232512 |
| PDK3 | 0.458406 | 0.0078659 | 0.0346101 |
| SNX30 | 0.4575111 | 0.001609 | 0.0156495 |
| ITSN2 | 0.4567614 | 0.0028918 | 0.0196073 |
| HECW2 | 0.4560554 | 0.0076263 | 0.0340438 |
| NUBPL | 0.4553666 | 0.0062207 | 0.030097 |
| RELCH | 0.4548769 | 0.0069048 | 0.0323753 |
| TOM1 | 0.4536653 | 0.0024416 | 0.0179029 |
| SDHB | 0.4533471 | 0.010788 | 0.0423008 |
| ATG2B | 0.4524219 | 0.0074516 | 0.0337017 |
| AP1B1 | 0.4476736 | 0.0073865 | 0.0335483 |
| S27A4 | 0.4469781 | 0.009899 | 0.0401035 |
| RASM | 0.4433873 | 0.005319 | 0.0274859 |
| REPS2 | 0.4422706 | 0.0052617 | 0.0272711 |
| ASAP1 | 0.4418197 | 0.012853 | 0.0470048 |
| ARP10 | 0.4413299 | 0.0023327 | 0.017657 |

| | | | |
|---|---|---|---|
| PKP4 | 0.4398585 | 0.0041378 | 0.0238918 |
| KI21A | 0.4393083 | 0.0088355 | 0.0373629 |
| THIL | 0.4392202 | 0.0101402 | 0.0406702 |
| MRVI1 | 0.4376827 | 0.0110136 | 0.0429185 |
| DHB8 | 0.4374846 | 0.0024541 | 0.0179029 |
| MARK1 | 0.4374295 | 0.004669 | 0.0255715 |
| SHC3 | 0.4368745 | 0.0035343 | 0.0219117 |
| PIPNA | 0.4368465 | 0.0032995 | 0.0211085 |
| VATG1 | 0.4367367 | 0.0025455 | 0.0181894 |
| GSTO1 | 0.4367036 | 0.0050826 | 0.0266875 |
| SNX4 | 0.4357321 | 0.0033308 | 0.021231 |
| TBCC | 0.4356724 | 0.0045928 | 0.0253041 |
| LZTL1 | 0.4355812 | 0.0103121 | 0.0411948 |
| IMPA1 | 0.4353573 | 0.0118059 | 0.0447984 |
| ACDSB | 0.4350184 | 0.0037664 | 0.0227481 |
| RCAN1 | 0.4343272 | 0.001697 | 0.0157054 |
| BAG4 | 0.4332784 | 0.0108464 | 0.042489 |
| GEPH | 0.4329556 | 0.0128297 | 0.0470048 |
| TTC1 | 0.4319375 | 0.0124638 | 0.0461305 |
| AP3M2 | 0.4315837 | 0.0104419 | 0.0414969 |
| HEMH | 0.4305748 | 0.0039625 | 0.0233717 |
| MTX3 | 0.4298326 | 0.0050166 | 0.0265873 |
| CACO1 | 0.4288391 | 0.005191 | 0.0270659 |
| KIF2A | 0.4248421 | 0.0076798 | 0.0341369 |
| DIP2C | 0.4228889 | 0.0049437 | 0.0263874 |
| CA198 | 0.4214535 | 0.0055462 | 0.0280298 |
| CRYL1 | 0.4130731 | 0.0094326 | 0.0388174 |
| FHIT | 0.4128811 | 0.003924 | 0.0232762 |
| SRGP3 | 0.4124124 | 0.0017416 | 0.0157054 |
| SATT | 0.4114085 | 0.0098567 | 0.0400567 |
| CDK14 | 0.4099556 | 0.0040952 | 0.0238306 |
| ARFP2 | 0.409616 | 0.0069379 | 0.03244 |
| PDPK1 | 0.4079653 | 0.00534 | 0.0275305 |
| SDHF2 | 0.4068986 | 0.0073715 | 0.033509 |
| DCTN1 | 0.4038374 | 0.009594 | 0.0393262 |
| NFU1 | 0.402126 | 0.0139221 | 0.049651 |
| RGN | 0.4019816 | 0.0098253 | 0.0400227 |
| LRP1B | 0.3992829 | 0.0118887 | 0.0449741 |
| FSD1 | 0.3973363 | 0.009567 | 0.0392466 |
| CDK5 | 0.3969403 | 0.008899 | 0.0374791 |
| CC127 | 0.394763 | 0.0078262 | 0.0345477 |
| NNRE | 0.394663 | 0.0011537 | 0.0140023 |
| CRBN | 0.3946431 | 0.013237 | 0.0478956 |
| BRK1 | 0.3945324 | 0.0064358 | 0.0307291 |
| ITSN1 | 0.3942683 | 0.0128498 | 0.0470048 |
| TB10B | 0.3937005 | 0.0050177 | 0.0265873 |
| KIF3A | 0.3933607 | 0.0050588 | 0.0266857 |
| AKT3 | 0.3924572 | 0.0099185 | 0.0401513 |

| | | | |
|---|---|---|---|
| MY18A | 0.3903603 | 0.0126131 | 0.0465755 |
| UBE2O | 0.3887025 | 0.0048518 | 0.0261066 |
| PCCB | 0.387924 | 0.0095237 | 0.0391304 |
| SMAP1 | 0.384505 | 0.0077834 | 0.0344795 |
| DMXL1 | 0.3836932 | 0.0066763 | 0.0315596 |
| DCTN6 | 0.3824245 | 0.0023668 | 0.0177867 |
| KAPCB | 0.3795741 | 0.0128095 | 0.0470012 |
| EP15R | 0.3759199 | 0.0106664 | 0.0420366 |
| ACYP1 | 0.3723422 | 0.0117739 | 0.044777 |
| DCTN4 | 0.3662842 | 0.0027105 | 0.0187432 |
| ANKY2 | 0.3637993 | 0.0123086 | 0.0457427 |
| NFS1 | 0.3621007 | 0.0059182 | 0.029221 |
| ATE1 | 0.3611159 | 0.0021708 | 0.0170918 |
| ARC1A | 0.3574792 | 0.0107322 | 0.0422002 |
| TAXB1 | 0.3541057 | 0.0046091 | 0.0253231 |
| NIF3L | 0.3536058 | 0.0063037 | 0.0302653 |
| GRB2 | 0.3528316 | 0.0097744 | 0.0399049 |
| HSBP1 | 0.3521474 | 0.0088538 | 0.0373796 |
| TMED8 | 0.3472883 | 0.0038098 | 0.022895 |
| ITPK1 | 0.3471624 | 0.0043491 | 0.0245947 |
| TTC9A | 0.3450463 | 0.0120777 | 0.0453696 |
| MTMR1 | 0.336632 | 0.0074735 | 0.0337118 |
| CCD91 | 0.3294548 | 0.0128736 | 0.0470048 |
| GRAP1 | 0.3282645 | 0.0100728 | 0.0404931 |
| PP1R7 | 0.3260338 | 0.0127111 | 0.0468177 |
| TACO1 | 0.3242182 | 0.0110761 | 0.0431299 |
| ABHD6 | 0.322164 | 0.0048366 | 0.0260972 |
| TPPC9 | 0.3151689 | 0.0113538 | 0.043916 |
| SIK3 | 0.3138586 | 0.0100513 | 0.0404378 |
| UH1BL | 0.3119087 | 0.012648 | 0.0466714 |
| MAT2B | 0.3008298 | 0.0109024 | 0.0426444 |
| CAMP1 | 0.2971318 | 0.0122229 | 0.0456661 |

| ID | FC | P.Value | adj.P.Val |
|---|---|---|---|
| H15 | -1.5469313 | 0.0006202 | 0.0121479 |
| FINC | -1.4611496 | 0.0021785 | 0.0170918 |
| SERPH | -1.4097686 | 0.0006133 | 0.0121472 |
| LAMA4 | -1.3953476 | 0.0000101 | 0.0098936 |
| IL1AP | -1.3505287 | 0.0013607 | 0.0147828 |
| FCGRN | -1.2977881 | 0.0042534 | 0.0243263 |
| H33 | -1.2559667 | 0.0003988 | 0.0114048 |
| NAMPT | -1.2183154 | 0.0090986 | 0.0378613 |
| K2C74 | -1.2169992 | 0.0118351 | 0.044846 |
| OSTC | -1.1990707 | 0.00042 | 0.0114048 |
| H4 | -1.1761419 | 0.0072805 | 0.0332402 |
| CO5A2 | -1.1520036 | 0.002054 | 0.0167092 |
| PDIA4 | -1.1365782 | 0.0028216 | 0.0193069 |
| CASP3 | -1.1241226 | 0.0109945 | 0.0429011 |
| PDIA1 | -1.1179258 | 0.0018291 | 0.015866 |
| APOBR | -1.1156923 | 0.004255 | 0.0243263 |
| CLIC1 | -1.110098 | 0.0106076 | 0.0419002 |
| CO5A3 | -1.1089953 | 0.0024406 | 0.0179029 |
| TM201 | -1.1075643 | 0.0006963 | 0.0123592 |
| P3H1 | -1.091449 | 0.0016809 | 0.0157054 |
| YBOX1 | -1.0878482 | 0.0021881 | 0.0171048 |
| LMNB1 | -1.0511236 | 0.0006648 | 0.0123201 |
| S39A7 | -1.0494883 | 0.0001253 | 0.0098936 |
| FLNA | -1.0337484 | 0.0119617 | 0.0450475 |
| MRC2 | -1.0291347 | 0.0024947 | 0.0180161 |
| PROF1 | -1.0286644 | 0.0045286 | 0.0250926 |
| RIR1 | -1.0212375 | 0.0000101 | 0.0098936 |
| SC11A | -1.0187395 | 0.002911 | 0.0196168 |
| RL15 | -1.0140529 | 0.0008162 | 0.0127211 |
| BUD31 | -1.0130407 | 0.0001143 | 0.0098936 |
| SSRG | -0.9882183 | 0.0003783 | 0.0114048 |
| TIMP1 | -0.981449 | 0.0136338 | 0.0489241 |
| SSRA | -0.9770929 | 0.0003279 | 0.0112859 |
| LAMB1 | -0.9681073 | 0.0000997 | 0.0098936 |
| PCNA | -0.9669212 | 0.0021154 | 0.0168688 |
| IKIP | -0.9585851 | 0.0016342 | 0.0156495 |
| AL5AP | -0.9506246 | 0.0046648 | 0.0255715 |
| CRTAP | -0.9501307 | 0.0027041 | 0.0187432 |
| RL18 | -0.9404537 | 0.000126 | 0.0098936 |
| BIP | -0.9378155 | 0.0015137 | 0.0155249 |
| IBP2 | -0.9255767 | 0.0030227 | 0.0200511 |
| RPN1 | -0.9244915 | 0.0009834 | 0.0134835 |
| AN32E | -0.9171287 | 0.0028019 | 0.0192228 |
| SMD3 | -0.915294 | 0.0001312 | 0.0098936 |
| PPIB | -0.9095946 | 0.0005531 | 0.0120166 |
| MYL6 | -0.9093652 | 0.0015157 | 0.0155249 |
| RL13A | -0.9085735 | 0.0000628 | 0.0098936 |

| | | | |
|---|---|---|---|
| H2A1B.H2A3 | -0.9050648 | 0.0057938 | 0.0288527 |
| PTN | -0.9040248 | 0.0064707 | 0.0308674 |
| PLOD3 | -0.9033392 | 0.0068619 | 0.0322033 |
| SMCA5 | -0.9007038 | 0.0047228 | 0.0256774 |
| STT3A | -0.8990673 | 0.0002029 | 0.0100934 |
| CKAP4 | -0.8952694 | 0.0010994 | 0.01388 |
| FHL3 | -0.8909838 | 0.0013547 | 0.0147767 |
| PLCH1 | -0.8896492 | 0.0043447 | 0.0245947 |
| PGBM | -0.8892803 | 0.0001167 | 0.0098936 |
| EMIL1 | -0.8882703 | 0.0016904 | 0.0157054 |
| FADS2 | -0.887129 | 0.0039516 | 0.0233601 |
| PODXL | -0.8811612 | 0.0020671 | 0.0167132 |
| RPN2 | -0.8793632 | 0.0024721 | 0.0179157 |
| ROA1 | -0.878183 | 0.0006658 | 0.0123201 |
| CALU | -0.8774025 | 0.0012411 | 0.0142673 |
| CRIP1 | -0.8742565 | 0.0062087 | 0.030097 |
| LBR | -0.8732277 | 0.0042681 | 0.0243435 |
| LTBP3 | -0.8615404 | 0.0015164 | 0.0155249 |
| RS9 | -0.8592064 | 0.0008053 | 0.0126095 |
| OLIG1 | -0.8556826 | 0.0038151 | 0.022895 |
| LAMC1 | -0.855538 | 0.0000219 | 0.0098936 |
| HMGB3 | -0.8550638 | 0.0041788 | 0.0240467 |
| LMAN1 | -0.8541532 | 0.0018815 | 0.0160561 |
| CO4A2 | -0.8537988 | 0.0007818 | 0.0125466 |
| EF2 | -0.8535002 | 0.0000502 | 0.0098936 |
| ENPL | -0.849034 | 0.0063294 | 0.0303603 |
| OST48 | -0.8490061 | 0.0002219 | 0.0104472 |
| DX39A | -0.8480058 | 0.0062215 | 0.030097 |
| PTBP1 | -0.8451998 | 0.0010366 | 0.0136139 |
| BCAT1 | -0.8429145 | 0.0011507 | 0.0140023 |
| CALR | -0.8421426 | 0.0022234 | 0.0172767 |
| PRDX4 | -0.8409541 | 0.0028057 | 0.0192231 |
| FKB10 | -0.8397555 | 0.0016731 | 0.0157032 |
| SSRD | -0.839484 | 0.001141 | 0.0140023 |
| FEN1 | -0.8383098 | 0.0035286 | 0.0219117 |
| CALX | -0.8363578 | 0.0005227 | 0.0118593 |
| MCM3 | -0.8346122 | 0.0031304 | 0.0203011 |
| IQGA1 | -0.8345259 | 0.0062301 | 0.030097 |
| RNH2B | -0.8310028 | 0.0075771 | 0.033945 |
| RL7 | -0.8292442 | 0.0001142 | 0.0098936 |
| ITA5 | -0.8245376 | 0.0031809 | 0.0205271 |
| PTH2 | -0.8240791 | 0.0025612 | 0.0182249 |
| SMD1 | -0.8173902 | 0.0024057 | 0.017854 |
| PTBP3 | -0.8161865 | 0.0110851 | 0.0431325 |
| MPRI | -0.8090397 | 0.0046065 | 0.0253231 |
| PDIA6 | -0.8059598 | 0.0069767 | 0.0325339 |
| MYD88 | -0.8019654 | 0.0047588 | 0.0258194 |
| APEX1 | -0.8015319 | 0.0001707 | 0.0100011 |

| | | | |
|---|---|---|---|
| HMGB1 | -0.7985304 | 0.0101347 | 0.0406702 |
| COIA1 | -0.7937581 | 0.0004282 | 0.0114048 |
| PDIA3 | -0.7855245 | 0.0024547 | 0.0179029 |
| MYH9 | -0.7852497 | 0.0047195 | 0.0256774 |
| GGH | -0.785016 | 0.0001866 | 0.0100934 |
| GT251 | -0.782944 | 0.0028796 | 0.0196008 |
| EF1A1 | -0.7823561 | 0.0050996 | 0.0267496 |
| VWF | -0.7773304 | 0.0004331 | 0.0114048 |
| TM100 | -0.7762523 | 0.0123341 | 0.0457721 |
| PHF6 | -0.7692959 | 0.0001057 | 0.0098936 |
| H2AZ.H2AV | -0.768009 | 0.0116729 | 0.0445298 |
| MGN.MGN2 | -0.7673728 | 0.0004293 | 0.0114048 |
| WIZ | -0.7668901 | 0.0058573 | 0.0290709 |
| SSRP1 | -0.7634452 | 0.0018977 | 0.0160604 |
| NID1 | -0.7607975 | 0.0001234 | 0.0098936 |
| U2AF5.U2AF | -0.7561732 | 0.0000911 | 0.0098936 |
| BAZ1B | -0.7503422 | 0.0020574 | 0.0167092 |
| BAF | -0.7492003 | 0.0008695 | 0.0128902 |
| TMEDA | -0.7472249 | 0.0014946 | 0.0154658 |
| TIF1B | -0.7455681 | 0.0011049 | 0.0139146 |
| FKBP9 | -0.7431884 | 0.0127214 | 0.0468177 |
| MATN2 | -0.7417867 | 0.0131964 | 0.047782 |
| TOP1 | -0.7411467 | 0.0006021 | 0.0121069 |
| ANKH1 | -0.7382838 | 0.0004913 | 0.0118046 |
| TLN1 | -0.7363599 | 0.0046752 | 0.0255786 |
| RASF2 | -0.7351481 | 0.0023675 | 0.0177867 |
| RL36A | -0.7340091 | 0.0010127 | 0.0135736 |
| VASP | -0.7335108 | 0.0076328 | 0.0340438 |
| PGLT1 | -0.73116 | 0.0007345 | 0.0124668 |
| MAGT1 | -0.7279936 | 0.0010052 | 0.0135177 |
| PDS5A | -0.7248402 | 0.0128686 | 0.0470048 |
| LAP2B | -0.7239735 | 0.0012951 | 0.014491 |
| LIMA1 | -0.7222296 | 0.0013945 | 0.0149305 |
| FBRL | -0.7219719 | 0.0003545 | 0.0113391 |
| RDH10 | -0.7210865 | 0.011657 | 0.0445298 |
| HMGA1 | -0.7201169 | 0.0105185 | 0.0416517 |
| MET16 | -0.7191778 | 0.0006969 | 0.0123592 |
| MPP10 | -0.7189021 | 0.0017539 | 0.0157054 |
| NID2 | -0.7155839 | 0.0001328 | 0.0098936 |
| AFG2H | -0.7096855 | 0.0065066 | 0.0309823 |
| AKAP8 | -0.709412 | 0.0002548 | 0.0109942 |
| RS4X | -0.7083771 | 0.0006871 | 0.0123592 |
| TMX1 | -0.7080227 | 0.0003481 | 0.0113186 |
| RS15 | -0.7067643 | 0.0069823 | 0.0325339 |
| RL1D1 | -0.706619 | 0.0135006 | 0.0486498 |
| CSPG4 | -0.7052288 | 0.0000925 | 0.0098936 |
| RS23 | -0.7048947 | 0.0010776 | 0.0138149 |
| S35U4 | -0.7017973 | 0.0058599 | 0.0290709 |

| | | | |
|---|---|---|---|
| RS10 | -0.7014452 | 0.0012624 | 0.0143714 |
| RUXGL.RUXG | -0.7008175 | 0.0003889 | 0.0114048 |
| DD19B | -0.7007121 | 0.0079559 | 0.0348744 |
| IDHC | -0.7006376 | 0.0116271 | 0.0445298 |
| ROAA | -0.7003698 | 0.004878 | 0.0261398 |
| RPAB1 | -0.7000223 | 0.0016244 | 0.0156495 |
| XRCC5 | -0.6999355 | 0.001035 | 0.0136139 |
| XRCC6 | -0.697315 | 0.0007181 | 0.0124393 |
| RBM8A | -0.6964091 | 0.0008257 | 0.0127211 |
| DNJC3 | -0.6959758 | 0.0005321 | 0.0118593 |
| RLA0 | -0.6954306 | 0.0002974 | 0.0112381 |
| RACK1 | -0.6930365 | 0.0001246 | 0.0098936 |
| BAG1 | -0.6925797 | 0.0022969 | 0.0174581 |
| RUXF | -0.6923302 | 0.0043979 | 0.0246664 |
| TBL2 | -0.6921399 | 0.0011918 | 0.0140408 |
| DJB11 | -0.6872874 | 0.0046938 | 0.0256266 |
| RL32 | -0.6869095 | 0.0007327 | 0.0124668 |
| TTC38 | -0.6852156 | 0.0016503 | 0.0156495 |
| PSMG2 | -0.6849963 | 0.0031123 | 0.0203011 |
| TMM43 | -0.6846447 | 0.0020512 | 0.0167092 |
| MYO9B | -0.683531 | 0.001787 | 0.0157164 |
| RCC2 | -0.6827566 | 0.001768 | 0.0157164 |
| DPM1 | -0.6799942 | 0.0007695 | 0.0124986 |
| HNRPM | -0.6746299 | 0.0035316 | 0.0219117 |
| RL31 | -0.671351 | 0.0089171 | 0.0375103 |
| SYNE2 | -0.6711688 | 0.0039737 | 0.0234111 |
| AGRIN | -0.6703739 | 0.012466 | 0.0461305 |
| MSH2 | -0.669274 | 0.0007683 | 0.0124986 |
| RS5 | -0.6687773 | 0.0011135 | 0.0139231 |
| CBX5 | -0.6683962 | 0.0011601 | 0.0140023 |
| PEBB | -0.667452 | 0.012271 | 0.0456681 |
| MRP | -0.6669059 | 0.0133246 | 0.0481052 |
| UHRF1 | -0.6668503 | 0.0015775 | 0.0156366 |
| DX39B | -0.6652099 | 0.0025097 | 0.018049 |
| RU2A | -0.6647962 | 0.0015344 | 0.0155647 |
| GOLI4 | -0.6635984 | 0.0002325 | 0.0107278 |
| HNRPQ | -0.6630139 | 0.0003476 | 0.0113186 |
| IF4A1 | -0.6602941 | 0.0009587 | 0.0134729 |
| RBBP4 | -0.6575009 | 0.0011902 | 0.0140408 |
| TMED9 | -0.6565934 | 0.0005366 | 0.0118593 |
| PAXI | -0.6554634 | 0.0083322 | 0.0358452 |
| DCK | -0.6553091 | 0.000289 | 0.0110797 |
| VWA1 | -0.6545235 | 0.000267 | 0.011047 |
| HNRPU | -0.6541 | 0.0007893 | 0.0125466 |
| EMIL2 | -0.6528396 | 0.0029867 | 0.0199028 |
| RO60 | -0.6524983 | 0.0005909 | 0.0121069 |
| HNRPC | -0.6518485 | 0.0016215 | 0.0156495 |
| RS16 | -0.6513742 | 0.0057517 | 0.0287253 |

| | | | |
|---|---|---|---|
| MCM5 | -0.6473534 | 0.0066215 | 0.0314429 |
| SYMPK | -0.6455845 | 0.0020653 | 0.0167132 |
| GLU2B | -0.6452652 | 0.0017303 | 0.0157054 |
| RL10 | -0.6445307 | 0.001329 | 0.0146615 |
| HNRH1 | -0.6442973 | 0.0009817 | 0.0134835 |
| MCM7 | -0.6423491 | 0.0043972 | 0.0246664 |
| TRA2A | -0.6420573 | 0.0005477 | 0.0119477 |
| VANG2 | -0.6417185 | 0.0026589 | 0.0185343 |
| TGFI1 | -0.6412016 | 0.0070566 | 0.032705 |
| CBX3 | -0.6400425 | 0.0053108 | 0.0274709 |
| PAPS1 | -0.6393486 | 0.0029812 | 0.0199025 |
| RS2 | -0.6381973 | 0.0011529 | 0.0140023 |
| TMED4 | -0.6366794 | 0.0020369 | 0.0167092 |
| PP1RA | -0.6362282 | 0.0080338 | 0.0351402 |
| SF3A3 | -0.6361409 | 0.0006056 | 0.0121069 |
| TIA1 | -0.6343018 | 0.0070366 | 0.0326702 |
| ERH | -0.6322969 | 0.0004539 | 0.0115456 |
| BZW1 | -0.6320124 | 0.0014094 | 0.0149974 |
| GANAB | -0.6319676 | 0.0050454 | 0.0266801 |
| RS26 | -0.630532 | 0.0090057 | 0.0376956 |
| FA50A | -0.6298713 | 0.0063932 | 0.0306098 |
| TOM40 | -0.6293035 | 0.0069799 | 0.0325339 |
| IMDH2 | -0.6292606 | 0.0048575 | 0.0261101 |
| SMHD1 | -0.629244 | 0.0018936 | 0.0160604 |
| RS17 | -0.6292414 | 0.001704 | 0.0157054 |
| RU2B | -0.6291475 | 0.0001684 | 0.0099806 |
| H2AY | -0.6287439 | 0.0000922 | 0.0098936 |
| ACL6A | -0.6277972 | 0.0029131 | 0.0196168 |
| TXND5 | -0.6262977 | 0.0096493 | 0.039491 |
| SAP18 | -0.6248969 | 0.0016541 | 0.0156495 |
| SMC4 | -0.6241806 | 0.0023657 | 0.0177867 |
| RL18A | -0.6208282 | 0.0014979 | 0.0154658 |
| RCN1 | -0.6205384 | 0.0009843 | 0.0134835 |
| PRKDC | -0.6204698 | 0.0009922 | 0.0135074 |
| RL35A | -0.6196742 | 0.0031602 | 0.0204432 |
| RS3 | -0.6189089 | 0.0023262 | 0.0176548 |
| U5S1 | -0.6171974 | 0.0002355 | 0.0107722 |
| RS8 | -0.6167784 | 0.0021926 | 0.0171141 |
| SRRT | -0.616573 | 0.0006464 | 0.012285 |
| PLGT3 | -0.6156159 | 0.0138107 | 0.0493887 |
| RS18 | -0.6148558 | 0.0006778 | 0.0123592 |
| SC24A | -0.6145953 | 0.0107676 | 0.0423008 |
| RS27 | -0.614436 | 0.0060022 | 0.0294432 |
| AN32A | -0.6139969 | 0.0102029 | 0.0408272 |
| RL38 | -0.6139542 | 0.0020305 | 0.0167092 |
| FCSD2 | -0.6139242 | 0.0086933 | 0.036881 |
| DCTP1 | -0.6132305 | 0.0030456 | 0.0201074 |
| HNRL1 | -0.6124747 | 0.0011956 | 0.0140408 |

| | | | |
|---|---|---|---|
| PPIG | -0.6122627 | 0.0008727 | 0.0128902 |
| RS13 | -0.6119703 | 0.0005318 | 0.0118593 |
| LA | -0.6112482 | 0.0041444 | 0.0239036 |
| SC23B | -0.6110505 | 0.0007114 | 0.0124393 |
| SC24D | -0.6068547 | 0.0017897 | 0.0157164 |
| RECQ1 | -0.6055238 | 0.0004676 | 0.0115541 |
| MCM6 | -0.6053856 | 0.0117487 | 0.0447462 |
| GSTK1 | -0.6012824 | 0.0018175 | 0.0158202 |
| RS6 | -0.6006502 | 0.003181 | 0.0205271 |
| AN32B | -0.6001114 | 0.0075686 | 0.033945 |
| BOP1 | -0.6001043 | 0.0017553 | 0.0157054 |
| MYO1B | -0.6000076 | 0.0034552 | 0.0218101 |
| DHX15 | -0.5986754 | 0.0020828 | 0.0167685 |
| RAB31 | -0.5980717 | 0.0004319 | 0.0114048 |
| NUP54 | -0.598004 | 0.0009977 | 0.0135177 |
| MUC18 | -0.597017 | 0.0017586 | 0.0157054 |
| LMF2 | -0.5968154 | 0.0045202 | 0.0250727 |
| HDAC1 | -0.5965065 | 0.002609 | 0.0184094 |
| MA1A2 | -0.5961836 | 0.0026761 | 0.0185793 |
| PSME3 | -0.5948364 | 0.0021197 | 0.0168733 |
| RS3A | -0.5944855 | 0.0006553 | 0.0123201 |
| SPCS2 | -0.5925302 | 0.0002848 | 0.0110797 |
| RL10A | -0.592453 | 0.0014976 | 0.0154658 |
| ILF2 | -0.5917641 | 0.0026002 | 0.0183957 |
| HNRPR | -0.5914679 | 0.0008851 | 0.012953 |
| PPM1G | -0.590642 | 0.0008002 | 0.0126095 |
| ACTN4 | -0.5901862 | 0.0037308 | 0.0226985 |
| RL14 | -0.5898549 | 0.0015686 | 0.0156366 |
| RBM39 | -0.5894144 | 0.0016477 | 0.0156495 |
| PEPL1 | -0.5880246 | 0.0032456 | 0.0208408 |
| GTF2I | -0.5877899 | 0.0043625 | 0.0246016 |
| NHP2 | -0.5871656 | 0.0025665 | 0.0182315 |
| DDX5 | -0.5858932 | 0.0010973 | 0.01388 |
| OFUT1 | -0.5855811 | 0.000601 | 0.0121069 |
| CTBL1 | -0.5844289 | 0.0026427 | 0.018495 |
| COPB | -0.5841775 | 0.0009149 | 0.0130695 |
| NUP62 | -0.583979 | 0.0007659 | 0.0124986 |
| RS7 | -0.5829336 | 0.000696 | 0.0123592 |
| PAPS2 | -0.5829032 | 0.0011343 | 0.0140023 |
| GCSP | -0.5816646 | 0.0078999 | 0.0347303 |
| RL13 | -0.5809409 | 0.0106662 | 0.0420366 |
| HNRPF | -0.580561 | 0.0055227 | 0.027984 |
| RFC3 | -0.5787085 | 0.0054788 | 0.0278937 |
| RPAB3 | -0.5785392 | 0.0100181 | 0.0403808 |
| NUP85 | -0.5782492 | 0.000721 | 0.0124482 |
| PXDN | -0.5754697 | 0.0008302 | 0.0127211 |
| RUVB2 | -0.5735703 | 0.0030754 | 0.0201697 |
| SEC63 | -0.573136 | 0.0010186 | 0.0136032 |

| | | | |
|---|---|---|---|
| NONO | -0.5727192 | 0.0037952 | 0.02285 |
| NCBP1 | -0.5723168 | 0.000166 | 0.0099806 |
| LC7L3 | -0.5708147 | 0.0089652 | 0.0376367 |
| ANM1 | -0.5699637 | 0.0021436 | 0.0169702 |
| SUN2 | -0.5691261 | 0.0067843 | 0.0320058 |
| FERM2 | -0.5688449 | 0.0043697 | 0.0246042 |
| RL9 | -0.5686157 | 0.0033849 | 0.0214445 |
| SON | -0.5673616 | 0.0011888 | 0.0140408 |
| CPNE3 | -0.5671217 | 0.0049445 | 0.0263874 |
| LC7L2 | -0.566842 | 0.0128315 | 0.0470048 |
| NCBP2 | -0.5668409 | 0.0017991 | 0.0157164 |
| PGS1 | -0.5668016 | 0.0004729 | 0.0116308 |
| SRSF7 | -0.5664704 | 0.0087173 | 0.0369529 |
| SMD2 | -0.5663732 | 0.0007406 | 0.0124668 |
| RL27A | -0.5651114 | 0.0097811 | 0.0399049 |
| AGAL | -0.5631782 | 0.0036248 | 0.0223665 |
| LRP1 | -0.5626621 | 0.001721 | 0.0157054 |
| TF2AA | -0.561889 | 0.001838 | 0.015866 |
| ROA3 | -0.5613938 | 0.0015649 | 0.0156366 |
| THYN1 | -0.5603761 | 0.0007151 | 0.0124393 |
| DNMT1 | -0.5601731 | 0.0044138 | 0.0246664 |
| NSRP1 | -0.5588895 | 0.0135731 | 0.0488743 |
| DEGS1 | -0.5570424 | 0.0032765 | 0.0210128 |
| LMAN2 | -0.556678 | 0.0011102 | 0.0139146 |
| SNW1 | -0.5557886 | 0.007118 | 0.0328145 |
| ITB1 | -0.5553952 | 0.0135014 | 0.0486498 |
| COPA | -0.5553263 | 0.0020691 | 0.0167132 |
| RPR1B | -0.554569 | 0.0055012 | 0.0279292 |
| X8ODP | -0.554481 | 0.0062865 | 0.0302583 |
| COPD | -0.5542418 | 0.0012702 | 0.0144286 |
| THOC4 | -0.5538926 | 0.0108699 | 0.0425491 |
| HYOU1 | -0.5536092 | 0.0035152 | 0.0219117 |
| ARL1 | -0.5534868 | 0.0015741 | 0.0156366 |
| RSF1 | -0.5531326 | 0.001065 | 0.0138134 |
| HNRPK | -0.5520758 | 0.0048451 | 0.0260972 |
| ARGL1 | -0.551831 | 0.0060029 | 0.0294432 |
| RAVR1 | -0.5516941 | 0.0107902 | 0.0423008 |
| CFA20 | -0.5503047 | 0.0009207 | 0.0130777 |
| RFA3 | -0.5502634 | 0.0129964 | 0.0472876 |
| NUP37 | -0.5501035 | 0.0018998 | 0.0160604 |
| TFR1 | -0.549776 | 0.0056665 | 0.0284361 |
| NU160 | -0.5496601 | 0.002457 | 0.0179029 |
| TF3C4 | -0.5496258 | 0.0007887 | 0.0125466 |
| LSM2 | -0.5493505 | 0.0022526 | 0.0172874 |
| MCM4 | -0.5487644 | 0.0110009 | 0.0429011 |
| SNR40 | -0.5483641 | 0.0012356 | 0.0142673 |
| RS25 | -0.5479474 | 0.0111959 | 0.0434342 |
| SMCA4 | -0.5473236 | 0.0027736 | 0.019104 |

| | | | |
|---|---|---|---|
| PCAT1 | -0.5471728 | 0.0044347 | 0.0247035 |
| BZW2 | -0.5464051 | 0.0003384 | 0.0112859 |
| RL5 | -0.5458562 | 0.0013061 | 0.0145513 |
| RPB1 | -0.5456957 | 0.0033237 | 0.0212113 |
| ROA0 | -0.5442796 | 0.0024619 | 0.0179029 |
| SBDS | -0.541769 | 0.002409 | 0.017854 |
| SMCE1 | -0.5411968 | 0.0055864 | 0.0281578 |
| MAPK2 | -0.5409194 | 0.0075781 | 0.033945 |
| G3BP1 | -0.5399608 | 0.0016535 | 0.0156495 |
| TR112 | -0.5393642 | 0.0005368 | 0.0118593 |
| SF3B1 | -0.5393284 | 0.0006616 | 0.0123201 |
| RL12 | -0.5380107 | 0.0008609 | 0.0128902 |
| MCM2 | -0.5371626 | 0.0078065 | 0.0344939 |
| FUBP3 | -0.536248 | 0.0085197 | 0.0363815 |
| AT131 | -0.5348098 | 0.003691 | 0.0226423 |
| EFS | -0.5344354 | 0.0033734 | 0.0214239 |
| SP16H | -0.5313015 | 0.0041273 | 0.0238581 |
| DDX23 | -0.5287773 | 0.0008923 | 0.0129963 |
| ADNP | -0.5280296 | 0.0017779 | 0.0157164 |
| RS12 | -0.5278994 | 0.0082001 | 0.0355701 |
| NUCB2 | -0.5270926 | 0.0039078 | 0.0232512 |
| GSDME | -0.5266546 | 0.0009475 | 0.0133885 |
| DCPS | -0.5254507 | 0.0127768 | 0.0469144 |
| ILF3 | -0.5241021 | 0.007231 | 0.0331012 |
| VINC | -0.5223505 | 0.0082675 | 0.0356843 |
| THOC5 | -0.5219661 | 0.0136325 | 0.0489241 |
| GNAI3 | -0.5218949 | 0.0074128 | 0.0336092 |
| SFPQ | -0.5217547 | 0.0075464 | 0.0339196 |
| DNLI3 | -0.5203397 | 0.0055155 | 0.0279746 |
| RS20 | -0.5189939 | 0.0077282 | 0.0342735 |
| TNPO1 | -0.5189104 | 0.0001283 | 0.0098936 |
| SIN3A | -0.5185266 | 0.0017995 | 0.0157164 |
| PARP1 | -0.5171847 | 0.0128965 | 0.0470225 |
| ARF4 | -0.517047 | 0.0050216 | 0.0265873 |
| KHDR1 | -0.5165785 | 0.0011347 | 0.0140023 |
| WDR18 | -0.5161223 | 0.001702 | 0.0157054 |
| YTHD2 | -0.5153533 | 0.0015534 | 0.0156055 |
| RL30 | -0.5143567 | 0.0016326 | 0.0156495 |
| NH2L1 | -0.5142048 | 0.0031583 | 0.0204432 |
| DHX9 | -0.5118317 | 0.0016307 | 0.0156495 |
| RHOC | -0.5116617 | 0.0054968 | 0.0279292 |
| NU133 | -0.5108877 | 0.004183 | 0.0240467 |
| SPF27 | -0.5099272 | 0.0009058 | 0.0130695 |
| U2AF2 | -0.5090348 | 0.0084279 | 0.0361376 |
| DDX17 | -0.5084791 | 0.0008619 | 0.0128902 |
| PMM2 | -0.5078481 | 0.0050661 | 0.0266857 |
| HNRPL | -0.5073026 | 0.0004999 | 0.0118593 |
| S39AE | -0.5044944 | 0.0035268 | 0.0219117 |

| | | | |
|---|---|---|---|
| SAE1 | -0.5044415 | 0.0030665 | 0.0201697 |
| DIDO1 | -0.504378 | 0.0031006 | 0.0202842 |
| SGTA | -0.504319 | 0.0010032 | 0.0135177 |
| PR40A | -0.503979 | 0.0092031 | 0.0381081 |
| ELAV1 | -0.5035396 | 0.0008012 | 0.0126095 |
| SMC2 | -0.5034776 | 0.0100359 | 0.0404072 |
| ITAV | -0.503401 | 0.0035537 | 0.0220059 |
| PRP4B | -0.5025784 | 0.000606 | 0.0121069 |
| ASPH | -0.5019738 | 0.0075375 | 0.033909 |
| PNKP | -0.5017046 | 0.0121614 | 0.0455829 |
| RBBP7 | -0.4995839 | 0.0028888 | 0.0196073 |
| HDGF | -0.4992575 | 0.0038267 | 0.0229336 |
| U520 | -0.4985808 | 0.0012858 | 0.0144765 |
| ANM3 | -0.4984175 | 0.0006276 | 0.0121638 |
| NOP58 | -0.4983759 | 0.0029772 | 0.0199014 |
| TIMP2 | -0.498081 | 0.0037106 | 0.022649 |
| T2FB | -0.4963235 | 0.0076764 | 0.0341369 |
| UBC9 | -0.495674 | 0.0113104 | 0.0437804 |
| DJC10 | -0.4954257 | 0.0031194 | 0.0203011 |
| PRP8 | -0.4946416 | 0.0010423 | 0.0136207 |
| IF5A1 | -0.4938982 | 0.0047492 | 0.0257938 |
| RSSA | -0.4938027 | 0.0037651 | 0.0227481 |
| RL3 | -0.4928312 | 0.0100216 | 0.0403808 |
| AKAP2 | -0.4924543 | 0.0082445 | 0.0356443 |
| RAD50 | -0.4922721 | 0.0099525 | 0.0402579 |
| LRC59 | -0.4920874 | 0.0091917 | 0.0381081 |
| TFCP2 | -0.4919371 | 0.003476 | 0.0218886 |
| ACOX1 | -0.4913747 | 0.0073204 | 0.0333191 |
| CDC5L | -0.4910723 | 0.0050227 | 0.0265873 |
| BAX | -0.4897162 | 0.0122539 | 0.0456661 |
| PKN1 | -0.4894721 | 0.0005732 | 0.0121069 |
| EWS | -0.488094 | 0.0059928 | 0.0294432 |
| SF3A2 | -0.4879488 | 0.0100011 | 0.0403603 |
| P5CR1 | -0.4873325 | 0.0048349 | 0.0260972 |
| PARVA | -0.4865757 | 0.0054008 | 0.0276619 |
| COPE | -0.4865111 | 0.0038415 | 0.0229961 |
| ILK | -0.4855128 | 0.0010543 | 0.0137079 |
| NUMA1 | -0.4844967 | 0.0078451 | 0.0345477 |
| NU205 | -0.4844353 | 0.0011949 | 0.0140408 |
| PCY1A | -0.4839722 | 0.0002797 | 0.011047 |
| AKP8L | -0.4831144 | 0.0031167 | 0.0203011 |
| PABP2 | -0.4830687 | 0.0014616 | 0.0153646 |
| ACINU | -0.4829355 | 0.0094312 | 0.0388174 |
| RS14 | -0.4821135 | 0.0020422 | 0.0167092 |
| PRP6 | -0.4819627 | 0.0039619 | 0.0233717 |
| RALY | -0.4818121 | 0.0077918 | 0.0344877 |
| LIMS1 | -0.4803406 | 0.0029734 | 0.0199014 |
| RL22 | -0.4802725 | 0.008984 | 0.037685 |

| | | | |
|---|---|---|---|
| QKI | -0.480178 | 0.0126696 | 0.0467181 |
| FAM3C | -0.4783862 | 0.0029769 | 0.0199014 |
| DKC1 | -0.4782021 | 0.0019504 | 0.0163353 |
| TADBP | -0.4779482 | 0.0025761 | 0.0182746 |
| SMC1A | -0.4775929 | 0.0080722 | 0.0352061 |
| MAVS | -0.4770562 | 0.0022612 | 0.0172874 |
| SYIC | -0.4765829 | 0.0004909 | 0.0118046 |
| TCEA1 | -0.4764989 | 0.0081711 | 0.0354738 |
| RS11 | -0.4740713 | 0.0111206 | 0.043198 |
| SF3B6 | -0.4729114 | 0.0076078 | 0.0340196 |
| MLEC | -0.4727561 | 0.0005205 | 0.0118593 |
| MTA2 | -0.4720924 | 0.0122554 | 0.0456661 |
| PBDC1 | -0.4720883 | 0.0044409 | 0.0247115 |
| RL4 | -0.47061 | 0.0138293 | 0.0494216 |
| ACSL3 | -0.4699983 | 0.0064321 | 0.0307291 |
| U3IP2 | -0.4693845 | 0.0094513 | 0.0388637 |
| FA98B | -0.4688946 | 0.0041628 | 0.0239832 |
| NFIX | -0.4681133 | 0.0024094 | 0.017854 |
| CCD12 | -0.4664618 | 0.0115205 | 0.0443307 |
| SAFB2 | -0.4659715 | 0.0116747 | 0.0445298 |
| COG8 | -0.4655687 | 0.0039907 | 0.0234845 |
| RL6 | -0.4655148 | 0.004106 | 0.0238391 |
| GCN1 | -0.465128 | 0.0016546 | 0.0156495 |
| RHG18 | -0.4635465 | 0.0032308 | 0.020771 |
| DDX46 | -0.4631131 | 0.0039153 | 0.0232512 |
| PPIL4 | -0.4627543 | 0.0040915 | 0.0238306 |
| MYDGF | -0.462566 | 0.0009532 | 0.013433 |
| IF2P | -0.4612934 | 0.0033465 | 0.021302 |
| TOP2B | -0.4608878 | 0.0104154 | 0.041455 |
| IF2B | -0.4607893 | 0.0010292 | 0.0136032 |
| GLYM | -0.4602674 | 0.0036635 | 0.0225522 |
| SYRC | -0.459012 | 0.0014455 | 0.0152879 |
| RBM12 | -0.4585748 | 0.0020487 | 0.0167092 |
| HNRL2 | -0.4584309 | 0.0016107 | 0.0156495 |
| COPB2 | -0.4582568 | 0.0042767 | 0.0243435 |
| VAV2 | -0.4576977 | 0.0009869 | 0.0134835 |
| PUR2 | -0.4566343 | 0.0084588 | 0.0362101 |
| NIPBL | -0.4555292 | 0.0068132 | 0.0320676 |
| MRE11 | -0.4550863 | 0.0022383 | 0.0172874 |
| CHD4 | -0.4541916 | 0.0048021 | 0.0259733 |
| ZN207 | -0.4539649 | 0.0053765 | 0.0275992 |
| TWF1 | -0.4531293 | 0.0071789 | 0.0329686 |
| SART3 | -0.4523518 | 0.0014909 | 0.0154658 |
| RBM15 | -0.452085 | 0.0079547 | 0.0348744 |
| UN45A | -0.4515395 | 0.0044275 | 0.0247035 |
| SYLC | -0.4513279 | 0.0012503 | 0.0143009 |
| SF3B3 | -0.4492537 | 0.0027073 | 0.0187432 |
| TGT | -0.4489235 | 0.0056003 | 0.0281851 |

| | | | |
|---|---|---|---|
| FACE1 | -0.4474171 | 0.0030404 | 0.0201074 |
| FUBP2 | -0.4473402 | 0.0131646 | 0.0476999 |
| PA2G4 | -0.4472357 | 0.0034617 | 0.0218251 |
| NSUN2 | -0.4458156 | 0.009074 | 0.0378192 |
| CNO10 | -0.4458124 | 0.0138755 | 0.0495188 |
| SGPL1 | -0.4445457 | 0.0086033 | 0.0366485 |
| RS15A | -0.4440296 | 0.0090444 | 0.0377563 |
| ZN512 | -0.4439867 | 0.0098983 | 0.0401035 |
| RU17 | -0.4436058 | 0.0057295 | 0.0286422 |
| SRS10 | -0.4432044 | 0.009037 | 0.0377557 |
| T3HPD | -0.4429562 | 0.0119111 | 0.0450031 |
| RUVB1 | -0.4396788 | 0.0075238 | 0.0338764 |
| ERF1 | -0.4378435 | 0.0062907 | 0.0302583 |
| HDAC2 | -0.437429 | 0.005564 | 0.028084 |
| PP4C | -0.4371486 | 0.0131236 | 0.0475845 |
| RFC4 | -0.4370174 | 0.0113063 | 0.0437804 |
| TMX2 | -0.436352 | 0.006922 | 0.0324273 |
| RL21 | -0.4341793 | 0.0088268 | 0.0373563 |
| NC2B | -0.4311622 | 0.0122406 | 0.0456661 |
| NU107 | -0.4311508 | 0.0022547 | 0.0172874 |
| AQR | -0.4299431 | 0.0028372 | 0.0193881 |
| PRRC1 | -0.4278479 | 0.0049724 | 0.0265093 |
| TR150 | -0.4274206 | 0.0119747 | 0.0450475 |
| RL8 | -0.4271789 | 0.0079532 | 0.0348744 |
| MEMO1 | -0.4258879 | 0.0017958 | 0.0157164 |
| MSI2H | -0.424135 | 0.0098486 | 0.0400553 |
| RS27L | -0.4212249 | 0.0057068 | 0.0285832 |
| THOC6 | -0.4211355 | 0.0038924 | 0.0232206 |
| IF2A | -0.4201094 | 0.0025946 | 0.0183806 |
| COG1 | -0.4195815 | 0.0088647 | 0.0373953 |
| PAF1 | -0.4194585 | 0.0024496 | 0.0179029 |
| UGGG1 | -0.4190196 | 0.0052425 | 0.0272092 |
| CCD47 | -0.4160011 | 0.0016292 | 0.0156495 |
| TMCO1 | -0.4149507 | 0.0059241 | 0.0292225 |
| DDX3X | -0.4148589 | 0.0060167 | 0.0294432 |
| PPAL | -0.4142245 | 0.0136143 | 0.0489241 |
| EF1G | -0.4133512 | 0.0078029 | 0.0344939 |
| CPSF5 | -0.4120669 | 0.0091873 | 0.0381081 |
| PABP1 | -0.4114763 | 0.0085735 | 0.0365512 |
| PRP19 | -0.4104359 | 0.0040225 | 0.0236107 |
| PRP16 | -0.4102183 | 0.0105208 | 0.0416517 |
| IF2G | -0.4095989 | 0.0031088 | 0.0203011 |
| DDX18 | -0.4064188 | 0.0073233 | 0.0333191 |
| RED | -0.4062289 | 0.011482 | 0.0442295 |
| LUC7L | -0.4057134 | 0.0080403 | 0.0351402 |
| THOC2 | -0.4048851 | 0.0060196 | 0.0294432 |
| RSU1 | -0.4038281 | 0.0050114 | 0.0265873 |
| AKT1 | -0.4004312 | 0.0058965 | 0.0291413 |

| | | | |
|---|---|---|---|
| I2BPL | -0.4004062 | 0.0085511 | 0.0364857 |
| NCOA5 | -0.3996628 | 0.0033119 | 0.0211622 |
| BUB3 | -0.3996264 | 0.0090961 | 0.0378613 |
| DOCK7 | -0.3970453 | 0.003128 | 0.0203011 |
| TRAF6 | -0.3966313 | 0.00128 | 0.0144581 |
| IF4A3 | -0.3942233 | 0.0035029 | 0.0218931 |
| XRN2 | -0.3939978 | 0.0071403 | 0.0328884 |
| EMC8 | -0.3938177 | 0.0124777 | 0.0461408 |
| UMPS | -0.3935822 | 0.0058231 | 0.0289621 |
| SEC13 | -0.3909704 | 0.010454 | 0.0415133 |
| TMUB1 | -0.3909547 | 0.0086156 | 0.0366526 |
| SUN1 | -0.390384 | 0.0120179 | 0.0451774 |
| CHERP | -0.3891948 | 0.0042468 | 0.0243263 |
| TOIP1 | -0.3890018 | 0.0040435 | 0.0236107 |
| NLGN3 | -0.3889901 | 0.0036412 | 0.0224411 |
| SC31A | -0.3888192 | 0.0102482 | 0.040977 |
| CCAR1 | -0.388319 | 0.0040438 | 0.0236107 |
| EIF3C | -0.3882679 | 0.0021799 | 0.0170918 |
| ABCE1 | -0.3878823 | 0.0045897 | 0.0253041 |
| CLPP | -0.3873577 | 0.0103303 | 0.0412104 |
| MESD | -0.3872975 | 0.0113987 | 0.0440241 |
| ITA1 | -0.3865764 | 0.0071027 | 0.0328018 |
| HGH1 | -0.3858297 | 0.0130668 | 0.0474117 |
| RBP2 | -0.3848701 | 0.0034963 | 0.0218931 |
| VIGLN | -0.3838874 | 0.0077303 | 0.0342735 |
| CPSF2 | -0.3835252 | 0.0051426 | 0.0268674 |
| SRP14 | -0.3834307 | 0.0119665 | 0.0450475 |
| LSM6 | -0.3825388 | 0.005433 | 0.0277723 |
| ZN326 | -0.3822585 | 0.0130337 | 0.0473575 |
| AP3B1 | -0.3819282 | 0.0135882 | 0.0488952 |
| RAN | -0.3811936 | 0.008476 | 0.0362541 |
| CTNA1 | -0.3785878 | 0.0029334 | 0.0197097 |
| SPEE | -0.3776607 | 0.0090082 | 0.0376956 |
| PCBP2 | -0.3774306 | 0.0053833 | 0.0275992 |
| RASA3 | -0.3761276 | 0.0119494 | 0.0450475 |
| SPT5H | -0.3750758 | 0.0117027 | 0.0446038 |
| TP53B | -0.3742055 | 0.0106032 | 0.0419002 |
| NBN | -0.3740623 | 0.0123819 | 0.045917 |
| NU155 | -0.3730992 | 0.0068145 | 0.0320676 |
| RUFY1 | -0.3727379 | 0.0091088 | 0.0378734 |
| CDC23 | -0.3725551 | 0.0054145 | 0.0277047 |
| RL26 | -0.3723109 | 0.0055479 | 0.0280298 |
| SRP68 | -0.3698972 | 0.0025158 | 0.0180506 |
| SRPRA | -0.3697427 | 0.0103405 | 0.0412195 |
| TF3C5 | -0.3691384 | 0.0136823 | 0.0490643 |
| ASNS | -0.3671804 | 0.0079753 | 0.0349143 |
| BRE1A | -0.366038 | 0.003751 | 0.0227459 |
| COPG1 | -0.362949 | 0.0059486 | 0.0293158 |

| | | | |
|---|---|---|---|
| SMRC2 | -0.3620641 | 0.010412 | 0.041455 |
| EIF3K | -0.3617637 | 0.0082233 | 0.0356398 |
| SRP72 | -0.3607614 | 0.0083584 | 0.0358689 |
| EI2BA | -0.3606218 | 0.0107035 | 0.0421192 |
| WBP11 | -0.3599136 | 0.0091768 | 0.0381081 |
| TPR | -0.3591666 | 0.0098753 | 0.0401012 |
| SF3A1 | -0.3588446 | 0.0113862 | 0.0440087 |
| TRM2A | -0.3572003 | 0.0081661 | 0.0354738 |
| DDX42 | -0.357013 | 0.0055894 | 0.0281578 |
| SYQ | -0.3566523 | 0.007077 | 0.0327508 |
| ZC3HF | -0.3558488 | 0.0067891 | 0.0320058 |
| THEM6 | -0.3550115 | 0.014037 | 0.0499584 |
| EIF3B | -0.3526992 | 0.0096149 | 0.0393809 |
| RM02 | -0.352041 | 0.0122293 | 0.0456661 |
| RENT1 | -0.3499682 | 0.0060591 | 0.0295805 |
| GOSR1 | -0.34992 | 0.0127363 | 0.0468177 |
| DPP9 | -0.3497869 | 0.0019396 | 0.0162855 |
| NOP56 | -0.346539 | 0.0091966 | 0.0381081 |
| NUP93 | -0.3459227 | 0.0137749 | 0.0492951 |
| GCP60 | -0.3435378 | 0.0065229 | 0.0310312 |
| CDC73 | -0.3343505 | 0.0139495 | 0.0497146 |
| COPZ1 | -0.3318258 | 0.0104895 | 0.0415909 |
| EIF3M | -0.3312542 | 0.0051111 | 0.026783 |
| ASCC3 | -0.3309105 | 0.0115472 | 0.0443677 |
| NU214 | -0.3250934 | 0.0062399 | 0.030097 |
| ADAS | -0.3221652 | 0.0063038 | 0.0302653 |
| TNPO3 | -0.3187824 | 0.0116174 | 0.0445298 |
| PRS6B | -0.3183377 | 0.0127415 | 0.0468177 |
| MYG1 | -0.3138427 | 0.0122975 | 0.0457342 |
| ERF3A | -0.3130833 | 0.0107885 | 0.0423008 |
| NAA10 | -0.3104265 | 0.0051309 | 0.026833 |
| FXR1 | -0.309395 | 0.0099991 | 0.0403603 |
| AIMP2 | -0.3079093 | 0.0043043 | 0.0244209 |
| XPO2 | -0.3064319 | 0.0098013 | 0.0399561 |
| RTRAF | -0.3045045 | 0.0090322 | 0.0377557 |
| EI2BB | -0.3007752 | 0.0117965 | 0.0447975 |
| IF6 | -0.2986035 | 0.0069434 | 0.03244 |
| WDR82 | -0.2977234 | 0.0089279 | 0.0375103 |
| RBM14 | -0.2900188 | 0.0118948 | 0.0449741 |
| THOP1 | -0.2834599 | 0.0106027 | 0.0419002 |
| GORS2 | -0.279669 | 0.0116159 | 0.0445298 |
| IF4E | -0.2792018 | 0.0115723 | 0.0444314 |
| HNRH3 | -0.2707366 | 0.0129274 | 0.0470695 |
| CLPX | -0.2673626 | 0.0138604 | 0.0494988 |

Table S4

| ID | logFC | P.Value |
|---|---|---|
| GIMAP4 | 0.5044089 | 0.0273661 |
| IL10RB | 0.5085863 | 0.030423 |
| LAIR1 | 0.5682877 | 0.0406484 |
| PYCARD | 0.5888728 | 0.0398686 |
| HCK | 0.5997392 | 0.0282673 |
| IL18 | 0.6208 | 0.0399859 |
| SYK | 0.6237309 | 0.0294847 |
| PTPN6 | 0.6241661 | 0.0282126 |
| TLR1 | 0.6321153 | 0.0192317 |
| TNFRSF1B | 0.6350316 | 0.0364417 |
| TGFBR2 | 0.6369627 | 0.0405372 |
| CTLA4 | 0.6554367 | 0.0215254 |
| NFAM1 | 0.6602977 | 0.0439145 |
| ITGAL | 0.6805326 | 0.0094531 |
| CD86 | 0.7120073 | 0.0095881 |
| HAVCR2 | 0.7244646 | 0.0248183 |
| TNFSF10 | 0.7416173 | 0.0307007 |
| DDX58 | 0.7568862 | 0.020359 |
| ITGB2 | 0.7719188 | 0.0478668 |
| HLA.DMA | 0.773669 | 0.03436 |
| TLR8 | 0.7858858 | 0.0233236 |
| PDZK1IP1 | 0.7868502 | 0.0459905 |
| HLA.DMB | 0.7927193 | 0.0255272 |
| CTSS | 0.8078646 | 0.0268605 |
| CASP1 | 0.8275412 | 0.010517 |
| GPR160 | 0.8331809 | 0.0155737 |
| CSF1R | 0.8508074 | 0.0283825 |
| FGL2 | 0.8521801 | 0.0200256 |
| WAS | 0.8580479 | 0.0020537 |
| CCR5 | 0.8588631 | 0.0106676 |
| CD80 | 0.8629372 | 0.015971 |
| GPR183 | 0.8635529 | 0.0349248 |
| SAMSN1 | 0.8679975 | 0.0208867 |
| B2M | 0.8810024 | 0.021346 |
| CYBB | 0.8825685 | 0.0105903 |
| C3AR1 | 0.8835628 | 0.0107012 |
| NKG7 | 0.8843768 | 0.0267198 |
| RSAD2 | 0.8873368 | 0.0439106 |
| CXCR6 | 0.9053936 | 0.037993 |
| CYTIP | 0.9096355 | 0.0205329 |
| CCL8 | 0.9203665 | 0.0243171 |
| FPR1 | 0.9239945 | 0.0114851 |
| CTSW | 0.9385189 | 0.0040498 |
| SLC2A5 | 0.9489559 | 0.0124637 |
| XCL1.2 | 0.9614837 | 0.0093555 |
| IL7R | 0.9620619 | 0.0383844 |
| CCL5 | 0.9660931 | 0.0203326 |
| TYROBP | 0.9752938 | 0.0023041 |
| TYMP | 0.9756711 | 0.0040295 |
| ADGRE1 | 0.9832305 | 0.0265772 |
| FBLN5 | 0.9834426 | 0.0313447 |
| LST1 | 0.9988561 | 0.0007638 |
| MS4A4A | 1.0006868 | 0.0187502 |
| THBD | 1.0015344 | 0.0376398 |

| | | |
|---|---|---|
| TLR2 | 1.002133 | 0.0037113 |
| FCGR1A | 1.0124443 | 0.0022648 |
| INPP5D | 1.0208205 | 0.0010025 |
| SLAMF8 | 1.0257031 | 0.0433021 |
| NOD2 | 1.0282385 | 0.0037384 |
| CD45RO | 1.0304146 | 0.0029601 |
| LRRC25 | 1.0327653 | 0.008456 |
| SERPING1 | 1.0354428 | 0.0355236 |
| LYN | 1.064156 | 0.0043595 |
| FCGR2A | 1.0743804 | 0.0007486 |
| CD14 | 1.0802793 | 0.0148277 |
| MSR1 | 1.110607 | 0.0151285 |
| FSTL3 | 1.1160251 | 0.0448561 |
| GZMA | 1.1184342 | 0.0306436 |
| FCER1G | 1.1206983 | 0.0059288 |
| C1QA | 1.1229491 | 0.000967 |
| IL6 | 1.1313921 | 0.0499399 |
| C1QC | 1.1495641 | 0.0010692 |
| IFI30 | 1.1725648 | 0.0489163 |
| SOD2 | 1.1897733 | 0.016203 |
| VCAM1 | 1.2413426 | 0.0469498 |
| C1QB | 1.2549536 | 0.0034295 |
| LY96 | 1.2561403 | 0.0027849 |
| SRGN | 1.3014894 | 0.0004357 |
| SERPINA1 | 1.3017292 | 0.003579 |
| GBP2 | 1.3044722 | 0.0337707 |
| C3 | 1.3127558 | 0.0035238 |
| CXCL9 | 1.3228696 | 0.0234966 |
| HMOX1 | 1.343813 | 0.0235982 |
| FCGR3A.B | 1.3459608 | 0.0003187 |
| CD44 | 1.3686692 | 0.0311225 |
| SOCS3 | 1.4080474 | 0.0061013 |
| IL32 | 1.4107321 | 0.0118407 |
| TIMP1 | 1.5206082 | 0.0262893 |
| FCGR3A | 1.5523295 | 0.0028111 |
| PTX3 | 1.6012741 | 0.0366688 |
| CP | 1.6425003 | 0.0132995 |
| CD24 | 1.6797719 | 0.0439394 |
| CCL2 | 1.7090867 | 0.0278848 |
| LOX | 2.0738345 | 0.0188861 |

| ID | logFC | P.Value |
|---|---|---|
| GSTM1 | -2.7567958 | 0.0335858 |
| SPRY4 | -1.9523095 | 0.0080539 |
| ARC | -1.8329005 | 0.0168142 |
| EZH2 | -1.7470871 | 0.0010958 |
| KIF2C | -1.7018869 | 0.0021027 |
| CENPF | -1.6763021 | 0.0091351 |
| MKI67 | -1.6737077 | 0.0242195 |
| UBE2C | -1.630859 | 0.0169927 |
| TBR1 | -1.6055684 | 0.0213353 |
| BIRC5 | -1.5608495 | 0.0319259 |
| SOX11 | -1.4973613 | 0.0214865 |
| CCNB1 | -1.4669632 | 0.0057549 |
| EXO1 | -1.419259 | 0.0111814 |
| FANCA | -1.339908 | 0.0016945 |
| STAT4 | -1.3306166 | 0.0224471 |
| CEP55 | -1.3020218 | 0.0311405 |
| HELLS | -1.2834074 | 0.0006508 |
| TYMS | -1.2673052 | 0.0082717 |
| CABLES1 | -1.2625066 | 0.0068049 |
| LMNB1 | -1.1950429 | 0.048854 |
| CDC7 | -1.1922492 | 0.0015245 |
| CDK6 | -1.1908918 | 0.0248926 |
| MFGE8 | -1.1693956 | 0.0011408 |
| BRIP1 | -1.1550242 | 0.0060115 |
| HOMER1 | -1.1292206 | 0.0272443 |
| DNA2 | -1.1254197 | 0.0042395 |
| PTTG1 | -1.1245883 | 0.0311299 |
| GNG4 | -1.0802532 | 0.0338612 |
| E2F1 | -1.050145 | 0.0244116 |
| FCRLA | -1.0198717 | 0.0462708 |
| DNMT1 | -1.0196618 | 0.0014469 |
| CDC20 | -1.0069843 | 0.0060017 |
| POLE | -0.9647415 | 0.0088741 |
| SMARCA4 | -0.953758 | 0.0016636 |
| PDGFA | -0.9493618 | 0.0393402 |
| FANCC | -0.9320332 | 0.025552 |
| BLM | -0.9288167 | 0.0164753 |
| CCNE1 | -0.923651 | 0.0079874 |
| HMGB1 | -0.9057884 | 0.0019018 |
| SFXN1 | -0.8678533 | 0.0022012 |
| CDH2 | -0.8657847 | 0.0131948 |
| EPM2AIP1 | -0.8586514 | 0.0013712 |
| RAD1 | -0.8579346 | 0.0101413 |
| DNMT3A | -0.8518267 | 0.0024799 |
| H2AFX | -0.8511434 | 0.0281552 |
| TLK2 | -0.8023844 | 0.0102202 |
| UBE2T | -0.7909834 | 0.0229721 |

| | | |
|---|---|---|
| E2F3 | -0.778725 | 0.0200369 |
| PPP3R2 | -0.7707444 | 0.0024798 |
| TNKS | -0.7636874 | 0.0186843 |
| PRKACB | -0.762643 | 0.0391689 |
| CASP9 | -0.7444636 | 0.0027709 |
| SDHA | -0.728687 | 0.0365596 |
| MSH2 | -0.7282455 | 0.0391278 |
| NRDE2 | -0.726003 | 0.0037349 |
| PIK3R1 | -0.7236635 | 0.0294683 |
| SUV39H2 | -0.7184873 | 0.0271793 |
| MTOR | -0.7147229 | 0.0125542 |
| ABCF1 | -0.7139268 | 0.0034839 |
| PIAS4 | -0.7104427 | 0.011954 |
| TCF3 | -0.709624 | 0.0300128 |
| LDHB | -0.7048559 | 0.0088105 |
| CCNI | -0.6919541 | 0.0133926 |
| ZEB1 | -0.6823334 | 0.0408985 |
| ST3GAL6 | -0.6748743 | 0.0447328 |
| NEG_A.0. | -0.6712319 | 0.0489906 |
| PRKACA | -0.6683186 | 0.0120324 |
| POLD1 | -0.6588858 | 0.0305665 |
| BCL2L11 | -0.6579638 | 0.0410967 |
| AKT2 | -0.6568674 | 0.0384537 |
| SETD1B | -0.6399521 | 0.0495198 |
| IHH | -0.6279649 | 0.0079573 |
| POLR2A | -0.6197266 | 0.0431825 |
| PIK3CA | -0.6077732 | 0.0085979 |
| KDM2B | -0.6062559 | 0.0153368 |
| KRAS | -0.5960067 | 0.0224012 |
| RAD51C | -0.5935974 | 0.0376969 |
| JARID2 | -0.5855277 | 0.0399136 |
| KDM5B | -0.570039 | 0.0472239 |
| AKT1 | -0.5684867 | 0.0172318 |
| SF3A1 | -0.5663597 | 0.0130668 |
| EHMT2 | -0.5570266 | 0.0184784 |
| RPL28 | -0.5555309 | 0.0217186 |
| MRPL19 | -0.5477308 | 0.0109473 |
| RAG1 | -0.5472171 | 0.041289 |
| MLH1 | -0.544391 | 0.0102409 |
| TNFRSF4 | -0.5172147 | 0.0424536 |
| MDC1 | -0.4535408 | 0.0379141 |
| API5 | -0.4466353 | 0.0362978 |

Table S5

| ID | logFC | P.Value | adj.P.Val |
|---|---|---|---|
| VAMP1 | -1.9161321 | 0.0001355 | 0.3533576 |
| GLNA | -1.5348498 | 0.0049045 | 0.6122614 |
| SYUB | -1.4076197 | 0.002436 | 0.6122614 |
| KCNA2 | -1.3450567 | 0.0126987 | 0.6122614 |
| SYT12 | -1.3378304 | 0.0006359 | 0.5820244 |
| RAB3A | -1.3328522 | 0.0230362 | 0.6122614 |
| VA0D1 | -1.3192305 | 0.0152637 | 0.6122614 |
| CPLX1 | -1.3133944 | 0.0001319 | 0.3533576 |
| SYN3 | -1.3055736 | 0.0113582 | 0.6122614 |
| SV2B | -1.2921664 | 0.0113375 | 0.6122614 |
| SYT1 | -1.2815658 | 0.0158495 | 0.6122614 |
| TBA4A | -1.2526036 | 0.0069121 | 0.6122614 |
| RP3A | -1.22577 | 0.021855 | 0.6122614 |
| SYPH | -1.1989708 | 0.0259752 | 0.6122614 |
| VISL1 | -1.1987098 | 0.0089491 | 0.6122614 |
| STXB1 | -1.1966585 | 0.0215475 | 0.6122614 |
| VGLU1 | -1.1906492 | 0.0375579 | 0.6122614 |
| CMGA | -1.1742684 | 0.0005548 | 0.5820244 |
| SYN1 | -1.1620803 | 0.0377893 | 0.6122614 |
| DYN1 | -1.1352304 | 0.0107011 | 0.6122614 |
| CAPS2 | -1.1339775 | 0.0012688 | 0.6122614 |
| SNG3 | -1.1251984 | 0.0122918 | 0.6122614 |
| TIAM2 | -1.1237614 | 0.0005072 | 0.5820244 |
| SYUA | -1.1195769 | 0.0363447 | 0.6122614 |
| DCE2 | -1.1190196 | 0.0006698 | 0.5820244 |
| STX1B | -1.1128338 | 0.0243135 | 0.6122614 |
| GBRA1 | -1.1114093 | 0.0063515 | 0.6122614 |
| CX6B1 | -1.0987014 | 0.0065296 | 0.6122614 |
| AMPH | -1.0969017 | 0.0118911 | 0.6122614 |
| TPRGL | -1.0920845 | 0.0028837 | 0.6122614 |
| COX5A | -1.0865588 | 0.0088102 | 0.6122614 |
| SYN2 | -1.0782485 | 0.0308782 | 0.6122614 |
| ACBD7 | -1.0736255 | 0.0196254 | 0.6122614 |
| RAB3C | -1.070009 | 0.0237262 | 0.6122614 |
| KCRU | -1.0665272 | 0.0267621 | 0.6122614 |
| K0513 | -1.0619468 | 0.0193526 | 0.6122614 |
| HPLN2 | -1.0556631 | 0.0104616 | 0.6122614 |
| VAS1 | -1.0374726 | 0.0093923 | 0.6122614 |
| HXK1 | -1.0353128 | 0.0259494 | 0.6122614 |
| GNAZ | -1.0274172 | 0.0068115 | 0.6122614 |
| NDUA5 | -1.0240395 | 0.0040072 | 0.6122614 |
| ATPB | -1.023083 | 0.018307 | 0.6122614 |
| LY6H | -1.0224812 | 0.0420652 | 0.6122614 |
| PACN1 | -1.0204742 | 0.0345738 | 0.6122614 |
| NPAL3 | -1.0173247 | 0.0044673 | 0.6122614 |
| PKHA1 | -1.0095299 | 0.0192722 | 0.6122614 |
| SEPT5 | -0.999244 | 0.0211302 | 0.6122614 |
| CYC | -0.9934732 | 0.0274088 | 0.6122614 |
| THAP4 | -0.9914851 | 0.0119959 | 0.6122614 |
| CX7A2 | -0.9896704 | 0.0130712 | 0.6122614 |
| PHF24 | -0.9868375 | 0.0251154 | 0.6122614 |
| NECA1 | -0.9864228 | 0.0106458 | 0.6122614 |
| HPRT | -0.9855392 | 0.0019489 | 0.6122614 |
| SCG2 | -0.9821051 | 0.015466 | 0.6122614 |

| | | | |
|---|---|---|---|
| TBA8 | -0.9799009 | 0.0332793 | 0.6122614 |
| DUS3L | -0.978503 | 0.0140288 | 0.6122614 |
| OGDHL | -0.9784997 | 0.011362 | 0.6122614 |
| PCSK1 | -0.9673638 | 0.0217368 | 0.6122614 |
| AT2B2 | -0.9647105 | 0.0229668 | 0.6122614 |
| BSN | -0.9635437 | 0.0180365 | 0.6122614 |
| TAGL3 | -0.9633738 | 0.0197847 | 0.6122614 |
| FRIL | -0.9600694 | 0.01456 | 0.6122614 |
| AATM | -0.9576599 | 0.0114231 | 0.6122614 |
| LGI1 | -0.9524092 | 0.0033534 | 0.6122614 |
| ENOG | -0.9520815 | 0.0257372 | 0.6122614 |
| VIAAT | -0.938863 | 0.0149671 | 0.6122614 |
| S12A5 | -0.9296717 | 0.0184154 | 0.6122614 |
| SNAB | -0.9256381 | 0.0257327 | 0.6122614 |
| KCAB2 | -0.9199272 | 0.0269328 | 0.6122614 |
| VGF | -0.9188089 | 0.0134489 | 0.6122614 |
| NDUB7 | -0.9166606 | 0.0037875 | 0.6122614 |
| QCR8 | -0.9141889 | 0.0082845 | 0.6122614 |
| HCN1 | -0.9049262 | 0.0045115 | 0.6122614 |
| RYR2 | -0.8947129 | 0.0228951 | 0.6122614 |
| NFH | -0.894589 | 0.0291958 | 0.6122614 |
| K1107 | -0.8926386 | 0.0100815 | 0.6122614 |
| OXR1 | -0.8914637 | 0.0106157 | 0.6122614 |
| ATIF1 | -0.8884965 | 0.0296766 | 0.6122614 |
| NDUA4 | -0.8841476 | 0.0201468 | 0.6122614 |
| 7B2 | -0.8833169 | 0.0013886 | 0.6122614 |
| 1433G | -0.8796245 | 0.0163848 | 0.6122614 |
| KCD16 | -0.877554 | 0.0337416 | 0.6122614 |
| NDUA8 | -0.8771896 | 0.0104816 | 0.6122614 |
| NDUV2 | -0.8707485 | 0.0103057 | 0.6122614 |
| GD1L1 | -0.8668483 | 0.003242 | 0.6122614 |
| PTPRN | -0.8663113 | 0.0166623 | 0.6122614 |
| FXYD6 | -0.8662087 | 0.0180043 | 0.6122614 |
| GBRL1 | -0.863813 | 0.0401269 | 0.6122614 |
| NDUA9 | -0.8621231 | 0.0027947 | 0.6122614 |
| QCR1 | -0.8607856 | 0.0150071 | 0.6122614 |
| NDUAA | -0.8565425 | 0.0092186 | 0.6122614 |
| CMC1 | -0.8542422 | 0.0458367 | 0.6122614 |
| CLH1 | -0.8539797 | 0.0199117 | 0.6122614 |
| KCC2G | -0.8506709 | 0.0270451 | 0.6122614 |
| EPDR1 | -0.847346 | 0.0010813 | 0.6122614 |
| SH3G2 | -0.8435172 | 0.0435346 | 0.6122614 |
| AATC | -0.8424037 | 0.019873 | 0.6122614 |
| NDUS1 | -0.8403404 | 0.0125109 | 0.6122614 |
| GLSK | -0.8399315 | 0.0097404 | 0.6122614 |
| UCRI | -0.8385664 | 0.0060221 | 0.6122614 |
| GGA3 | -0.8370825 | 0.0280024 | 0.6122614 |
| CD47 | -0.8350238 | 0.0469828 | 0.6122614 |
| ST4A1 | -0.8341773 | 0.0030178 | 0.6122614 |
| KPCB | -0.832919 | 0.0279158 | 0.6122614 |
| sept6 | -0.8291398 | 0.0090937 | 0.6122614 |
| DLG4 | -0.8275773 | 0.0446438 | 0.6122614 |
| ACTN2 | -0.8256731 | 0.0132969 | 0.6122614 |
| FUMH | -0.8249704 | 0.0092987 | 0.6122614 |
| LEGL | -0.8244628 | 0.0048204 | 0.6122614 |
| NDUB4 | -0.8204543 | 0.0292152 | 0.6122614 |

| | | | |
|---|---|---|---|
| CPNE5 | -0.8173477 | 0.0022357 | 0.6122614 |
| KKCC2 | -0.8120307 | 0.0076809 | 0.6122614 |
| RPGF4 | -0.8119029 | 0.0498726 | 0.6122614 |
| ACPM | -0.8001031 | 0.0052113 | 0.6122614 |
| CISD1 | -0.7955468 | 0.0303367 | 0.6122614 |
| GUAD | -0.7951154 | 0.018414 | 0.6122614 |
| GBB1 | -0.7936719 | 0.0306106 | 0.6122614 |
| ABLM2 | -0.7923969 | 0.0204838 | 0.6122614 |
| ODO1 | -0.7904368 | 0.006101 | 0.6122614 |
| I5P1 | -0.7893685 | 0.0056263 | 0.6122614 |
| GNAI1 | -0.7882277 | 0.0384312 | 0.6122614 |
| LGI3 | -0.7859722 | 0.0317256 | 0.6122614 |
| IDH3G | -0.7823884 | 0.0159961 | 0.6122614 |
| QCR9 | -0.7810156 | 0.0087551 | 0.6122614 |
| NDUAB | -0.7807068 | 0.028651 | 0.6122614 |
| NDUS8 | -0.7795864 | 0.0329528 | 0.6122614 |
| GLE1 | -0.7777953 | 0.0040218 | 0.6122614 |
| ADA23 | -0.7775552 | 0.0334648 | 0.6122614 |
| EF1A2 | -0.7773957 | 0.0440877 | 0.6122614 |
| AP2A1 | -0.7747111 | 0.0286505 | 0.6122614 |
| AP2A2 | -0.7747004 | 0.0079508 | 0.6122614 |
| NDUS5 | -0.7734534 | 0.0457211 | 0.6122614 |
| DYN3 | -0.7718463 | 0.0273364 | 0.6122614 |
| PDE1B | -0.7708695 | 0.0274956 | 0.6122614 |
| SUCB1 | -0.7705371 | 0.0156633 | 0.6122614 |
| NDUA1 | -0.7693221 | 0.0146789 | 0.6122614 |
| NDUV1 | -0.763485 | 0.0045038 | 0.6122614 |
| DYN2 | -0.7610634 | 0.0280445 | 0.6122614 |
| VATA | -0.7588058 | 0.0321671 | 0.6122614 |
| VAMP2 | -0.7585973 | 0.0470989 | 0.6122614 |
| TBB4A | -0.7551989 | 0.0179525 | 0.6122614 |
| RGS6 | -0.7521182 | 0.031218 | 0.6122614 |
| AP180 | -0.7511568 | 0.0243744 | 0.6122614 |
| CCKN | -0.7508839 | 0.0489005 | 0.6122614 |
| RIMS2 | -0.7488088 | 0.0481678 | 0.6122614 |
| AJM1 | -0.7455625 | 0.044196 | 0.6122614 |
| PPM1H | -0.7432318 | 0.0172805 | 0.6122614 |
| ATP5I | -0.7412644 | 0.016569 | 0.6122614 |
| MPC1 | -0.7407459 | 0.0418142 | 0.6122614 |
| AUXI | -0.7389502 | 0.0482582 | 0.6122614 |
| ATPA | -0.7388288 | 0.0440105 | 0.6122614 |
| RDH13 | -0.7351633 | 0.0218718 | 0.6122614 |
| PLD3 | -0.7308894 | 0.0165498 | 0.6122614 |
| KAP3 | -0.7254717 | 0.0213686 | 0.6122614 |
| CRAC1 | -0.7253764 | 0.0040183 | 0.6122614 |
| AP2M1 | -0.7253545 | 0.0128131 | 0.6122614 |
| DLGP1 | -0.7226114 | 0.045033 | 0.6122614 |
| CAD13 | -0.7208076 | 0.0204255 | 0.6122614 |
| PXL2A | -0.7205082 | 0.0062723 | 0.6122614 |
| IGS21 | -0.7204373 | 0.0197759 | 0.6122614 |
| RAP2A | -0.7202192 | 0.0163878 | 0.6122614 |
| AP2S1 | -0.7200956 | 0.0054853 | 0.6122614 |
| SYT7 | -0.7197373 | 0.0476746 | 0.6122614 |
| MDHM | -0.7183405 | 0.0471594 | 0.6122614 |
| MYO5A | -0.7182343 | 0.0330457 | 0.6122614 |
| NDRG4 | -0.7059248 | 0.0376548 | 0.6122614 |

| | | | |
|---|---|---|---|
| ATP5H | -0.7058952 | 0.039646 | 0.6122614 |
| CAPS1 | -0.7051419 | 0.0497157 | 0.6122614 |
| MAP6 | -0.7023462 | 0.029521 | 0.6122614 |
| DHPR | -0.7021934 | 0.0186029 | 0.6122614 |
| E41L3 | -0.7020182 | 0.0358412 | 0.6122614 |
| AP2B1 | -0.7013193 | 0.0202801 | 0.6122614 |
| IDH3A | -0.7010477 | 0.0349817 | 0.6122614 |
| SL9A1 | -0.7000809 | 0.0499765 | 0.6122614 |
| NDUS6 | -0.6981554 | 0.0386768 | 0.6122614 |
| SHLB2 | -0.6979082 | 0.0166022 | 0.6122614 |
| GNB5 | -0.697905 | 0.0177789 | 0.6122614 |
| LIPA3 | -0.6966972 | 0.0183209 | 0.6122614 |
| PCLO | -0.6964695 | 0.0417679 | 0.6122614 |
| AAK1 | -0.6908652 | 0.0082165 | 0.6122614 |
| ATPG | -0.6908131 | 0.0435214 | 0.6122614 |
| STXB5 | -0.6906736 | 0.0266551 | 0.6122614 |
| TPD53 | -0.6906432 | 0.0396341 | 0.6122614 |
| PURA | -0.6906139 | 0.020728 | 0.6122614 |
| PTN5 | -0.6886443 | 0.0104156 | 0.6122614 |
| HOME1 | -0.6859908 | 0.0321122 | 0.6122614 |
| VATE1 | -0.6813383 | 0.0451256 | 0.6122614 |
| NDUC2 | -0.6812958 | 0.0310374 | 0.6122614 |
| FRIH | -0.6812941 | 0.0457718 | 0.6122614 |
| GSTM3 | -0.6812858 | 0.0292589 | 0.6122614 |
| VATB2 | -0.6808115 | 0.0420132 | 0.6122614 |
| QCR2 | -0.6785109 | 0.0201836 | 0.6122614 |
| NDUB3 | -0.6769103 | 0.0284197 | 0.6122614 |
| PTPR2 | -0.6752381 | 0.0461001 | 0.6122614 |
| SHAN1 | -0.6745912 | 0.0334565 | 0.6122614 |
| QCR7 | -0.6710879 | 0.0386606 | 0.6122614 |
| NEC1 | -0.6689533 | 0.0253405 | 0.6122614 |
| NDUA2 | -0.6640488 | 0.0331815 | 0.6122614 |
| CNTP2 | -0.6619207 | 0.0312862 | 0.6122614 |
| IDH3B | -0.661597 | 0.0387816 | 0.6122614 |
| ODPA | -0.6605505 | 0.0393826 | 0.6122614 |
| HCN2 | -0.6590775 | 0.017244 | 0.6122614 |
| MADD | -0.657436 | 0.0365501 | 0.6122614 |
| NDUS2 | -0.6549938 | 0.0159021 | 0.6122614 |
| PAK1 | -0.6538767 | 0.0190005 | 0.6122614 |
| NDUBA | -0.6517847 | 0.0358393 | 0.6122614 |
| NDUS3 | -0.6508651 | 0.0396027 | 0.6122614 |
| SCAI | -0.6503144 | 0.021089 | 0.6122614 |
| NDUS4 | -0.647231 | 0.0104505 | 0.6122614 |
| MGLL | -0.638165 | 0.0361834 | 0.6122614 |
| CPEB3 | -0.6371274 | 0.0157677 | 0.6122614 |
| ADA22 | -0.6319171 | 0.021302 | 0.6122614 |
| FKBP8 | -0.6298978 | 0.0124916 | 0.6122614 |
| ATPK | -0.6291291 | 0.0280955 | 0.6122614 |
| SDHA | -0.6271181 | 0.0289529 | 0.6122614 |
| NDUA6 | -0.624359 | 0.0182342 | 0.6122614 |
| CTRO | -0.6240137 | 0.0201854 | 0.6122614 |
| VATC1 | -0.6238751 | 0.0470364 | 0.6122614 |
| PI4KA | -0.6233475 | 0.013773 | 0.6122614 |
| NDUAC | -0.6211741 | 0.0303028 | 0.6122614 |
| SYNPO | -0.6200269 | 0.0454536 | 0.6122614 |
| CA2D2 | -0.6174285 | 0.0263911 | 0.6122614 |

| | | | |
|---|---|---|---|
| FAHD1 | -0.6170401 | 0.0268547 | 0.6122614 |
| CA2D3 | -0.6157551 | 0.03449 | 0.6122614 |
| BDH | -0.6154002 | 0.0155298 | 0.6122614 |
| SYFA | -0.6147337 | 0.0377062 | 0.6122614 |
| DNM1L | -0.6102965 | 0.0435199 | 0.6122614 |
| ATLA1 | -0.6101131 | 0.0202956 | 0.6122614 |
| CNTP1 | -0.6058874 | 0.0420257 | 0.6122614 |
| ARFG1 | -0.6043776 | 0.0165888 | 0.6122614 |
| IQEC1 | -0.6030945 | 0.0363297 | 0.6122614 |
| IP3KA | -0.6006285 | 0.0297095 | 0.6122614 |
| ECHM | -0.597867 | 0.030329 | 0.6122614 |
| CTBP1 | -0.5958787 | 0.0142205 | 0.6122614 |
| TINAL | -0.5950654 | 0.0124353 | 0.6122614 |
| MTX2 | -0.5948404 | 0.0048272 | 0.6122614 |
| MIC26 | -0.5935517 | 0.0233229 | 0.6122614 |
| RT36 | -0.5934752 | 0.0218745 | 0.6122614 |
| AP3B2 | -0.5931657 | 0.0397128 | 0.6122614 |
| ODPB | -0.5919448 | 0.0473675 | 0.6122614 |
| MTMR2 | -0.5915321 | 0.0139054 | 0.6122614 |
| NDUAD | -0.5894765 | 0.0352744 | 0.6122614 |
| COX5B | -0.5875847 | 0.0425144 | 0.6122614 |
| ATPD | -0.5873988 | 0.0279935 | 0.6122614 |
| DCE1 | -0.5795727 | 0.0139942 | 0.6122614 |
| F1712 | -0.5774141 | 0.0248347 | 0.6122614 |
| BRSK2 | -0.5760693 | 0.0417148 | 0.6122614 |
| AP1B1 | -0.5756104 | 0.0056516 | 0.6122614 |
| NGEF | -0.5737125 | 0.0317604 | 0.6122614 |
| DGLA | -0.5727657 | 0.0307244 | 0.6122614 |
| PEX5R | -0.5652537 | 0.0396703 | 0.6122614 |
| ARBK1 | -0.5638927 | 0.0210875 | 0.6122614 |
| RPGF2 | -0.5637727 | 0.0279488 | 0.6122614 |
| CISY | -0.5615277 | 0.0419789 | 0.6122614 |
| MTURN | -0.5590395 | 0.0386158 | 0.6122614 |
| NRX1A | -0.5584255 | 0.042672 | 0.6122614 |
| SRCN1 | -0.5569017 | 0.0298772 | 0.6122614 |
| ODP2 | -0.5542997 | 0.0364885 | 0.6122614 |
| WDR47 | -0.5520669 | 0.0260497 | 0.6122614 |
| ARP3B | -0.5483109 | 0.0418312 | 0.6122614 |
| SDHB | -0.5475362 | 0.0181417 | 0.6122614 |
| REEP2 | -0.5473889 | 0.0318676 | 0.6122614 |
| GAS7 | -0.5470009 | 0.035871 | 0.6122614 |
| NDUB9 | -0.545824 | 0.0164525 | 0.6122614 |
| NFU1 | -0.5445816 | 0.012459 | 0.6122614 |
| ODO2 | -0.5395322 | 0.037023 | 0.6122614 |
| DNJB2 | -0.5392933 | 0.0465215 | 0.6122614 |
| DCD | -0.5374464 | 0.0483786 | 0.6122614 |
| NECP1 | -0.5370394 | 0.0261576 | 0.6122614 |
| WDR7 | -0.5355072 | 0.0168145 | 0.6122614 |
| TIP | -0.5337287 | 0.0494136 | 0.6122614 |
| VATF | -0.5320111 | 0.0429683 | 0.6122614 |
| AUHM | -0.531178 | 0.0271907 | 0.6122614 |
| AP3D1 | -0.5311234 | 0.0231198 | 0.6122614 |
| RAB2B | -0.5302046 | 0.0197791 | 0.6122614 |
| ATAD1 | -0.5295665 | 0.0210006 | 0.6122614 |
| RPGP1 | -0.5294819 | 0.0486294 | 0.6122614 |
| SKT | -0.5226273 | 0.0357186 | 0.6122614 |

| | | | |
|---|---|---|---|
| GGT7 | -0.522259 | 0.0425175 | 0.6122614 |
| NRBP | -0.520111 | 0.0364699 | 0.6122614 |
| 2A5E | -0.5159614 | 0.009232 | 0.6122614 |
| BIG2 | -0.5153025 | 0.0141721 | 0.6122614 |
| 2A5G | -0.5138808 | 0.0303081 | 0.6122614 |
| WDR37 | -0.511559 | 0.0433414 | 0.6122614 |
| PACS1 | -0.5048023 | 0.0350769 | 0.6122614 |
| STML2 | -0.5035667 | 0.0153854 | 0.6122614 |
| OCRL | -0.5027713 | 0.0072472 | 0.6122614 |
| TBC24 | -0.5019654 | 0.033577 | 0.6122614 |
| NMT2 | -0.4981883 | 0.0372455 | 0.6122614 |
| OSB10 | -0.4969316 | 0.0373561 | 0.6122614 |
| CSN8 | -0.4951164 | 0.0075627 | 0.6122614 |
| MTMR5 | -0.492938 | 0.0390187 | 0.6122614 |
| TIM9 | -0.4910341 | 0.0426388 | 0.6122614 |
| S27A4 | -0.4873104 | 0.021083 | 0.6122614 |
| ISCU | -0.478332 | 0.0427371 | 0.6122614 |
| PP1G | -0.4752402 | 0.0478734 | 0.6122614 |
| LIGO1 | -0.4750654 | 0.0492963 | 0.6122614 |
| TLN2 | -0.4674803 | 0.0273061 | 0.6122614 |
| TMX3 | -0.4672181 | 0.0204848 | 0.6122614 |
| IF2M | -0.4656495 | 0.0305607 | 0.6122614 |
| RETR2 | -0.4628199 | 0.0286241 | 0.6122614 |
| RFIP5 | -0.4605523 | 0.0422077 | 0.6122614 |
| PDZD8 | -0.4593938 | 0.0307031 | 0.6122614 |
| SPN90 | -0.4556661 | 0.0476963 | 0.6122614 |
| KLC2 | -0.4526125 | 0.0190092 | 0.6122614 |
| ATPF1 | -0.4503601 | 0.0270644 | 0.6122614 |
| PPCEL | -0.4450179 | 0.0412109 | 0.6122614 |
| PDPK1 | -0.4435168 | 0.0211468 | 0.6122614 |
| AGK | -0.4428372 | 0.0285164 | 0.6122614 |
| HINT3 | -0.4380346 | 0.044944 | 0.6122614 |
| FIBP | -0.4374665 | 0.0440131 | 0.6122614 |
| ACSL6 | -0.4335556 | 0.0490894 | 0.6122614 |
| ACYP1 | -0.4200796 | 0.0443514 | 0.6122614 |
| T22D3 | -0.4102447 | 0.0453333 | 0.6122614 |
| HD | -0.4097 | 0.0162159 | 0.6122614 |
| SYSM | -0.4093563 | 0.0429659 | 0.6122614 |
| CACO1 | -0.4085221 | 0.0313094 | 0.6122614 |
| KAD3 | -0.4068561 | 0.0368049 | 0.6122614 |
| EFHD2 | -0.4044802 | 0.029991 | 0.6122614 |
| KAPCB | -0.403785 | 0.0398715 | 0.6122614 |
| ACOT9 | -0.4010187 | 0.0318571 | 0.6122614 |
| SRBS2 | -0.4001903 | 0.0434565 | 0.6122614 |
| LONM | -0.3983875 | 0.0322512 | 0.6122614 |
| ASAP2 | -0.3970362 | 0.037684 | 0.6122614 |
| TPC10 | -0.3929641 | 0.0248649 | 0.6122614 |
| UBE2O | -0.3908356 | 0.0298568 | 0.6122614 |
| PPME1 | -0.3892235 | 0.026393 | 0.6122614 |
| ARFP2 | -0.3891169 | 0.0373456 | 0.6122614 |
| ABHDA | -0.3861107 | 0.0266389 | 0.6122614 |
| HEMH | -0.3855628 | 0.0488916 | 0.6122614 |
| APLP2 | -0.3850029 | 0.0288359 | 0.6122614 |
| DCTN5 | -0.3811458 | 0.0382848 | 0.6122614 |
| RAE1 | -0.3760279 | 0.0484502 | 0.6122614 |
| MYCB2 | -0.3744373 | 0.0429979 | 0.6122614 |

| | | | |
|---|---|---|---|
| VPS35 | -0.3679952 | 0.0459013 | 0.6122614 |
| NFS1 | -0.3645402 | 0.0309009 | 0.6122614 |
| DEST | -0.3499986 | 0.0358002 | 0.6122614 |
| WDR13 | -0.3317436 | 0.047172 | 0.6122614 |
| WDR48 | -0.3317063 | 0.0458369 | 0.6122614 |
| DCTN6 | -0.322857 | 0.0475871 | 0.6122614 |
| CUL1 | -0.2923196 | 0.0482253 | 0.6122614 |

| ID | logFC | P.Value | adj.P.Val |
|---|---|---|---|
| SMAD4 | 2.1589213 | 0.0231002 | 0.6122614 |
| GCP6 | 1.6892211 | 0.0034596 | 0.6122614 |
| CAH3 | 1.6423506 | 0.0065237 | 0.6122614 |
| NCOR1 | 1.5971358 | 0.0013721 | 0.6122614 |
| MK | 1.4111774 | 0.0418228 | 0.6122614 |
| PGAM2 | 1.3019285 | 0.0309045 | 0.6122614 |
| APOBR | 1.1234251 | 0.0290503 | 0.6122614 |
| GBP1 | 1.1212286 | 0.0273091 | 0.6122614 |
| FLNC | 1.0852877 | 0.044621 | 0.6122614 |
| PDLI3 | 1.0232883 | 0.0353063 | 0.6122614 |
| DEFI6 | 0.9989998 | 0.0229554 | 0.6122614 |
| CO5A3 | 0.9838497 | 0.0328326 | 0.6122614 |
| HMGN2 | 0.9795392 | 0.0348595 | 0.6122614 |
| LV545;LV539 | 0.9699828 | 0.009455 | 0.6122614 |
| VAMP8 | 0.9620738 | 0.0433136 | 0.6122614 |
| ADSV | 0.9587892 | 0.0379239 | 0.6122614 |
| XIAP | 0.9462831 | 0.0269072 | 0.6122614 |
| IFIT2 | 0.9400532 | 0.035067 | 0.6122614 |
| AMPL | 0.9292757 | 0.040085 | 0.6122614 |
| PSME2 | 0.921252 | 0.0179644 | 0.6122614 |
| PSME1 | 0.9120681 | 0.030724 | 0.6122614 |
| MRC2 | 0.8687587 | 0.0339626 | 0.6122614 |
| GILT | 0.8687079 | 0.0106549 | 0.6122614 |
| RNF14 | 0.8412913 | 0.0201532 | 0.6122614 |
| RNH2B | 0.8149627 | 0.0472827 | 0.6122614 |
| C1QA | 0.7928702 | 0.0098674 | 0.6122614 |
| THAS | 0.7889978 | 0.042286 | 0.6122614 |
| MYD88 | 0.7797336 | 0.0178273 | 0.6122614 |
| AL1L2 | 0.7700861 | 0.0322181 | 0.6122614 |
| NASP | 0.7595471 | 0.0410561 | 0.6122614 |
| LUM | 0.757482 | 0.0449706 | 0.6122614 |
| AP5Z1 | 0.755764 | 0.0026003 | 0.6122614 |
| ATX1 | 0.7539799 | 0.0083468 | 0.6122614 |
| WDR12 | 0.752143 | 0.0117692 | 0.6122614 |
| DTX3L | 0.7486202 | 0.0104751 | 0.6122614 |
| MED22 | 0.7309918 | 0.049064 | 0.6122614 |
| DNJC2 | 0.7186368 | 0.0025756 | 0.6122614 |
| FCGR1;FCG | 0.6862104 | 0.0100865 | 0.6122614 |
| MARCS | 0.679098 | 0.0471201 | 0.6122614 |
| NEK9 | 0.6747374 | 0.0234239 | 0.6122614 |
| RUXF | 0.6506346 | 0.0442924 | 0.6122614 |
| PHF5A | 0.6417201 | 0.0173081 | 0.6122614 |
| LIPB1 | 0.6299563 | 0.0243074 | 0.6122614 |
| BUD31 | 0.6164846 | 0.0454187 | 0.6122614 |
| RBM6 | 0.613181 | 0.037194 | 0.6122614 |
| ZPR1 | 0.6074285 | 0.0078147 | 0.6122614 |
| PYGL | 0.607311 | 0.0350241 | 0.6122614 |

| | | | |
|---|---|---|---|
| NCKPL | 0.6025029 | 0.014654 | 0.6122614 |
| NRP1 | 0.5524801 | 0.0422143 | 0.6122614 |
| TMM43 | 0.5516545 | 0.0325396 | 0.6122614 |
| MET16 | 0.5402562 | 0.0316722 | 0.6122614 |
| RFC3 | 0.5401557 | 0.0476257 | 0.6122614 |
| ZC3HD | 0.5226054 | 0.047481 | 0.6122614 |
| ELP5 | 0.5074665 | 0.0385747 | 0.6122614 |
| PP4C | 0.4895949 | 0.0476586 | 0.6122614 |
| RFC1 | 0.4883453 | 0.0400275 | 0.6122614 |
| TARA | 0.4871502 | 0.0220461 | 0.6122614 |
| GOGA1 | 0.4830151 | 0.0393932 | 0.6122614 |
| SLIK2 | 0.48205 | 0.0403838 | 0.6122614 |
| TANC1 | 0.4798445 | 0.0310182 | 0.6122614 |
| NHP2 | 0.4782759 | 0.0466373 | 0.6122614 |
| RO60 | 0.4742059 | 0.0360904 | 0.6122614 |
| UBE2H | 0.4719062 | 0.0458377 | 0.6122614 |
| ANS1A | 0.4714103 | 0.0340775 | 0.6122614 |
| EVI5 | 0.4685348 | 0.0381365 | 0.6122614 |
| THOC6 | 0.4669189 | 0.0213874 | 0.6122614 |
| APEX1 | 0.4607766 | 0.0493364 | 0.6122614 |
| HEM3 | 0.4545916 | 0.0261224 | 0.6122614 |
| ZC3H4 | 0.4534148 | 0.0397642 | 0.6122614 |
| RNZ2 | 0.4426157 | 0.0279268 | 0.6122614 |
| AQR | 0.4388979 | 0.0223833 | 0.6122614 |
| PHAG1 | 0.4356019 | 0.0227067 | 0.6122614 |
| STEA3 | 0.4320354 | 0.0365992 | 0.6122614 |
| PP1R8 | 0.4160211 | 0.019005 | 0.6122614 |
| RECQ1 | 0.3979276 | 0.0429105 | 0.6122614 |
| TGT | 0.3854563 | 0.0499481 | 0.6122614 |
| H1BP3 | 0.3839421 | 0.0400897 | 0.6122614 |
| SELB | 0.3835108 | 0.0485005 | 0.6122614 |
| FA84B | 0.3809678 | 0.0337423 | 0.6122614 |
| GCN1 | 0.3714801 | 0.0400348 | 0.6122614 |
| WBP11 | 0.3692747 | 0.0476057 | 0.6122614 |
| KI13B | 0.35866 | 0.04795 | 0.6122614 |

Table S6

| OLR1 | logFC | P.Value |
|---|---|---|
| APOC2 | -2.3845603 | 0.0000134 |
| SPP1 | -2.2097648 | 0.0147855 |
| OLR1 | -1.6849744 | 0.0003142 |
| PADI2 | -1.6683084 | 0.0000883 |
| HK2 | -1.6444173 | 0.0000926 |
| PLIN2 | -1.6052967 | 0.0041471 |
| NCF1 | -1.5773076 | 0.0002337 |
| HMOX1 | -1.5668096 | 0.0011128 |
| GFAP | -1.5072616 | 0.0422588 |
| MT1X | -1.5060106 | 0.0013016 |
| RGCC | -1.4210037 | 0.0004821 |
| APOC1 | -1.4205663 | 0.0104208 |
| ALOX5AP | -1.3714169 | 0.0017791 |
| MT1G | -1.3319214 | 0.0188823 |
| SLC16A3 | -1.310631 | 0.0008964 |
| MT3 | -1.2529458 | 0.0374735 |
| IGFBP5 | -1.2464083 | 0.0053746 |
| HILPDA | -1.1901802 | 0.0103319 |
| PLAUR | -1.1725798 | 0.0269894 |
| SLC2A5 | -1.1661916 | 0.0035983 |
| RAB42 | -1.1475586 | 0.0048088 |
| ICOSLG | -1.1318137 | 0.0001203 |
| MTHFD2 | -1.1271699 | 0.0013553 |
| ATG16L2 | -1.1250691 | 0.0008513 |
| SLC11A1 | -1.1206672 | 0.0079837 |
| RNASET2 | -1.1173631 | 0.0010836 |
| DNER | -1.1154985 | 0.000464 |
| RGS10 | -1.0976283 | 0.0034341 |
| NPC2 | -1.0827763 | 0.0162408 |
| DENND3 | -1.0685221 | 0.0000802 |
| DDIT4 | -1.061467 | 0.001994 |
| ITGB2 | -1.0560877 | 0.0019709 |
| ITGAX | -1.0534995 | 0.0006653 |
| TGFB1 | -1.0516798 | 0.0004673 |
| SOCS6 | -1.0484991 | 0.0004168 |
| RNASE4 | -1.0364395 | 0.0036471 |
| PFKFB3 | -1.0326743 | 0.0053175 |
| NUPR1 | -1.0267837 | 0.021969 |
| HPCAL1 | -1.0228657 | 0.0013528 |
| CTNND1 | -1.0219494 | 0.0005938 |
| RHOG | -1.0184139 | 0.0007475 |
| ADSS1 | -1.0165344 | 0.0089346 |
| MT1M | -1.0091213 | 0.0054879 |
| ALKBH5 | -1.008012 | 0.0001286 |
| TYROBP | -1.0065049 | 0.0137569 |
| PSD4 | -1.0061361 | 0.0053471 |
| IRF8 | -1.0047139 | 0.0030993 |
| MT2A | -1.0024551 | 0.0142252 |
| BIN1 | -0.9985446 | 0.0031469 |
| COTL1 | -0.9841151 | 0.0068076 |
| PHB2 | -0.9769166 | 0.0049455 |
| PLD4 | -0.9677272 | 0.002625 |
| GLRX | -0.9666499 | 0.0024403 |
| PIM1 | -0.962645 | 0.0052912 |

| | | |
|---|---|---|
| CEBPG | -0.957899 | 0.0003624 |
| CYFIP1 | -0.9568966 | 0.0038013 |
| BCL6 | -0.9544142 | 0.0055472 |
| CCND2 | -0.9529942 | 0.0198036 |
| SLC2A1 | -0.9509477 | 0.0026626 |
| CXCL16 | -0.9400505 | 0.0014174 |
| FYB1 | -0.9357266 | 0.0048722 |
| RNF181 | -0.9353913 | 0.0065982 |
| PLXDC2 | -0.9322933 | 0.0100744 |
| CENPB | -0.9316205 | 0.001058 |
| GPX1 | -0.9289003 | 0.011985 |
| MAFF | -0.9271049 | 0.0049632 |
| TBXAS1 | -0.9255751 | 0.0012937 |
| GPI | -0.9218765 | 0.0096452 |
| CYTH4 | -0.9213451 | 0.0499447 |
| MACROH2A | -0.9186783 | 0.001446 |
| PPAN | -0.9186435 | 0.0026986 |
| ARPC1B | -0.9149385 | 0.0120593 |
| CARHSP1 | -0.9124985 | 0.0085896 |
| APBB1IP | -0.9055515 | 0.0017706 |
| PTCRA | -0.8914738 | 0.0049831 |
| HCLS1 | -0.8906904 | 0.0269706 |
| VKORC1 | -0.8873102 | 0.001151 |
| DNASE2 | -0.8819009 | 0.0192062 |
| ZBED5 | -0.8818614 | 0.00566 |
| IL13RA1 | -0.8818432 | 0.0040557 |
| P4HA1 | -0.8784004 | 0.0101472 |
| SAMSN1 | -0.8746155 | 0.0087292 |
| MGAT1 | -0.8736309 | 0.0049951 |
| HCN2 | -0.8715731 | 0.0169262 |
| PLEKHO1 | -0.8685103 | 0.0076856 |
| TP53I3 | -0.8672279 | 0.0013231 |
| ERO1A | -0.8622256 | 0.0242632 |
| AP1B1 | -0.8617247 | 0.0087468 |
| ABHD4 | -0.861483 | 0.0022964 |
| TREM2 | -0.8610157 | 0.0184171 |
| MMP24OS | -0.8589075 | 0.0014896 |
| ADGRG1 | -0.8585016 | 0.0123845 |
| H2AC16 | -0.856743 | 0.0118756 |
| UBE2A | -0.8561661 | 0.0015531 |
| LHFPL2 | -0.8540972 | 0.0035287 |
| PFKP | -0.8540491 | 0.0108785 |
| PPT1 | -0.8522629 | 0.0100936 |
| TTC7A | -0.8517416 | 0.0019552 |
| CERK | -0.8511899 | 0.002819 |
| SYNGR2 | -0.8504463 | 0.0034953 |
| LILRB4 | -0.848657 | 0.0159041 |
| SLC25A37 | -0.8482243 | 0.009582 |
| H2BC7 | -0.8460161 | 0.0058982 |
| TCEAL4 | -0.8457746 | 0.0006478 |
| DDX41 | -0.8457595 | 0.0025802 |
| VPS54 | -0.8431637 | 0.0263271 |
| FOXG1 | -0.8427578 | 0.0277631 |
| SHMT2 | -0.8407196 | 0.0016082 |
| BCAT1 | -0.8352603 | 0.0085116 |
| SLC15A3 | -0.8342869 | 0.0126357 |

| | | |
|---|---|---|
| ARHGEF12 | -0.8308973 | 0.0447635 |
| CEBPA | -0.8205609 | 0.020744 |
| MTSS2 | -0.8201827 | 0.0038788 |
| IRF2BP2 | -0.8178837 | 0.0094775 |
| RPS6KA1 | -0.8126501 | 0.0121458 |
| ETS2 | -0.8121557 | 0.0007406 |
| CEBPB | -0.8115962 | 0.0066784 |
| EIF4A1 | -0.8060435 | 0.0036334 |
| INPP5D | -0.803216 | 0.007554 |
| ZNF703 | -0.8028933 | 0.0089395 |
| EHD2 | -0.8024351 | 0.0165004 |
| FERMT3 | -0.802255 | 0.0052983 |
| BHLHE41 | -0.8013259 | 0.0108965 |
| GPN3 | -0.7974069 | 0.025561 |
| WDR81 | -0.7967994 | 0.028711 |
| H4C2 | -0.7956803 | 0.0072083 |
| LPCAT2 | -0.795188 | 0.0107946 |
| COL4A1 | -0.7950809 | 0.0049311 |
| SCAF1 | -0.7942337 | 0.0038528 |
| RNF166 | -0.7937168 | 0.0082039 |
| INF2 | -0.7930928 | 0.0122765 |
| NUDCD2 | -0.7918649 | 0.0111867 |
| CD37 | -0.7911304 | 0.0323498 |
| PPP1R3C | -0.7906476 | 0.0162806 |
| DCXR | -0.7887786 | 0.0093494 |
| MRPS23 | -0.7840253 | 0.009941 |
| ORMDL3 | -0.7806801 | 0.0252116 |
| PFKM | -0.7801318 | 0.0235223 |
| SPI1 | -0.7796197 | 0.018808 |
| SSPN | -0.7786809 | 0.0196563 |
| CSTB | -0.7774382 | 0.0496062 |
| CDKN1A | -0.77499 | 0.0365976 |
| SLC25A19 | -0.7747488 | 0.0105149 |
| MOB1A | -0.7744756 | 0.0022778 |
| CD300A | -0.7726407 | 0.0138511 |
| UNC93B1 | -0.7718996 | 0.0208002 |
| CD9 | -0.7700271 | 0.0331823 |
| FOXN3 | -0.7691473 | 0.0126194 |
| COL4A2 | -0.7679592 | 0.0057585 |
| TCIRG1 | -0.7669857 | 0.0410172 |
| ZDHHC14 | -0.7668103 | 0.037495 |
| SMPD4 | -0.7658016 | 0.0012905 |
| TNFRSF12A | -0.7644959 | 0.0362209 |
| CD53 | -0.7643035 | 0.0207902 |
| PRKAR2A | -0.7625915 | 0.037973 |
| DDB2 | -0.7608561 | 0.0019556 |
| PLAC8 | -0.7553535 | 0.0045227 |
| HLA-DMA | -0.7538579 | 0.0265455 |
| RNASE6 | -0.7479745 | 0.0143518 |
| ANGPTL4 | -0.7472219 | 0.0174477 |
| CLN8 | -0.7413647 | 0.0044074 |
| CD83 | -0.7404914 | 0.0117325 |
| HAVCR2 | -0.7399377 | 0.0036794 |
| SLC7A5 | -0.7374245 | 0.0102845 |
| PTGS1 | -0.7373426 | 0.0223006 |
| HNRNPK | -0.7369588 | 0.0082089 |

| | | |
|---|---|---|
| PLK3 | -0.7363665 | 0.0106339 |
| FMNL3 | -0.7357711 | 0.0131208 |
| SH3TC1 | -0.7352347 | 0.0071469 |
| SAP30 | -0.732604 | 0.0075389 |
| ERCC1 | -0.7301218 | 0.0034793 |
| TSPO | -0.7299505 | 0.0029255 |
| KDM2B | -0.7292148 | 0.0068948 |
| SHC2 | -0.7285286 | 0.0095286 |
| UBA52 | -0.7269362 | 0.0348858 |
| MLC1 | -0.7266099 | 0.0142871 |
| PGAP1 | -0.7242122 | 0.0406637 |
| KCNJ5 | -0.7238933 | 0.0102382 |
| PALD1 | -0.7236806 | 0.0473503 |
| EIF3D | -0.7223818 | 0.0075512 |
| AXL | -0.7207556 | 0.0042514 |
| CAPG | -0.7189921 | 0.0214602 |
| SEPTIN9 | -0.7179011 | 0.0245351 |
| NFE2L2 | -0.7162337 | 0.0120956 |
| NRXN2 | -0.7161213 | 0.016741 |
| ADRA1A | -0.7158679 | 0.0091686 |
| ARF5 | -0.7148668 | 0.0173776 |
| RELA | -0.7146891 | 0.0062135 |
| SF3A2 | -0.7124593 | 0.0160163 |
| PCBP1 | -0.7123332 | 0.0041753 |
| SMC2 | -0.7095198 | 0.0028632 |
| SLC7A7 | -0.7094428 | 0.0116941 |
| DUS3L | -0.7092825 | 0.0062575 |
| SPARC | -0.7054388 | 0.0490724 |
| GAA | -0.6988893 | 0.0112459 |
| EPB41L2 | -0.6962555 | 0.0164389 |
| LGI1 | -0.6956888 | 0.0270105 |
| NINJ1 | -0.6941821 | 0.0406189 |
| BHLHE40 | -0.6933279 | 0.0180402 |
| MRPS24 | -0.6926061 | 0.0124295 |
| TMC7 | -0.6908659 | 0.0341801 |
| FAM20C | -0.690734 | 0.0378325 |
| LPL | -0.6906399 | 0.016988 |
| DYNLT1 | -0.6873071 | 0.0041864 |
| PTPN6 | -0.6859374 | 0.0337359 |
| SNX12 | -0.6841596 | 0.0167766 |
| SOX9 | -0.6836043 | 0.0300623 |
| KCNQ3 | -0.6825468 | 0.0359209 |
| DDX27 | -0.682542 | 0.0064323 |
| TMEM176A | -0.6795995 | 0.0357021 |
| ATP11A | -0.6792192 | 0.0092218 |
| VASH1 | -0.6782038 | 0.0068833 |
| CYBA | -0.6774873 | 0.0408565 |
| UBXN6 | -0.6759259 | 0.0259799 |
| DDX59 | -0.6751391 | 0.0243427 |
| CERS5 | -0.673993 | 0.0244189 |
| CNBP | -0.6736741 | 0.0207086 |
| FAM149A | -0.6705889 | 0.0323194 |
| RGS6 | -0.6634671 | 0.0410311 |
| ARHGAP45 | -0.6630622 | 0.0245266 |
| SCIN | -0.6620893 | 0.0245581 |
| TMEM259 | -0.6606043 | 0.0112365 |

| | | |
|---|---|---|
| SPATA13 | -0.6598289 | 0.002482 |
| UCK1 | -0.6596505 | 0.0072971 |
| AKT1 | -0.6579853 | 0.0069166 |
| JOSD2 | -0.6570401 | 0.0079072 |
| FLNC | -0.6567426 | 0.0294744 |
| KDELR1 | -0.6548531 | 0.0035895 |
| CYTH1 | -0.6547628 | 0.0248834 |
| FSCN1 | -0.6505463 | 0.027592 |
| ARPC3 | -0.6481552 | 0.0151691 |
| BAGE3 | -0.6454744 | 0.0195706 |
| MRPL15 | -0.6454397 | 0.0229311 |
| DIRAS1 | -0.6446285 | 0.0285622 |
| GGA2 | -0.6434492 | 0.0253178 |
| CNOT1 | -0.6431755 | 0.0059898 |
| ADAM28 | -0.6414667 | 0.043411 |
| ITGB1 | -0.6400749 | 0.0275443 |
| SH3BGRL3 | -0.639955 | 0.0304534 |
| PFKFB4 | -0.6394122 | 0.0207206 |
| POFUT1 | -0.63847 | 0.0216397 |
| SH3BP1 | -0.6352716 | 0.0308403 |
| RPS13 | -0.6349742 | 0.0496778 |
| ID2 | -0.6306116 | 0.0455058 |
| MZT2A | -0.628365 | 0.0134084 |
| CIAO2B | -0.626914 | 0.0203768 |
| FRG1 | -0.6246003 | 0.0112103 |
| PABPC1 | -0.6231532 | 0.0227611 |
| TATDN2 | -0.6230084 | 0.015596 |
| GSTM2 | -0.6218391 | 0.0149832 |
| PNPLA6 | -0.6218142 | 0.0398495 |
| PABPC4 | -0.6195574 | 0.0147029 |
| SMPD1 | -0.6186692 | 0.0187894 |
| COMT | -0.6180691 | 0.0350857 |
| CNPY3 | -0.6176604 | 0.0160034 |
| LMO1 | -0.6168259 | 0.0209061 |
| RPL22L1 | -0.6166799 | 0.0190444 |
| ADAMTSL2 | -0.6165838 | 0.0276946 |
| FHOD1 | -0.6160983 | 0.0111339 |
| TNRC18 | -0.6148733 | 0.0220397 |
| TRAF3IP2 | -0.6141269 | 0.0203417 |
| SH2B3 | -0.6128323 | 0.0055384 |
| DHRS13 | -0.610254 | 0.0124879 |
| KDM2A | -0.6080778 | 0.0182062 |
| OXTR | -0.6069578 | 0.0384014 |
| ALOX5 | -0.6066037 | 0.0322978 |
| SDHD | -0.6064398 | 0.0068485 |
| AGPAT5 | -0.6053246 | 0.0302564 |
| MEX3D | -0.6043634 | 0.0334752 |
| NRBP1 | -0.6037396 | 0.0350575 |
| TMUB1 | -0.6030657 | 0.0058185 |
| PGLS | -0.6029896 | 0.0178367 |
| UBA6 | -0.6027716 | 0.010273 |
| PYCARD | -0.6027473 | 0.0241373 |
| C3orf70 | -0.6020351 | 0.0463109 |
| HS6ST1 | -0.6008946 | 0.0209535 |
| CORO1C | -0.5988271 | 0.0229868 |
| PIK3AP1 | -0.5982077 | 0.0496261 |

| | | |
|---|---|---|
| PGRMC2 | -0.5981516 | 0.0253173 |
| HSPA9 | -0.5977918 | 0.0285936 |
| RBM23 | -0.5973881 | 0.0200789 |
| LILRA6 | -0.5961853 | 0.0348084 |
| GNA11 | -0.5925527 | 0.0486914 |
| SRCAP | -0.5924474 | 0.0137992 |
| APPBP2 | -0.592246 | 0.0143338 |
| GNA13 | -0.5912076 | 0.0394955 |
| RAD51B | -0.5907536 | 0.0138839 |
| SLC39A1 | -0.589069 | 0.0130637 |
| YTHDF2 | -0.5889383 | 0.0155134 |
| MLXIP | -0.5884901 | 0.0235322 |
| BCKDK | -0.5863936 | 0.0331268 |
| ERGIC1 | -0.5833925 | 0.0082973 |
| MT1HL1 | -0.5823164 | 0.02021 |
| EIF3M | -0.5822026 | 0.0105699 |
| TKT | -0.5818758 | 0.0073745 |
| SELENOH | -0.5813111 | 0.0277573 |
| MICOS13 | -0.5812538 | 0.0374823 |
| JMJD6 | -0.5806641 | 0.017093 |
| PRELID1 | -0.5799764 | 0.0098643 |
| RRP7A | -0.577713 | 0.0261459 |
| UQCRFS1 | -0.5736431 | 0.0430312 |
| PWWP3A | -0.5732649 | 0.0423461 |
| CCDC59 | -0.5727891 | 0.0161767 |
| NT5C2 | -0.5725799 | 0.0456199 |
| COPE | -0.5723047 | 0.0094343 |
| SALL1 | -0.5691721 | 0.0257502 |
| SUPT6H | -0.5675798 | 0.0333628 |
| ZEB1 | -0.5666011 | 0.027794 |
| ERGIC3 | -0.5660227 | 0.0151333 |
| RAPGEF1 | -0.5641761 | 0.0323957 |
| RAB11FIP5 | -0.5640954 | 0.0197553 |
| RNF5 | -0.5633017 | 0.0403549 |
| ECI1 | -0.5631153 | 0.0371121 |
| CLPP | -0.5620124 | 0.0333735 |
| KCNMA1 | -0.5619828 | 0.0266229 |
| PPP1R18 | -0.5614768 | 0.0234846 |
| YKT6 | -0.5598966 | 0.0188095 |
| PCGF2 | -0.5591325 | 0.047266 |
| NIBAN2 | -0.5590734 | 0.0456936 |
| OST4 | -0.5584707 | 0.0442612 |
| WDR5 | -0.5567898 | 0.0254158 |
| TXNL1 | -0.556718 | 0.0325875 |
| PRAMEF6 | -0.556447 | 0.0211422 |
| NR2E1 | -0.55604 | 0.036107 |
| NDUFA13 | -0.5554089 | 0.0197043 |
| VPS28 | -0.5551547 | 0.027626 |
| KBTBD11 | -0.5549834 | 0.0185083 |
| NOTCH3 | -0.5544297 | 0.0365336 |
| EIF3B | -0.5541475 | 0.0153889 |
| PTPRC | -0.5540439 | 0.0444069 |
| PMPCB | -0.5537664 | 0.0157969 |
| PLEKHJ1 | -0.5535978 | 0.0138477 |
| PGPEP1 | -0.552232 | 0.0232643 |
| GPANK1 | -0.5513399 | 0.0431334 |

| | | |
|---|---|---|
| MED15 | -0.5495312 | 0.0100697 |
| HSP90B1 | -0.5488689 | 0.0346876 |
| CAPZB | -0.5469797 | 0.0367908 |
| MPST | -0.5449971 | 0.0347244 |
| COQ10A | -0.544784 | 0.0265043 |
| CD99 | -0.544412 | 0.0486665 |
| DPM2 | -0.5441272 | 0.0122014 |
| RASA3 | -0.543983 | 0.043127 |
| MYO15B | -0.5438817 | 0.019643 |
| VDAC2 | -0.5437199 | 0.0305912 |
| RAB12 | -0.5431633 | 0.0216723 |
| PKN1 | -0.5423985 | 0.009822 |
| RPS10 | -0.541866 | 0.0371977 |
| AP3D1 | -0.5387815 | 0.0299395 |
| ANGPTL2 | -0.53848 | 0.0248502 |
| CCAR2 | -0.5381711 | 0.0239666 |
| COL5A1 | -0.5372401 | 0.0304851 |
| MRI1 | -0.5352054 | 0.0435055 |
| QRICH1 | -0.5334143 | 0.0307346 |
| RASSF4 | -0.5317454 | 0.0494664 |
| COMMD5 | -0.530727 | 0.0084644 |
| DDX1 | -0.5293856 | 0.018225 |
| C1QTNF5 | -0.5252597 | 0.0480394 |
| LMO2 | -0.5248484 | 0.0471972 |
| TTLL12 | -0.5236781 | 0.0344775 |
| MRPS17 | -0.5234336 | 0.0379249 |
| TSEN34 | -0.5203822 | 0.0309339 |
| PLD2 | -0.5202982 | 0.0298388 |
| MXRA7 | -0.5199934 | 0.0380353 |
| PRRC2B | -0.5180846 | 0.0467571 |
| MPLKIP | -0.5179586 | 0.0424655 |
| MT1A | -0.5175593 | 0.0391452 |
| FKBP8 | -0.5164604 | 0.0400683 |
| TOR1AIP1 | -0.5156887 | 0.0275471 |
| R3HCC1 | -0.5144297 | 0.0416739 |
| ARHGEF6 | -0.5129356 | 0.0372524 |
| MFSD2A | -0.5128865 | 0.0265341 |
| LIN7A | -0.5098093 | 0.0431016 |
| CFAP251 | -0.5088348 | 0.0423208 |
| PPIL2 | -0.5067299 | 0.0325642 |
| NMT1 | -0.5062802 | 0.0414871 |
| ABCB8 | -0.5062182 | 0.03797 |
| PLEKHA2 | -0.5056758 | 0.0491256 |
| PIEZO1 | -0.5047728 | 0.0237377 |
| VHL | -0.5038639 | 0.0461879 |
| TICAM1 | -0.5009166 | 0.0298681 |
| FAM216A | -0.5006589 | 0.0326171 |
| CREB1 | -0.5001581 | 0.022587 |
| MAP2K2 | -0.4997153 | 0.0204998 |
| ATP6V1D | -0.4979242 | 0.0391052 |
| TP53BP2 | -0.4963727 | 0.0400549 |
| HSD17B4 | -0.4961522 | 0.0420473 |
| STX8 | -0.4951154 | 0.0459742 |
| ARID1B | -0.4950419 | 0.04453 |
| DDAH2 | -0.4932424 | 0.0368506 |
| CKLF | -0.4932194 | 0.0262455 |

| | | |
|---|---|---|
| SRA1 | -0.4930314 | 0.0305237 |
| BCAP31 | -0.4925872 | 0.0329744 |
| PLXNB2 | -0.4908359 | 0.0314444 |
| SLC1A5 | -0.4907138 | 0.044612 |
| LRFN4 | -0.4906897 | 0.048946 |
| DCK | -0.4825747 | 0.0348932 |
| CSNK1E | -0.4820252 | 0.0441899 |
| TAF4 | -0.4815249 | 0.0437699 |
| NDUFS2 | -0.477311 | 0.0370397 |
| HUWE1 | -0.4772063 | 0.0430378 |
| DEGS1 | -0.4758563 | 0.0340192 |
| C1orf43 | -0.4744245 | 0.0286061 |
| TIMM8A | -0.4729313 | 0.0386506 |
| VGLL4 | -0.4727837 | 0.0379867 |
| WDR45B | -0.4709373 | 0.0360815 |
| SFI1 | -0.4702554 | 0.0354444 |
| LRWD1 | -0.4659777 | 0.0489073 |
| SMG1 | -0.4656298 | 0.0461773 |
| PCDH10 | -0.4640796 | 0.0331202 |
| PPP1R9B | -0.461386 | 0.0260936 |
| FOXO3 | -0.4606748 | 0.0346056 |
| SNAPIN | -0.4591833 | 0.0316594 |
| SLC41A3 | -0.4585768 | 0.047872 |
| TFR2 | -0.4568213 | 0.0478203 |
| VPS35 | -0.4564066 | 0.0249951 |
| ALDH4A1 | -0.4562506 | 0.03544 |
| TNFRSF1B | -0.4548164 | 0.0402812 |
| COPG1 | -0.4539764 | 0.0374648 |
| FLII | -0.4529814 | 0.0271644 |
| LSM14A | -0.4472742 | 0.0453442 |
| SLC25A1 | -0.446908 | 0.0346104 |
| PKD2 | -0.4449837 | 0.0462183 |
| PHACTR4 | -0.4324408 | 0.0446425 |
| DDX46 | -0.430363 | 0.0465611 |
| DBNL | -0.4298013 | 0.032535 |
| SRC | -0.4265275 | 0.0403582 |
| RAB13 | -0.4175504 | 0.0420797 |
| TSFM | -0.4153521 | 0.0455168 |
| SLC33A1 | -0.4089932 | 0.036816 |

| ID | logFC | P.Value |
|---|---|---|
| IGF2 | 1.6789914 | 0.0007921 |
| NRGN | 1.6675348 | 0.0053165 |
| PITX1 | 1.6508146 | 0.0005549 |
| RNASE1 | 1.6402113 | 0.0018627 |
| IGKC | 1.6341025 | 0.001617 |
| CHN1 | 1.6043383 | 0.000239 |
| F13A1 | 1.4848667 | 0.0001851 |
| FOS | 1.428404 | 0.0299027 |
| IGHG4 | 1.3989946 | 0.001388 |
| EGR1 | 1.3779621 | 0.0208264 |
| NDUFS1 | 1.2858085 | 0.0000214 |
| SNAP25 | 1.2424923 | 0.0224096 |
| ALMS1 | 1.2133173 | 0.0025099 |
| C4B | 1.1991814 | 0.0061898 |
| OGN | 1.1091701 | 0.0095475 |
| CRTAM | 1.1059582 | 0.0002241 |
| LEF1 | 1.0937478 | 0.0011333 |
| IGHG2 | 1.0916691 | 0.0157975 |
| GRIN1 | 1.074898 | 0.0214933 |
| APLP1 | 1.0674123 | 0.0407923 |
| CALM3 | 1.057922 | 0.0094413 |
| SLC1A2 | 1.0552945 | 0.0223779 |
| LDOC1 | 1.0530725 | 0.0005608 |
| FGF12 | 1.0405419 | 0.0071962 |
| KCNJ11 | 1.0204205 | 0.0018892 |
| CLDN11 | 1.0175789 | 0.0477177 |
| VAMP2 | 0.9964397 | 0.0117559 |
| CASD1 | 0.984824 | 0.0042135 |
| CTIF | 0.9847798 | 0.0014306 |
| HOXB3 | 0.9821507 | 0.0020959 |
| PLPP3 | 0.9765597 | 0.0089307 |
| ADCY2 | 0.9740211 | 0.0003348 |
| CXCL2 | 0.9707422 | 0.0031994 |
| DGKA | 0.9629874 | 0.003403 |
| CAMK2N1 | 0.959096 | 0.0169297 |
| RIMS3 | 0.957883 | 0.0045955 |
| AMPH | 0.9570559 | 0.0036793 |
| DNM1 | 0.9526351 | 0.019853 |
| ZFAND5 | 0.9500484 | 0.0044711 |
| PRKACB | 0.9476663 | 0.0026519 |
| MBNL1 | 0.9422687 | 0.0040202 |
| KLHL7 | 0.9418055 | 0.0062917 |
| TMEM130 | 0.9324504 | 0.0092811 |
| EMP1 | 0.9321706 | 0.0214806 |
| C1QTNF4 | 0.927861 | 0.001377 |
| CFH | 0.9260553 | 0.0243156 |
| C1GALT1 | 0.9247792 | 0.0000411 |
| NFIB | 0.9247469 | 0.020066 |
| RAD52 | 0.9136237 | 0.0024457 |
| SMARCA1 | 0.9129341 | 0.0042131 |
| IGHG1 | 0.91261 | 0.0382914 |
| TAAR2 | 0.912304 | 0.0114582 |
| KCNJ10 | 0.9077024 | 0.0032247 |
| TSPYL2 | 0.9060223 | 0.0047731 |
| CABP1 | 0.905972 | 0.0021837 |

| | | |
|---|---|---|
| KRCC1 | 0.9005364 | 0.0004618 |
| ADAMDEC1 | 0.9003299 | 0.0131295 |
| COL20A1 | 0.898632 | 0.0195161 |
| KIF5C | 0.897812 | 0.0300647 |
| CNTN2 | 0.8965247 | 0.0354192 |
| SCG5 | 0.896506 | 0.0038859 |
| CPLX2 | 0.885416 | 0.0067582 |
| UBLCP1 | 0.8843508 | 0.0041904 |
| HDAC9 | 0.8843065 | 0.0079935 |
| CDH18 | 0.8789822 | 0.0060799 |
| GATA4 | 0.878884 | 0.0012153 |
| PTN | 0.8766359 | 0.0184566 |
| SNX22 | 0.8670765 | 0.0166309 |
| SCN8A | 0.8634724 | 0.0053564 |
| NEFM | 0.8590418 | 0.0333329 |
| SLPI | 0.8573302 | 0.0068103 |
| HHATL | 0.8541268 | 0.0419572 |
| CEMIP | 0.8525766 | 0.0347574 |
| ZDHHC2 | 0.8524516 | 0.0124583 |
| CALM2 | 0.8517681 | 0.0490433 |
| DYNC2LI1 | 0.8461304 | 0.0008166 |
| ZNF346 | 0.8414914 | 0.0293731 |
| MAPRE2 | 0.8406887 | 0.0044527 |
| APBB2 | 0.84049 | 0.0021553 |
| SMCHD1 | 0.8380332 | 0.0011523 |
| NFYA | 0.8358163 | 0.03332 |
| LAMTOR3 | 0.8314004 | 0.0231934 |
| FXYD1 | 0.8289237 | 0.0310243 |
| MPEG1 | 0.8272061 | 0.0073881 |
| FBXL17 | 0.8271041 | 0.0131189 |
| DEFB134 | 0.8267108 | 0.0015758 |
| COL11A1 | 0.8240089 | 0.0040987 |
| PLIN4 | 0.8231636 | 0.0045898 |
| AK5 | 0.8222695 | 0.0029429 |
| KRT15 | 0.8218208 | 0.0019892 |
| FOLR2 | 0.8202229 | 0.0332557 |
| EEF1AKMT2 | 0.8200678 | 0.0297408 |
| PLEKHH1 | 0.8136892 | 0.0289323 |
| CEP170B | 0.8090064 | 0.0151291 |
| IGSF11 | 0.804349 | 0.0037204 |
| GABBR1 | 0.8038011 | 0.0122976 |
| TSPYL1 | 0.7989715 | 0.0096839 |
| CPSF3 | 0.797574 | 0.0019741 |
| GPATCH11 | 0.7965449 | 0.0008472 |
| OSGIN1 | 0.7962563 | 0.0119252 |
| C3orf14 | 0.7945224 | 0.0006787 |
| SRSF5 | 0.7906234 | 0.016373 |
| FOLR1 | 0.7905604 | 0.0053501 |
| SV2B | 0.7903546 | 0.0314751 |
| SLC22A17 | 0.7879316 | 0.0231042 |
| ZNF709 | 0.7875861 | 0.0147741 |
| PLD3 | 0.7871156 | 0.0154723 |
| SELENOP | 0.7844032 | 0.0435525 |
| ANK1 | 0.7833559 | 0.0201961 |
| C16orf78 | 0.7801624 | 0.0023475 |
| DLL1 | 0.779604 | 0.0065305 |

| | | |
|---|---|---|
| XAF1 | 0.7785262 | 0.0287849 |
| ATL1 | 0.7785117 | 0.0045814 |
| CCL3L3 | 0.7775129 | 0.0398172 |
| FAM156A | 0.7707358 | 0.0110891 |
| SRCIN1 | 0.7666561 | 0.0252614 |
| NSG2 | 0.7641435 | 0.0179961 |
| CEP83 | 0.7621162 | 0.010188 |
| COL1A1 | 0.7612845 | 0.0260096 |
| FARP2 | 0.7612221 | 0.0086103 |
| CAMK4 | 0.759322 | 0.0078882 |
| NDST1 | 0.7542781 | 0.0249982 |
| C1QL3 | 0.754158 | 0.0127885 |
| CLSTN3 | 0.7526002 | 0.0273661 |
| IFI44 | 0.7511358 | 0.0222008 |
| SLC12A4 | 0.7443497 | 0.007438 |
| DNM3 | 0.7425965 | 0.0109174 |
| LGMN | 0.7405448 | 0.0406457 |
| DGKB | 0.7398003 | 0.0263025 |
| ACTR2 | 0.7359068 | 0.0084664 |
| SGCD | 0.7319719 | 0.0040064 |
| LANCL1 | 0.7289468 | 0.0253698 |
| HACL1 | 0.7289025 | 0.0138235 |
| MASTL | 0.7275473 | 0.0071092 |
| PIP4K2C | 0.7240566 | 0.0087134 |
| CALCRL | 0.7238946 | 0.0465014 |
| MAZ | 0.7226523 | 0.0254388 |
| SPACA3 | 0.7224624 | 0.0187839 |
| SRD5A1 | 0.7219341 | 0.0349542 |
| RGS4 | 0.7191858 | 0.0385441 |
| TYR | 0.7191406 | 0.0194682 |
| C1orf216 | 0.7161206 | 0.0114516 |
| PDCD10 | 0.7156721 | 0.0086849 |
| FAM13B | 0.7142433 | 0.0120632 |
| DUSP5 | 0.7130481 | 0.0178468 |
| NDUFAF2 | 0.7128407 | 0.0032635 |
| RBM12B | 0.7111366 | 0.0195521 |
| NEURL1B | 0.7111104 | 0.0395546 |
| SPATA24 | 0.7083748 | 0.0031555 |
| OR51J1 | 0.7072305 | 0.0049164 |
| APOBEC3A_ | 0.7046461 | 0.0296751 |
| STAT1 | 0.7038669 | 0.0448729 |
| SPRY3 | 0.7030491 | 0.0219369 |
| PARP15 | 0.7026755 | 0.006867 |
| RGN | 0.7022551 | 0.0047062 |
| ZNF506 | 0.7017092 | 0.028399 |
| ESR1 | 0.7009082 | 0.0038849 |
| OARD1 | 0.7006537 | 0.034839 |
| IGLL5 | 0.698082 | 0.0341509 |
| LRCOL1 | 0.6966001 | 0.0396622 |
| FOLH1 | 0.6951756 | 0.0243585 |
| KLHL9 | 0.6918862 | 0.0077942 |
| SAAL1 | 0.6884983 | 0.0166907 |
| IGHG3 | 0.6828478 | 0.0462508 |
| GXYLT2 | 0.6803037 | 0.0048794 |
| TCEAL5 | 0.6772599 | 0.0182363 |
| TTC31 | 0.6770324 | 0.0089235 |

| | | |
|---|---|---|
| RASL10B | 0.6766279 | 0.0308826 |
| PAN3 | 0.6753933 | 0.0114323 |
| CDX1 | 0.6751941 | 0.0127571 |
| PRC1 | 0.6734319 | 0.0056045 |
| ZNF827 | 0.6732861 | 0.0048685 |
| AREG | 0.6712446 | 0.0116682 |
| PCMTD2 | 0.6700216 | 0.0108405 |
| TBCEL | 0.6690647 | 0.0073217 |
| RUSC2 | 0.6689486 | 0.0012259 |
| NMNAT2 | 0.6679088 | 0.0291595 |
| GABBR2 | 0.6678173 | 0.0251371 |
| SRP54 | 0.667244 | 0.0400247 |
| LPGAT1 | 0.6670278 | 0.0055976 |
| ZNF805 | 0.665695 | 0.0050166 |
| NECTIN3 | 0.6649479 | 0.0157741 |
| CXCR1 | 0.6631851 | 0.0248605 |
| COLEC12 | 0.6603426 | 0.0342346 |
| CLEC11A | 0.6601537 | 0.0197141 |
| SMARCA4 | 0.6595065 | 0.0063593 |
| ZNF821 | 0.6592155 | 0.0047694 |
| ARCN1 | 0.6589906 | 0.0260899 |
| DDHD2 | 0.6583581 | 0.0027846 |
| ARHGAP23 | 0.6575306 | 0.0115875 |
| EPDR1 | 0.6560564 | 0.0170703 |
| PIK3R1 | 0.6560158 | 0.0331231 |
| SNAPC4 | 0.6551397 | 0.0287378 |
| TMEM151A | 0.6545048 | 0.0273225 |
| ABCB9 | 0.6542423 | 0.0308685 |
| MICAL3 | 0.6538649 | 0.0434737 |
| PDCD4 | 0.6535028 | 0.0151452 |
| HELLS | 0.652539 | 0.009677 |
| PAFAH1B1 | 0.6495761 | 0.026547 |
| ARHGAP25 | 0.6465573 | 0.0090087 |
| TMEM101 | 0.6462623 | 0.0116974 |
| BLCAP | 0.6457045 | 0.0182616 |
| CCDC179 | 0.6455985 | 0.0135628 |
| PLXNA1 | 0.6450654 | 0.0172442 |
| SELP | 0.6442628 | 0.0482492 |
| OSBPL1A | 0.642864 | 0.0288505 |
| RABGAP1L | 0.6411981 | 0.004598 |
| CYB5D2 | 0.6410489 | 0.0344033 |
| WBP1L | 0.6393116 | 0.010527 |
| CSTF1 | 0.6379738 | 0.0143689 |
| DCTD | 0.6379149 | 0.0114518 |
| PPP1R3E | 0.6376675 | 0.0142669 |
| METTL14 | 0.6359565 | 0.0083658 |
| APOL1 | 0.6357053 | 0.0106592 |
| BMP8A | 0.6348789 | 0.0301046 |
| NAA38 | 0.6338176 | 0.0385007 |
| MORN3 | 0.6336306 | 0.0141354 |
| USP14 | 0.6327833 | 0.0323056 |
| EXOSC10 | 0.6325062 | 0.0230705 |
| ATP5MK | 0.6323976 | 0.0192943 |
| ZNF69 | 0.6322174 | 0.019553 |
| PRNP | 0.6315985 | 0.0408296 |
| HAPLN2 | 0.6308591 | 0.0103029 |

| | | |
|---|---|---|
| MOK | 0.6293779 | 0.0056642 |
| DENND1B | 0.6289568 | 0.039613 |
| GIMAP8 | 0.6288835 | 0.0242506 |
| METTL3 | 0.6226788 | 0.0231244 |
| SCNM1 | 0.6224464 | 0.0055098 |
| SLC35B4 | 0.6208483 | 0.0493497 |
| MDC1 | 0.617982 | 0.0143161 |
| PTS | 0.6169652 | 0.0106969 |
| NUCB2 | 0.6165619 | 0.0191609 |
| SLC12A5 | 0.6155926 | 0.0395364 |
| DGCR8 | 0.6152041 | 0.0321689 |
| KLHDC1 | 0.6140199 | 0.0194506 |
| GTF2H1 | 0.6138457 | 0.0167613 |
| MPP3 | 0.6135216 | 0.005964 |
| DSTN | 0.6100893 | 0.0445869 |
| RDH13 | 0.6089261 | 0.0138849 |
| NPDC1 | 0.6078751 | 0.025457 |
| HIRA | 0.607685 | 0.0108545 |
| ECD | 0.6054304 | 0.0249023 |
| C1D | 0.6042704 | 0.0325798 |
| CDH1 | 0.6042401 | 0.0446526 |
| NPNT | 0.6040561 | 0.0453311 |
| GLUD1 | 0.603954 | 0.0384698 |
| MSTO1 | 0.602919 | 0.0332758 |
| CKS2 | 0.6026856 | 0.0096502 |
| C2orf69 | 0.6024963 | 0.010698 |
| ENPP4 | 0.601858 | 0.0465065 |
| UBE2Z | 0.6017372 | 0.0173932 |
| JMJD1C | 0.601617 | 0.0158931 |
| PLPP4 | 0.6006361 | 0.0233601 |
| ZNF235 | 0.6005679 | 0.0113058 |
| FAM72B | 0.599448 | 0.0180021 |
| ZNF362 | 0.5972423 | 0.0085518 |
| EPM2AIP1 | 0.5965805 | 0.0117935 |
| PAPLN | 0.5962386 | 0.034997 |
| SLC24A3 | 0.5952402 | 0.0112192 |
| SRRM3 | 0.5951562 | 0.0154019 |
| NEK4 | 0.5940416 | 0.0446695 |
| HARS1 | 0.5929703 | 0.0251946 |
| NDUFA9 | 0.5904621 | 0.0279823 |
| TTBK2 | 0.5888763 | 0.0063953 |
| GRIK5 | 0.5876591 | 0.008922 |
| TTC3 | 0.5867506 | 0.0134798 |
| CRY1 | 0.5865795 | 0.0142382 |
| SLC25A17 | 0.5863469 | 0.0308687 |
| KLC3 | 0.5860696 | 0.025271 |
| VDR | 0.5858892 | 0.0263046 |
| NR2C1 | 0.5854791 | 0.0266962 |
| CCDC14 | 0.5850119 | 0.0140589 |
| OR8D1 | 0.5825703 | 0.0146243 |
| TCEA3 | 0.5823716 | 0.0149624 |
| RNF141 | 0.5815332 | 0.0131743 |
| ESCO2 | 0.5809666 | 0.0229959 |
| SATB2 | 0.5803253 | 0.007181 |
| RGS7 | 0.5789342 | 0.0336722 |
| POLR3F | 0.5784837 | 0.0360217 |

| | | |
|---|---|---|
| CHSY3 | 0.5779103 | 0.0418534 |
| ERBB4 | 0.5778755 | 0.0262117 |
| RAG1 | 0.5777701 | 0.0331192 |
| DIPK1C | 0.5777094 | 0.0445491 |
| BRME1 | 0.577028 | 0.0139121 |
| CABLES1 | 0.5767004 | 0.0499443 |
| TSGA10 | 0.5737447 | 0.0218799 |
| ATAD1 | 0.5726647 | 0.0155173 |
| DMD | 0.57077 | 0.0415463 |
| NBPF26 | 0.5704699 | 0.0197364 |
| FGFR1OP2 | 0.5694438 | 0.0206013 |
| CSTF3 | 0.569398 | 0.0242442 |
| ESRP1 | 0.5691603 | 0.0219486 |
| CEP126 | 0.5684261 | 0.0329092 |
| CCR10 | 0.568208 | 0.0418995 |
| PPP2R1B | 0.5671871 | 0.0400592 |
| CHGB | 0.5668643 | 0.0251489 |
| MON2 | 0.565859 | 0.0105772 |
| S100A12 | 0.5642401 | 0.0181064 |
| CCNC | 0.5640932 | 0.0134582 |
| ZNF195 | 0.5593947 | 0.0196408 |
| CLRN1 | 0.5589343 | 0.0449559 |
| CLDN34 | 0.5568787 | 0.0209201 |
| NANOS3 | 0.5558354 | 0.0233985 |
| ZNF436 | 0.555577 | 0.0452966 |
| NBR1 | 0.5540597 | 0.0464413 |
| THBS3 | 0.5535939 | 0.0231423 |
| USP54 | 0.5525732 | 0.0457349 |
| SCYL1 | 0.5520573 | 0.0285484 |
| TOM1 | 0.5520069 | 0.0485902 |
| TRAPPC10 | 0.5519939 | 0.0312909 |
| NPIPB6 | 0.5511153 | 0.0260472 |
| KYNU | 0.5495925 | 0.0165611 |
| UAP1 | 0.5493775 | 0.02598 |
| ESF1 | 0.5493327 | 0.0145688 |
| RECQL5 | 0.5491944 | 0.0198075 |
| SHISA4 | 0.549177 | 0.0409881 |
| BBIP1 | 0.5489483 | 0.0489378 |
| AGPAT2 | 0.5488983 | 0.0337487 |
| PHACTR1 | 0.5485544 | 0.0229224 |
| CHIC1 | 0.5474613 | 0.0386272 |
| FABP1 | 0.5470916 | 0.0257077 |
| MIA | 0.5458529 | 0.0395043 |
| SAA2 | 0.5457429 | 0.0441338 |
| EIF4G3 | 0.5455725 | 0.0278566 |
| USP25 | 0.5455534 | 0.0159349 |
| WWOX | 0.5455085 | 0.0397192 |
| CTTN | 0.5453776 | 0.0496068 |
| LRRC75B | 0.5433379 | 0.0288525 |
| WHRN | 0.5414868 | 0.0353681 |
| BCKDHB | 0.5407443 | 0.0480304 |
| SLC25A27 | 0.5403897 | 0.0127187 |
| CAMTA1 | 0.5393942 | 0.0430572 |
| RNF170 | 0.5393516 | 0.0067578 |
| POLR2I | 0.5392855 | 0.0432883 |
| PSMB2 | 0.5384 | 0.0107494 |

| | | |
|---|---|---|
| GPR158 | 0.5379842 | 0.0418385 |
| MPPED2 | 0.5369431 | 0.0320478 |
| PHIP | 0.5367383 | 0.0175447 |
| NRG1 | 0.5351946 | 0.0312109 |
| TUBG2 | 0.5336895 | 0.0180387 |
| MRM1 | 0.5332699 | 0.0347345 |
| ETFRF1 | 0.5321137 | 0.0379224 |
| SLTM | 0.531875 | 0.0169525 |
| PACRGL | 0.5318272 | 0.0164526 |
| PHLDA3 | 0.5304845 | 0.0219582 |
| XKR8 | 0.5304179 | 0.0148444 |
| LIME1 | 0.5302179 | 0.0296353 |
| NEO1 | 0.5293843 | 0.015985 |
| RRP36 | 0.5284824 | 0.0280224 |
| TSPAN2 | 0.5283536 | 0.0315582 |
| NDUFAF8 | 0.5276218 | 0.0222183 |
| ARGLU1 | 0.5267813 | 0.0434972 |
| SENP7 | 0.5265801 | 0.0267968 |
| MYO9A | 0.5253973 | 0.0431833 |
| SMPDL3A | 0.5251693 | 0.0411213 |
| DEDD | 0.5248783 | 0.0126043 |
| PDLIM1 | 0.5219255 | 0.0296818 |
| QSOX1 | 0.5199793 | 0.0386028 |
| NFAM1 | 0.5192775 | 0.0279533 |
| COG2 | 0.5191083 | 0.0265223 |
| SRARP | 0.5184065 | 0.0443012 |
| IFIT2 | 0.5181051 | 0.0194852 |
| KDM4A | 0.5178146 | 0.0400326 |
| ZC3H14 | 0.517688 | 0.0449897 |
| MMP3 | 0.5175907 | 0.0297857 |
| SMIM19 | 0.5164801 | 0.0261056 |
| LRPPRC | 0.5159975 | 0.0454325 |
| ELK1 | 0.5153676 | 0.022324 |
| LRP6 | 0.5149624 | 0.0275868 |
| KANSL2 | 0.5145771 | 0.0183963 |
| PDGFA | 0.5141166 | 0.0470019 |
| KCNT1 | 0.5134726 | 0.039762 |
| RANBP10 | 0.5127465 | 0.0473954 |
| ATMIN | 0.5096624 | 0.0418448 |
| SH3YL1 | 0.5095349 | 0.0496974 |
| CSRNP2 | 0.5081889 | 0.0412809 |
| PDE7A | 0.5065974 | 0.0268199 |
| GART | 0.50624 | 0.0194537 |
| RIOX2 | 0.5061627 | 0.0238242 |
| B3GALNT1 | 0.5037282 | 0.0236365 |
| ZNF609 | 0.5035425 | 0.01873 |
| KANK1 | 0.5028153 | 0.0419575 |
| PPP1R37 | 0.5021211 | 0.0261851 |
| STARD3NL | 0.5018101 | 0.0401488 |
| KIF3B | 0.5011625 | 0.0262132 |
| PTPN13 | 0.5010229 | 0.0217694 |
| USP34 | 0.5008673 | 0.0343071 |
| UFSP2 | 0.5001067 | 0.0256278 |
| MYBPC2 | 0.4990442 | 0.0236708 |
| DDX58 | 0.4990165 | 0.0396104 |
| NOS2 | 0.4988515 | 0.0288666 |

| | | |
|---|---|---|
| NAP1L4 | 0.4957702 | 0.0405183 |
| TEX264 | 0.4947683 | 0.0407673 |
| JAGN1 | 0.492057 | 0.0390586 |
| C2orf88 | 0.4918108 | 0.0476337 |
| STOM | 0.4887981 | 0.0310465 |
| SELENOO | 0.4887861 | 0.0191466 |
| POLRMT | 0.4883673 | 0.0276581 |
| TJP1 | 0.4870665 | 0.0217438 |
| FOXO1 | 0.4861369 | 0.044244 |
| METTL21A | 0.4851833 | 0.0277048 |
| PDLIM7 | 0.4831161 | 0.0397329 |
| MRPS34 | 0.4824788 | 0.0392269 |
| LIMCH1 | 0.4820646 | 0.0405686 |
| COPS7A | 0.4816415 | 0.0230987 |
| NDEL1 | 0.4803945 | 0.0287115 |
| RAB39A | 0.4803373 | 0.0435222 |
| MPC1L | 0.4800561 | 0.0394467 |
| FARSA | 0.4787655 | 0.0345545 |
| FBH1 | 0.4781444 | 0.0290422 |
| TIMM50 | 0.4776007 | 0.0470677 |
| KIAA0754 | 0.4765369 | 0.0319528 |
| PCSK7 | 0.474995 | 0.0346171 |
| ATXN7L3 | 0.4731441 | 0.046624 |
| CDK14 | 0.4711793 | 0.0489594 |
| ZNF45 | 0.4707327 | 0.0348196 |
| ZNF532 | 0.4701226 | 0.0381008 |
| EED | 0.469508 | 0.024058 |
| SMIM8 | 0.4679117 | 0.0257542 |
| RFK | 0.4669962 | 0.0339225 |
| EEF1E1 | 0.4665835 | 0.0372327 |
| SS18L1 | 0.4652047 | 0.0426347 |
| SIGLEC15 | 0.4648724 | 0.0464188 |
| ABCB7 | 0.4644042 | 0.0387288 |
| POLN | 0.4643446 | 0.025375 |
| METTL5 | 0.4642611 | 0.0456569 |
| HRNR | 0.4617419 | 0.0395977 |
| SERTAD1 | 0.4613716 | 0.0364532 |
| DHRS11 | 0.4596569 | 0.0463639 |
| RASA1 | 0.4556433 | 0.037251 |
| SNRNP40 | 0.4553561 | 0.0340475 |
| ZNF283 | 0.4544591 | 0.0359762 |
| C6orf132 | 0.4538491 | 0.0467087 |
| CBWD2 | 0.4504565 | 0.0418192 |
| DHX30 | 0.44802 | 0.0494061 |
| ZZEF1 | 0.4471495 | 0.046413 |
| CCSAP | 0.4452755 | 0.0467878 |
| FOXN2 | 0.4407219 | 0.0465693 |
| PRRG3 | 0.439597 | 0.0335164 |
| EFCAB13 | 0.4395259 | 0.0413388 |
| C5orf24 | 0.4379853 | 0.0421981 |
| C1QTNF2 | 0.4379087 | 0.0401499 |
| BTN3A3 | 0.432431 | 0.0412709 |
| RXYLT1 | 0.4308214 | 0.0400262 |
| ADAR | 0.4235004 | 0.0409324 |
| C5orf15 | 0.4139224 | 0.0416395 |
| MTA3 | 0.4102437 | 0.046842 |

| | | |
|---|---|---|
| KDM5C | 0.4075365 | 0.0468691 |
| NR0B1 | 0.4064771 | 0.0329357 |
| METTL6 | 0.3911415 | 0.0440806 |
| SCLT1 | 0.3838622 | 0.0371658 |

Table S7

| ID | logFC | P.Value |
|---|---|---|
| SFRP4 | -2.8128826 | 0.0000158 |
| RNASE1 | -2.657715 | 0.0001549 |
| IGKC | -2.5321957 | 0.0000016 |
| IGHG2 | -2.4736153 | 0.0000005 |
| SFRP2 | -2.4441727 | 0.0002998 |
| HSPA1A | -2.273249 | 0.0132022 |
| C1S | -2.2390515 | 0.0019551 |
| IGHG4 | -2.2163944 | 0.0000035 |
| FKBP5 | -2.1824098 | 0.0002576 |
| CCN2 | -2.12248 | 0.0059394 |
| HBA1 | -2.0982289 | 0.0016125 |
| SELENOP | -1.9703198 | 0.0176369 |
| PLTP | -1.9446039 | 0.0049946 |
| C1QA | -1.8936972 | 0.0056084 |
| COL3A1 | -1.8845997 | 0.0046664 |
| IGFBP5 | -1.8745168 | 0.0173186 |
| C1QC | -1.8286292 | 0.0070731 |
| FRZB | -1.8277615 | 0.0100564 |
| IGHG3 | -1.769627 | 0.0000659 |
| HSPA6 | -1.7409873 | 0.0043742 |
| MGP | -1.7341016 | 0.0038812 |
| CD163 | -1.7029873 | 0.0126839 |
| LUM | -1.6898142 | 0.0023563 |
| MRC1 | -1.6572454 | 0.0001812 |
| KLF2 | -1.6430103 | 0.0010107 |
| IGFBP4 | -1.6415658 | 0.0070957 |
| CEBPD | -1.6382497 | 0.0100775 |
| IGHG1 | -1.6348182 | 0.0005267 |
| SERPING1 | -1.6278292 | 0.0110565 |
| FBLN2 | -1.6075885 | 0.0004476 |
| HLA-DPB1 | -1.6074454 | 0.0077714 |
| C7 | -1.6009759 | 0.017041 |
| OGN | -1.5893602 | 0.0024844 |
| CLEC3B | -1.5874203 | 0.000177 |
| DNAJB1 | -1.5838622 | 0.0345506 |
| NPC2 | -1.5796185 | 0.0324872 |
| MSR1 | -1.5490336 | 0.0269704 |
| TSC22D3 | -1.5400149 | 0.001541 |
| OLFML3 | -1.5269219 | 0.0088686 |
| C1R | -1.5043749 | 0.0341234 |
| TMEM176B | -1.4828967 | 0.0052941 |
| ZFP36L2 | -1.4707743 | 0.0041026 |
| C1QB | -1.4677922 | 0.0317828 |
| CD68 | -1.4601109 | 0.013923 |
| CFD | -1.4588702 | 0.0014183 |
| NUPR1 | -1.4553449 | 0.0258922 |

| | | |
|---|---|---|
| CFH | -1.4525667 | 0.0024532 |
| HCLS1 | -1.4522932 | 0.0015611 |
| DCN | -1.448362 | 0.0439163 |
| DAB2 | -1.4315855 | 0.000201 |
| TXNIP | -1.4313836 | 0.0172399 |
| NNMT | -1.4286814 | 0.0133817 |
| F13A1 | -1.4248098 | 0.0013776 |
| IFITM1 | -1.4108588 | 0.0042257 |
| MMP2 | -1.3994265 | 0.0005243 |
| IER5 | -1.3885488 | 0.004776 |
| HERPUD1 | -1.3859778 | 0.0002355 |
| SYPL2 | -1.3495781 | 0.0411502 |
| HEXA | -1.337436 | 0.000374 |
| WFS1 | -1.3272869 | 0.0053184 |
| CTSS | -1.3265409 | 0.009809 |
| SLA | -1.3199226 | 0.002243 |
| TIMP3 | -1.3160431 | 0.0198991 |
| HBB | -1.3054609 | 0.0266849 |
| CEMIP | -1.2896522 | 0.0043469 |
| MS4A7 | -1.2858998 | 0.0148494 |
| PPP1R15B | -1.2791279 | 0.0049485 |
| STAT6 | -1.2750854 | 0.0003649 |
| ZBTB16 | -1.2745055 | 0.0087776 |
| HSPE1 | -1.268248 | 0.0115933 |
| TBC1D14 | -1.2573958 | 0.0018528 |
| ZIC4 | -1.2560781 | 0.0131209 |
| GSTM1 | -1.2557435 | 0.013944 |
| PFKFB2 | -1.2350492 | 0.0046349 |
| PHB2 | -1.2308516 | 0.0036695 |
| DCTN4 | -1.2159286 | 0.0194855 |
| FOXP1 | -1.2145204 | 0.0316608 |
| PRRX1 | -1.2119783 | 0.0185718 |
| FGL2 | -1.1836373 | 0.0023299 |
| SMAD5 | -1.180252 | 0.0008219 |
| FBLN1 | -1.1789882 | 0.0120698 |
| NUDCD2 | -1.1783176 | 0.0091958 |
| THBS1 | -1.1678551 | 0.0156019 |
| HCFC2 | -1.1671671 | 0.0054307 |
| CYB561A3 | -1.1578872 | 0.0144303 |
| ZIC1 | -1.1534349 | 0.0018277 |
| GAS6 | -1.146417 | 0.0012682 |
| OXTR | -1.1422939 | 0.0212685 |
| MXRA8 | -1.1420588 | 0.0416748 |
| CMKLR1 | -1.1396877 | 0.0038835 |
| SMAP2 | -1.1391182 | 0.005845 |
| BSND | -1.1379805 | 0.0064332 |
| ALDH1A1 | -1.1377661 | 0.0027201 |
| SLCO2B1 | -1.1287297 | 0.0393262 |

| | | |
|---|---|---|
| PLEKHO1 | -1.1275175 | 0.0044227 |
| IFI44L | -1.12601 | 0.0345979 |
| LGMN | -1.1219275 | 0.0370654 |
| BDH2 | -1.1189935 | 0.0200779 |
| AHNAK | -1.1156795 | 0.0162938 |
| SDCBP | -1.112324 | 0.0182388 |
| OSBPL2 | -1.1077347 | 0.0028134 |
| KLF9 | -1.1060945 | 0.0031987 |
| DEPP1 | -1.105283 | 0.0254636 |
| CD37 | -1.1041823 | 0.0204202 |
| VPS25 | -1.1004264 | 0.0006465 |
| TBCK | -1.0997912 | 0.0063032 |
| NTNG1 | -1.0955142 | 0.0393997 |
| PTAFR | -1.0944356 | 0.0247293 |
| SLC40A1 | -1.0896357 | 0.0275917 |
| USP17L12 | -1.0872304 | 0.0341098 |
| ALDH7A1 | -1.0856478 | 0.0218826 |
| PLCE1 | -1.0570644 | 0.0018764 |
| GLIPR1 | -1.0567817 | 0.0265507 |
| LYZ | -1.0562064 | 0.0103344 |
| TYROBP | -1.054305 | 0.0290836 |
| COPZ2 | -1.0527563 | 0.020402 |
| SERPINF1 | -1.0520324 | 0.0186876 |
| OLFML2B | -1.0465453 | 0.0050836 |
| EREG | -1.0391909 | 0.005154 |
| ENG | -1.0367254 | 0.0134078 |
| PGM3 | -1.0364344 | 0.0412552 |
| HLA-DMA | -1.0357603 | 0.0485598 |
| PLXDC2 | -1.032759 | 0.0384433 |
| GYS2 | -1.0258701 | 0.0260651 |
| FCGR2C | -1.0247323 | 0.0481219 |
| DDIT4 | -1.0209938 | 0.0339844 |
| FGF11 | -1.0198215 | 0.0387436 |
| C1orf174 | -1.0162379 | 0.0403707 |
| TUBA3E | -1.0116719 | 0.0266323 |
| GTPBP6 | -1.0116061 | 0.0116776 |
| FOLR2 | -1.0111569 | 0.0487284 |
| PLSCR4 | -1.0098222 | 0.0412686 |
| ADGRB2 | -1.005412 | 0.0084852 |
| LCP1 | -0.9919122 | 0.0120309 |
| C2orf49 | -0.9866714 | 0.0463594 |
| GIMAP4 | -0.9860181 | 0.0071358 |
| PFKM | -0.9766652 | 0.0267131 |
| CXCR4 | -0.9698121 | 0.0177769 |
| RNF144B | -0.9678879 | 0.0083128 |
| TANK | -0.9653491 | 0.0330324 |
| KCTD12 | -0.961717 | 0.0312338 |
| RGS6 | -0.9604287 | 0.0448931 |

| | | |
|---|---|---|
| DCXR | -0.9579577 | 0.0239685 |
| FRS2 | -0.956011 | 0.03201 |
| TMEM51 | -0.9505797 | 0.0191124 |
| SLC11A1 | -0.9502457 | 0.0371178 |
| OFD1 | -0.9466491 | 0.0099233 |
| FNDC4 | -0.9427252 | 0.0033662 |
| VPS54 | -0.9413233 | 0.0329302 |
| ZBED6 | -0.927861 | 0.0322218 |
| TRIM22 | -0.9250411 | 0.0492793 |
| UNC93B1 | -0.9227609 | 0.047718 |
| MOSPD2 | -0.9218228 | 0.0139712 |
| C5AR1 | -0.9212206 | 0.043716 |
| DHRS3 | -0.9162057 | 0.0185025 |
| MPEG1 | -0.9151831 | 0.0039451 |
| ARPC1B | -0.9114509 | 0.0409373 |
| TPSAB1 | -0.9086543 | 0.0129766 |
| HDAC1 | -0.9086361 | 0.0041484 |
| MAF | -0.9033879 | 0.0309243 |
| EPHX1 | -0.9003594 | 0.0220962 |
| THAP5 | -0.8982176 | 0.0388258 |
| PABIR2 | -0.8970926 | 0.0199508 |
| CSF3R | -0.895848 | 0.0329154 |
| YTHDF1 | -0.8956085 | 0.0390216 |
| TDRD15 | -0.8942862 | 0.0435265 |
| HES7 | -0.8877452 | 0.0260972 |
| GPN3 | -0.8830135 | 0.0467059 |
| CYP1B1 | -0.8786665 | 0.0373084 |
| UROD | -0.8758141 | 0.0113403 |
| LMAN2L | -0.8751159 | 0.0144966 |
| NR4A3 | -0.8725561 | 0.0297596 |
| MOB3A | -0.8643671 | 0.004946 |
| ATG2B | -0.8567521 | 0.0106373 |
| PIAS1 | -0.8553154 | 0.0026118 |
| RILPL2 | -0.85369 | 0.0291166 |
| LGALS7B | -0.8514039 | 0.0095436 |
| DGLUCY | -0.8507855 | 0.0371445 |
| MRPS7 | -0.8507613 | 0.0400137 |
| HS2ST1 | -0.8447376 | 0.0192516 |
| SUCLG2 | -0.8410758 | 0.0271037 |
| ZNF416 | -0.8401843 | 0.0207483 |
| LGALS3 | -0.8395656 | 0.0434159 |
| CNN2 | -0.8376603 | 0.0176908 |
| ATG16L2 | -0.8370516 | 0.0114051 |
| UBE2A | -0.8346102 | 0.0057661 |
| SERTAD2 | -0.834415 | 0.0164604 |
| COMMD6 | -0.8342679 | 0.0479491 |
| NBPF14 | -0.8334351 | 0.0240918 |
| PPP1R15A | -0.8330758 | 0.0456251 |

| | | |
|---|---|---|
| ARFGEF1 | -0.8317795 | 0.0036011 |
| PECAM1 | -0.83068 | 0.0293162 |
| UBE2D4 | -0.8247319 | 0.0047794 |
| RNF149 | -0.8146499 | 0.01783 |
| UTP6 | -0.8132411 | 0.0012957 |
| RIPOR2 | -0.8060474 | 0.0110987 |
| THEMIS2 | -0.8050003 | 0.0362687 |
| STXBP2 | -0.8028684 | 0.0136072 |
| DPY19L2 | -0.8022127 | 0.0335818 |
| TBXAS1 | -0.8009426 | 0.0268149 |
| MIA2 | -0.799426 | 0.0170445 |
| C2CD5 | -0.7987686 | 0.0088873 |
| APH1A | -0.7966485 | 0.0435447 |
| TMEM97 | -0.7940659 | 0.0209932 |
| DOCK7 | -0.7940195 | 0.009415 |
| FHIP2B | -0.7909547 | 0.0163488 |
| WNT6 | -0.7905705 | 0.0313979 |
| TAS2R7 | -0.7900469 | 0.0194699 |
| NUBP2 | -0.7880797 | 0.0103516 |
| KRTAP12-3 | -0.7868762 | 0.0161394 |
| INPPL1 | -0.7849297 | 0.0170156 |
| TGM4 | -0.7839388 | 0.043726 |
| KANSL3 | -0.7813743 | 0.0049374 |
| GGTA1 | -0.7795062 | 0.0469101 |
| RBM23 | -0.7786957 | 0.0025352 |
| CYLD | -0.7783984 | 0.0495366 |
| PDE4C | -0.7774407 | 0.0264762 |
| FXYD5 | -0.7757651 | 0.0444263 |
| SCIN | -0.7730473 | 0.0388336 |
| CPQ | -0.7667247 | 0.0461294 |
| IZUMO4 | -0.766696 | 0.0212225 |
| DUSP10 | -0.7650108 | 0.0155482 |
| IGFBP6 | -0.7627749 | 0.0385405 |
| PSMD12 | -0.7616355 | 0.0335715 |
| SHPRH | -0.7574803 | 0.0059966 |
| CERK | -0.7574607 | 0.0294819 |
| SLC25A39 | -0.7573615 | 0.0317971 |
| LAMB2 | -0.7565821 | 0.0354412 |
| SNX7 | -0.756311 | 0.0335073 |
| EIF4ENIF1 | -0.7534294 | 0.0178381 |
| MARVELD1 | -0.751587 | 0.033219 |
| MYO1G | -0.750955 | 0.0382997 |
| JMY | -0.7506583 | 0.01132 |
| BHLHE40 | -0.7482283 | 0.0471159 |
| GTPBP2 | -0.7460006 | 0.0083603 |
| ANXA11 | -0.7440657 | 0.0208784 |
| YIPF5 | -0.7422363 | 0.0152179 |
| NAXD | -0.7414707 | 0.0213545 |

| | | |
|---|---|---|
| TENT5C | -0.7407603 | 0.0465258 |
| LRRC59 | -0.7393478 | 0.016681 |
| TGFBR2 | -0.7362827 | 0.0213853 |
| LIME1 | -0.7356848 | 0.0129939 |
| PIK3AP1 | -0.7332592 | 0.0298641 |
| TMEM104 | -0.7327938 | 0.0131913 |
| RNF213 | -0.7310913 | 0.0450875 |
| DNAAF5 | -0.7305884 | 0.0260422 |
| ALDH9A1 | -0.7293247 | 0.0436827 |
| FPR1 | -0.7278347 | 0.0370206 |
| TNS1 | -0.7272667 | 0.0425289 |
| PIK3IP1 | -0.7252706 | 0.0389173 |
| BCHE | -0.7226054 | 0.027079 |
| PPP1R10 | -0.7215447 | 0.0135873 |
| EVA1B | -0.7194986 | 0.0432708 |
| RAD51D | -0.7142335 | 0.0182198 |
| TMEM19 | -0.714223 | 0.0217238 |
| GALNT18 | -0.712 | 0.0499051 |
| ZNF212 | -0.7113585 | 0.0251782 |
| ZNF280C | -0.708115 | 0.0311631 |
| PHKB | -0.7073304 | 0.0349316 |
| PPM1F | -0.7061659 | 0.0291858 |
| SP140L | -0.7034938 | 0.0282535 |
| GRIK4 | -0.7011855 | 0.0460608 |
| HEATR3 | -0.7005723 | 0.0240465 |
| CYHR1 | -0.6999764 | 0.0422528 |
| MEF2A | -0.6993339 | 0.0084335 |
| ATF6 | -0.6990897 | 0.0313694 |
| GASK1A | -0.6965811 | 0.0404782 |
| ATP13A1 | -0.6965546 | 0.033784 |
| XXYLT1 | -0.6949171 | 0.0226716 |
| NHSL1 | -0.6945069 | 0.0467429 |
| TAF1 | -0.6942735 | 0.0444842 |
| RNF166 | -0.691207 | 0.0479978 |
| PAXX | -0.6835455 | 0.0317728 |
| DNAJC16 | -0.6782817 | 0.0347837 |
| CPLANE1 | -0.6775843 | 0.0123012 |
| SULF1 | -0.6716753 | 0.0339205 |
| ADH5 | -0.6623025 | 0.0154995 |
| TGFBR3 | -0.661364 | 0.0292847 |
| NCOA4 | -0.6611935 | 0.0369762 |
| MPLKIP | -0.6600587 | 0.0261334 |
| GFER | -0.6553265 | 0.005877 |
| UBE2L6 | -0.655041 | 0.0353623 |
| ADPGK | -0.6536154 | 0.0361759 |
| MINDY1 | -0.6514812 | 0.0367316 |
| PIEZO1 | -0.6496091 | 0.0278267 |
| ETV6 | -0.6446932 | 0.0125526 |

| | | |
|---|---|---|
| ELMO2 | -0.6443836 | 0.0315728 |
| IL13 | -0.6415907 | 0.045898 |
| C2orf42 | -0.6386268 | 0.0343297 |
| PRAC1 | -0.6354866 | 0.0445429 |
| MLANA | -0.6337427 | 0.0459249 |
| RETREG3 | -0.6333629 | 0.0279548 |
| MAN2B2 | -0.6328913 | 0.0425828 |
| EEF1E1 | -0.6297207 | 0.0378957 |
| SLC39A7 | -0.6273383 | 0.0283162 |
| MRPS11 | -0.6262439 | 0.0335342 |
| YY1 | -0.619608 | 0.0427011 |
| ADCY6 | -0.6090405 | 0.0314991 |
| N4BP1 | -0.604749 | 0.0191353 |
| CNOT8 | -0.5997484 | 0.0112474 |
| TSNAX | -0.5924658 | 0.0191581 |
| SIAH2 | -0.58684 | 0.0291129 |
| ZBTB22 | -0.5827379 | 0.0401276 |
| RAMAC | -0.5816169 | 0.0192341 |
| DRG2 | -0.5770698 | 0.0238234 |
| TMEM11 | -0.5734654 | 0.03229 |
| IFFO2 | -0.5724996 | 0.0427167 |
| RREB1 | -0.5450433 | 0.0329386 |
| KCTD5 | -0.5421623 | 0.0486235 |
| RIPK1 | -0.5273514 | 0.0482197 |
| BET1L | -0.5198666 | 0.0403169 |
| CREBBP | -0.5071834 | 0.0463599 |
| DHRS4 | -0.497386 | 0.0269833 |

| ID | logFC | P.Value |
|---|---|---|
| TRIM49D2 | 0.5242036 | 0.048983 |
| DMXL2 | 0.5286786 | 0.0413068 |
| SCAMP5 | 0.530414 | 0.0418049 |
| MAPRE3 | 0.5378341 | 0.038262 |
| ROGDI | 0.5378684 | 0.0267771 |
| NDUFA8 | 0.5435791 | 0.0463249 |
| KLHDC2 | 0.5501447 | 0.0242936 |
| CILK1 | 0.5545269 | 0.0366216 |
| COG2 | 0.5548117 | 0.0329044 |
| RTL8C | 0.5722564 | 0.0431847 |
| TNFRSF25 | 0.5783318 | 0.0492614 |
| TRIP11 | 0.5793929 | 0.0365935 |
| GNAO1 | 0.5802055 | 0.0463937 |
| PSMD1 | 0.5842476 | 0.0217297 |
| CCDC12 | 0.5873775 | 0.0385084 |
| KHDC4 | 0.5911948 | 0.0448856 |
| C8orf58 | 0.5931157 | 0.0132679 |
| MRTFA | 0.5968671 | 0.039644 |
| SEC16A | 0.5977039 | 0.0389501 |
| GATB | 0.5978006 | 0.0197561 |
| HTT | 0.5997346 | 0.0363817 |
| APTX | 0.6006768 | 0.0362765 |
| EVI5 | 0.6033261 | 0.0357065 |
| MARK3 | 0.6037453 | 0.0455729 |
| ZNRD2 | 0.6065226 | 0.0179895 |
| TENT2 | 0.6093861 | 0.0377352 |
| FASN | 0.6101746 | 0.0375968 |
| DTX1 | 0.6111445 | 0.0376281 |
| ZNF594 | 0.6111563 | 0.0306333 |
| DGKH | 0.6129631 | 0.0289139 |
| SERF1A | 0.6137708 | 0.0433047 |
| TG | 0.6147609 | 0.0365589 |
| LDB1 | 0.616809 | 0.0289732 |
| RBIS | 0.617486 | 0.0496503 |
| SDCCAG8 | 0.6192256 | 0.0420363 |
| ING5 | 0.619535 | 0.0303442 |
| C20orf96 | 0.6201043 | 0.020235 |
| OSBP2 | 0.6208463 | 0.0460883 |
| SECISBP2L | 0.6209028 | 0.044992 |
| AGO1 | 0.6225817 | 0.0387006 |
| LSM1 | 0.6244452 | 0.0395668 |
| RNF220 | 0.6247978 | 0.0281345 |
| SLC9B2 | 0.6344419 | 0.0207638 |
| LTV1 | 0.6348205 | 0.0329617 |
| POLR1E | 0.6385867 | 0.0352172 |
| SETD4 | 0.6400439 | 0.0220228 |
| RNF144A | 0.6411006 | 0.025491 |

| | | |
|---|---|---|
| MACROH2A | 0.641251 | 0.0397796 |
| CAMK1D | 0.645494 | 0.0392021 |
| MAVS | 0.6477177 | 0.0399828 |
| GABPA | 0.6477975 | 0.0093133 |
| SNU13 | 0.6493249 | 0.0257158 |
| ZNF577 | 0.6545162 | 0.0390718 |
| ABI2 | 0.6563741 | 0.0499226 |
| SORCS2 | 0.6576281 | 0.0408113 |
| TAGLN3 | 0.6580659 | 0.0300329 |
| MCUR1 | 0.6590909 | 0.044311 |
| HIVEP2 | 0.6595057 | 0.0263027 |
| ZSCAN26 | 0.6602937 | 0.0357555 |
| ELAC2 | 0.6624177 | 0.0343815 |
| SPG7 | 0.6632117 | 0.0201954 |
| GID8 | 0.6632705 | 0.0144898 |
| FBXL2 | 0.6640187 | 0.0273287 |
| PRKCB | 0.6660126 | 0.046104 |
| MITD1 | 0.6664879 | 0.021507 |
| EPB41L1 | 0.6695999 | 0.0464934 |
| ALG5 | 0.6703285 | 0.04876 |
| ABHD17A | 0.6718333 | 0.0251504 |
| SEC22B | 0.6742429 | 0.0249804 |
| IGSF9B | 0.6744025 | 0.0470921 |
| NUP35 | 0.6779122 | 0.0404867 |
| FUNDC1 | 0.6810352 | 0.0167241 |
| AMY2B | 0.6817521 | 0.0129649 |
| NMNAT2 | 0.6858087 | 0.0491385 |
| SEC22A | 0.6865439 | 0.0279385 |
| RAB5B | 0.6868857 | 0.0481022 |
| SLC1A4 | 0.6870502 | 0.0316266 |
| OPTN | 0.6883281 | 0.006124 |
| DBF4B | 0.6906121 | 0.0229431 |
| RAD51B | 0.6906672 | 0.0252558 |
| NKX2-2 | 0.6913927 | 0.0465074 |
| GRSF1 | 0.6945256 | 0.0338407 |
| ATL1 | 0.6950908 | 0.0222353 |
| MAP3K7CL | 0.6958281 | 0.0298216 |
| KLC2 | 0.6980578 | 0.0304553 |
| SHC4 | 0.6986806 | 0.0454459 |
| RHOT2 | 0.6994927 | 0.0478014 |
| ALYREF | 0.6998895 | 0.0284059 |
| ATF2 | 0.7016622 | 0.0347011 |
| CLK1 | 0.7022997 | 0.030599 |
| GLRA3 | 0.7031516 | 0.0329048 |
| RASA4 | 0.704064 | 0.0347701 |
| TOP2A | 0.7066797 | 0.0250782 |
| ZNF317 | 0.7104406 | 0.0312949 |
| TIMM21 | 0.7105996 | 0.0285684 |

| | | |
|---|---|---|
| GTPBP1 | 0.7109764 | 0.0192068 |
| ZDHHC21 | 0.7110954 | 0.0144816 |
| PTK2 | 0.7117538 | 0.0126953 |
| ANKRD11 | 0.7118799 | 0.0314852 |
| NPRL3 | 0.7144975 | 0.0096638 |
| IMP4 | 0.7158521 | 0.0153508 |
| CEP170 | 0.7198583 | 0.0464536 |
| SIAH1 | 0.7205076 | 0.0268163 |
| CNOT9 | 0.7213654 | 0.0270255 |
| RUFY2 | 0.7243673 | 0.006168 |
| PAIP2 | 0.725646 | 0.0347464 |
| SLIT3 | 0.7257633 | 0.0234302 |
| INTS12 | 0.7288022 | 0.0413029 |
| SRRM3 | 0.7289583 | 0.023142 |
| STK25 | 0.7301944 | 0.0184763 |
| CXXC1 | 0.7315211 | 0.0105114 |
| LAS1L | 0.7339987 | 0.0194149 |
| PRR4 | 0.7341688 | 0.0497588 |
| HARS1 | 0.7356386 | 0.0183037 |
| PIK3CB | 0.7370492 | 0.0317024 |
| REPIN1 | 0.7390355 | 0.0385701 |
| NPLOC4 | 0.740629 | 0.0398081 |
| TOMM20 | 0.7465128 | 0.0439788 |
| LANCL1 | 0.7474174 | 0.0310926 |
| ABCB10 | 0.7516521 | 0.0126879 |
| PDXP | 0.7524308 | 0.025398 |
| GOLGA4 | 0.7524505 | 0.0197087 |
| TYRO3 | 0.7536619 | 0.027023 |
| STRN3 | 0.7573245 | 0.0152335 |
| GNAI1 | 0.7585698 | 0.033841 |
| GCDH | 0.7588369 | 0.0293468 |
| ZMYM2 | 0.7590571 | 0.0220948 |
| PGBD5 | 0.764424 | 0.0178677 |
| PLPPR1 | 0.7691254 | 0.0279654 |
| PDP1 | 0.7726059 | 0.0346903 |
| GAS2L3 | 0.7733424 | 0.0201626 |
| DYNLL1 | 0.7750605 | 0.0168813 |
| WSB2 | 0.7768364 | 0.0462504 |
| PSIP1 | 0.7788282 | 0.0156036 |
| RPS19BP1 | 0.7796275 | 0.0206505 |
| ARHGDIG | 0.7848269 | 0.0469046 |
| MTUS1 | 0.7862342 | 0.0431076 |
| PSMB7 | 0.7897888 | 0.0117678 |
| SLITRK2 | 0.7898568 | 0.0364954 |
| PAK1 | 0.7937898 | 0.0096331 |
| MAP1A | 0.7950669 | 0.0168209 |
| LMBR1 | 0.7970021 | 0.0403881 |
| ATP2B2 | 0.79906 | 0.0189019 |

| | | |
|---|---|---|
| TCEA2 | 0.7993198 | 0.0088265 |
| CNKSR2 | 0.8015202 | 0.0267716 |
| KIF5B | 0.8059566 | 0.0157698 |
| CALY | 0.8070696 | 0.0482355 |
| ILF3 | 0.8130013 | 0.0306071 |
| PRR18 | 0.8146763 | 0.0110357 |
| KRAS | 0.817112 | 0.0110335 |
| TPD52 | 0.8193845 | 0.041259 |
| ZFP3 | 0.8200116 | 0.015552 |
| NRBP2 | 0.8200999 | 0.0332364 |
| PODXL2 | 0.8205071 | 0.0323581 |
| KCTD17 | 0.8230463 | 0.0305993 |
| CERS4 | 0.8236274 | 0.0110604 |
| TENT4A | 0.8236641 | 0.0341965 |
| TTLL7 | 0.8250863 | 0.0119185 |
| SLAIN1 | 0.8263885 | 0.0328536 |
| NEUROD6 | 0.827865 | 0.0113967 |
| RTL5 | 0.8281241 | 0.0095881 |
| SLC12A2 | 0.8316699 | 0.0175096 |
| ZNF841 | 0.8320019 | 0.0370193 |
| MAPK8IP3 | 0.833608 | 0.0086628 |
| TTLL11 | 0.8374154 | 0.0190167 |
| CTTN | 0.8443942 | 0.0178468 |
| ANKS1B | 0.8457789 | 0.0374604 |
| LY6H | 0.8492956 | 0.0250077 |
| RABGEF1 | 0.8504407 | 0.0028088 |
| IL1B | 0.8516976 | 0.0157779 |
| MAPK10 | 0.854329 | 0.0169599 |
| FN3K | 0.8581629 | 0.021214 |
| HIPK2 | 0.8646233 | 0.044526 |
| MAP1LC3A | 0.866821 | 0.0075188 |
| TRIT1 | 0.8689973 | 0.0043246 |
| EGFL8 | 0.8696428 | 0.0161693 |
| GET4 | 0.8700727 | 0.0204412 |
| DOT1L | 0.8702781 | 0.0251942 |
| TMEM59L | 0.8731976 | 0.0164481 |
| NPDC1 | 0.8736893 | 0.0404317 |
| C3orf70 | 0.8756821 | 0.0299764 |
| SRCIN1 | 0.8818734 | 0.019416 |
| ACTR3B | 0.8828849 | 0.005631 |
| NREP | 0.8844859 | 0.0362375 |
| JPH4 | 0.8858933 | 0.0200921 |
| MAPRE2 | 0.8882403 | 0.0041862 |
| C1orf198 | 0.8885969 | 0.0215193 |
| SLITRK3 | 0.8933798 | 0.0033809 |
| CWC22 | 0.8936363 | 0.0039102 |
| CEBPG | 0.8954889 | 0.0034089 |
| RRP7A | 0.8959528 | 0.0147359 |

| | | |
|---|---|---|
| GPD1 | 0.9011628 | 0.0037565 |
| ADIPOR2 | 0.9011921 | 0.0303521 |
| SETD2 | 0.9028423 | 0.0047793 |
| MINDY3 | 0.9047778 | 0.0066017 |
| HID1 | 0.9059754 | 0.0034747 |
| PLG | 0.9062949 | 0.009031 |
| PEG3 | 0.9138353 | 0.0248815 |
| NEFM | 0.9144504 | 0.0281789 |
| LGI3 | 0.9163613 | 0.029665 |
| ADGRV1 | 0.9174747 | 0.0489761 |
| TRIM24 | 0.9269843 | 0.0468977 |
| SNRPN | 0.9279747 | 0.0265262 |
| CITED1 | 0.9286814 | 0.0417064 |
| MRPS35 | 0.9332526 | 0.0020294 |
| VAMP2 | 0.9369313 | 0.0394722 |
| PNMA2 | 0.9430948 | 0.0260662 |
| EPN2 | 0.9501121 | 0.0131249 |
| NAPB | 0.9510364 | 0.0413856 |
| DHCR24 | 0.9538157 | 0.0120989 |
| CEND1 | 0.9539019 | 0.0419378 |
| TMEFF2 | 0.9539536 | 0.0269316 |
| SYTL2 | 0.9563018 | 0.0400613 |
| LBHD1 | 0.9582456 | 0.0042015 |
| ADAM23 | 0.9602243 | 0.0053935 |
| PKD1 | 0.968076 | 0.0278365 |
| SLC22A17 | 0.9750245 | 0.0220391 |
| RNF11 | 0.9782289 | 0.0071877 |
| DDX59 | 0.9812633 | 0.0079886 |
| G3BP2 | 0.9820182 | 0.0204545 |
| KCNMB4 | 0.9828555 | 0.0408817 |
| SEL1L3 | 0.9872477 | 0.0454384 |
| FEZ1 | 0.9953849 | 0.0313559 |
| BRSK1 | 0.9974972 | 0.0017572 |
| TMOD2 | 1.0009757 | 0.0258472 |
| PALM | 1.0023206 | 0.0065192 |
| PHLDB1 | 1.0135623 | 0.0048206 |
| EFCAB2 | 1.0151879 | 0.0011607 |
| SCG5 | 1.0168615 | 0.0314576 |
| CNTN1 | 1.0192546 | 0.0127878 |
| AGAP1 | 1.0305148 | 0.0050237 |
| YWHAG | 1.031574 | 0.0175293 |
| SNURF | 1.0443153 | 0.0126647 |
| INPP5F | 1.0455595 | 0.0058151 |
| IGSF8 | 1.0469318 | 0.0014093 |
| ZNF428 | 1.0481263 | 0.0103258 |
| WASHC5 | 1.0486006 | 0.003607 |
| ATP8A1 | 1.0501813 | 0.0077788 |
| ATRNL1 | 1.0629626 | 0.0146083 |

| | | |
|---|---|---|
| HPCAL4 | 1.0635363 | 0.0102603 |
| STX1A | 1.0717961 | 0.0020094 |
| CPSF6 | 1.0721908 | 0.0022755 |
| SYT11 | 1.0730472 | 0.0097585 |
| GRIA2 | 1.0786939 | 0.0070205 |
| YWHAH | 1.0818069 | 0.0095001 |
| KCNQ2 | 1.0881236 | 0.0077214 |
| TCEAL7 | 1.0885335 | 0.0008364 |
| STMN4 | 1.0923544 | 0.0103521 |
| BSCL2 | 1.1030608 | 0.0009156 |
| FAIM2 | 1.1204175 | 0.0051222 |
| ALDH1A3 | 1.1251115 | 0.0281471 |
| VSNL1 | 1.1276159 | 0.028293 |
| C1orf21 | 1.1306347 | 0.0027318 |
| RNF157 | 1.1316546 | 0.014161 |
| EGR3 | 1.1385488 | 0.0264076 |
| LARP6 | 1.1442856 | 0.0062558 |
| SCN2A | 1.1453161 | 0.0024323 |
| KXD1 | 1.1467662 | 0.0005697 |
| CALM3 | 1.1488655 | 0.0248909 |
| NGFR | 1.1533802 | 0.0388333 |
| CNTN2 | 1.1537011 | 0.0232773 |
| EXOC1 | 1.1591219 | 0.0024591 |
| H3C7 | 1.16629 | 0.0457467 |
| ARC | 1.1692467 | 0.0486351 |
| TUBB2A | 1.1760343 | 0.0340314 |
| SNCG | 1.1846093 | 0.0478032 |
| LZTS3 | 1.1864623 | 0.0100683 |
| NRXN1 | 1.2014037 | 0.0032032 |
| CALM1 | 1.2234888 | 0.0228978 |
| DIRAS1 | 1.2395755 | 0.0014 |
| RAB3A | 1.2452831 | 0.0137669 |
| CAMK2N1 | 1.2490452 | 0.0470045 |
| KIF1A | 1.2514605 | 0.0075364 |
| QDPR | 1.2568246 | 0.0018352 |
| CLIP3 | 1.2761187 | 0.0063247 |
| RBFOX1 | 1.2811094 | 0.0070522 |
| DNAJC10 | 1.3079445 | 0.0407053 |
| KIF5A | 1.3523114 | 0.0348401 |
| MAG | 1.3553616 | 0.0357851 |
| SNAP25 | 1.4309929 | 0.0462805 |
| GP1BB | 1.4914397 | 0.0001968 |
| SEPTIN5 | 1.5165537 | 0.0087584 |
| CAMK1 | 1.5169066 | 0.0031358 |
| S100B | 1.5369072 | 0.0435468 |
| APLP1 | 1.5617067 | 0.0184266 |
| KIF5C | 1.5779305 | 0.0039099 |
| SYP | 1.5925466 | 0.0119637 |

| | | |
|---|---|---|
| PRMT3 | 1.6517491 | 0.0000977 |
| MAP2 | 1.7594252 | 0.0046353 |
| STMN1 | 1.820418 | 0.0048325 |
| OLFM1 | 1.8338646 | 0.0464422 |
| NRGN | 1.9190336 | 0.008786 |
| PCSK1N | 1.9214053 | 0.0383515 |

Table S8

| ID | logFC | P.Value |
|---|---|---|
| HSPA1B | -1.1770605 | 0.0033788 |
| HSPA1A | -1.0795489 | 0.0023284 |
| CNR1 | -0.8159163 | 0.0446435 |
| RAB31 | -0.7887974 | 0.0190535 |
| CCT8 | -0.6281744 | 0.0027655 |
| S1PR3 | -0.6160168 | 0.0035066 |
| TNFRSF21 | -0.5903934 | 0.0212191 |
| TPM1 | -0.5707825 | 0.0057537 |
| KCNJ16 | -0.5688971 | 0.0133253 |
| CDKN2B | -0.5582175 | 0.0160868 |
| CAV2 | -0.5486257 | 0.018535 |
| DCN | -0.5181897 | 0.0158091 |
| DHCR24 | -0.5139064 | 0.0168441 |
| DSTN | -0.5119612 | 0.0410817 |
| LIX1 | -0.5077133 | 0.0031899 |
| ARL6IP6 | -0.5067517 | 0.0285103 |
| COL4A1 | -0.5017285 | 0.0477487 |
| TPRKB | -0.4952938 | 0.0181186 |
| WIPF2 | -0.4927143 | 0.0424567 |
| C21orf62 | -0.4881124 | 0.0274935 |
| DCAF12 | -0.4828536 | 0.0348104 |
| LRRN1 | -0.4809925 | 0.0038355 |
| PAICS | -0.4776644 | 0.049203 |
| YTHDF3 | -0.4754159 | 0.0054674 |
| APOLD1 | -0.4717068 | 0.0242592 |
| RBMXL1 | -0.4640337 | 0.0182586 |
| MIEN1 | -0.4616738 | 0.0044235 |
| SCX | -0.4564102 | 0.008105 |
| SH3RF3 | -0.4553902 | 0.0241615 |
| MEMO1 | -0.4504666 | 0.0218457 |
| UBE2A | -0.4489075 | 0.0057499 |
| CCDC80 | -0.4483892 | 0.0341261 |
| ERG28 | -0.4452695 | 0.0011624 |
| GGH | -0.4413511 | 0.0281892 |
| ZEB1 | -0.4410447 | 0.0337107 |
| SSR3 | -0.4390706 | 0.0488627 |
| MRPL2 | -0.43608 | 0.0179588 |
| ZMAT3 | -0.4354652 | 0.0160317 |
| SPRY2 | -0.4207749 | 0.0486065 |
| ACVR1 | -0.4179451 | 0.0033595 |
| CCDC124 | -0.4171997 | 0.0226609 |
| COL3A1 | -0.4147777 | 0.0301962 |
| LRRC59 | -0.414389 | 0.0156936 |
| SCG3 | -0.4117702 | 0.039339 |
| RAB5IF | -0.4116814 | 0.0218281 |
| AKT1 | -0.4043014 | 0.0179026 |

| | | |
|---|---|---|
| NAXE | -0.4035576 | 0.0403111 |
| ETFDH | -0.4010158 | 0.0199089 |
| NDUFB9 | -0.3922037 | 0.03022 |
| DNM1L | -0.3915344 | 0.0150656 |
| STRBP | -0.3885971 | 0.0276138 |
| ECE1 | -0.3873116 | 0.0402022 |
| DNAJA1 | -0.3842476 | 0.0432633 |
| TCEAL8 | -0.3838105 | 0.0150475 |
| DERA | -0.3833367 | 0.041322 |
| OR10AD1 | -0.382545 | 0.0431657 |
| MCTS1 | -0.381452 | 0.0070973 |
| TSPAN13 | -0.3811393 | 0.0272492 |
| AFG3L2 | -0.380462 | 0.0227156 |
| TMPO | -0.3797068 | 0.0497572 |
| ALG14 | -0.3776877 | 0.0442935 |
| TIMM10 | -0.3772349 | 0.0162571 |
| ATP2B2 | -0.3748111 | 0.0434886 |
| TRA2B | -0.3694955 | 0.0345813 |
| RASSF4 | -0.3670981 | 0.0227592 |
| STX5 | -0.3663829 | 0.0431185 |
| RAB6A | -0.3636873 | 0.0391181 |
| EI24 | -0.3622858 | 0.0101172 |
| IFT52 | -0.359279 | 0.0456021 |
| NAP1L3 | -0.3585745 | 0.0201923 |
| POGLUT1 | -0.3585692 | 0.0481102 |
| MEF2C | -0.3541552 | 0.0337522 |
| WASL | -0.3521433 | 0.0455381 |
| ZBTB33 | -0.3520752 | 0.0161319 |
| TCEAL9 | -0.351092 | 0.0247186 |
| BROX | -0.3507257 | 0.0290375 |
| BPGM | -0.3485042 | 0.0221151 |
| DNAI1 | -0.3462277 | 0.0411508 |
| PPM1G | -0.3448795 | 0.0375719 |
| MRPS33 | -0.344755 | 0.0172933 |
| ICMT | -0.3443795 | 0.0493585 |
| JMJD6 | -0.3365024 | 0.039294 |
| ZNF195 | -0.3358738 | 0.0119512 |
| ATP6V0D1 | -0.3329368 | 0.0335694 |
| CUL4B | -0.3273248 | 0.0358527 |
| ACSL4 | -0.3258645 | 0.028818 |
| APOO | -0.325862 | 0.0384187 |
| SMURF2 | -0.3253889 | 0.0188157 |
| INTS10 | -0.3246999 | 0.0318955 |
| CCDC14 | -0.3210361 | 0.0315943 |
| GOSR1 | -0.3192576 | 0.0427459 |
| CRIP2 | -0.3188298 | 0.0261698 |
| SOHLH1 | -0.318419 | 0.046719 |
| HDAC3 | -0.3183372 | 0.0117426 |

| | | |
|---|---|---|
| SNRPB2 | -0.3180244 | 0.049995 |
| GSTA4 | -0.3123298 | 0.0436639 |
| TSPAN6 | -0.3101634 | 0.0321489 |
| DESI1 | -0.3053045 | 0.0276643 |
| SMG7 | -0.3050284 | 0.0334413 |
| PNP | -0.3040894 | 0.0302306 |
| RBBP8 | -0.3003465 | 0.045583 |
| CHCHD5 | -0.2977098 | 0.0263434 |
| POGLUT3 | -0.2976725 | 0.030689 |
| PRDX3 | -0.2936703 | 0.0398789 |
| CHD3 | -0.29337 | 0.0385965 |
| PYGO2 | -0.2922746 | 0.0384237 |
| ENTPD6 | -0.2895131 | 0.0484216 |
| SCAMP5 | -0.2888543 | 0.0309383 |
| CKAP2 | -0.2875637 | 0.0468237 |
| HCCS | -0.2855259 | 0.0257819 |
| SLC41A1 | -0.2851292 | 0.0218041 |
| C1orf216 | -0.2846545 | 0.0242562 |
| UBE2R2 | -0.2838997 | 0.0459922 |
| EIF2S2 | -0.2819105 | 0.0464597 |
| ASAP1 | -0.2713219 | 0.0337838 |
| SREK1 | -0.2701779 | 0.0421032 |
| OXR1 | -0.2647283 | 0.0374153 |
| DR1 | -0.2616101 | 0.0385892 |
| PFDN4 | -0.2491911 | 0.0490218 |
| ELMO2 | -0.2465914 | 0.0419222 |
| SETD4 | -0.240314 | 0.0368224 |
| RPF2 | -0.2247194 | 0.0473559 |

Table S9

| ID | logFC | P.Value |
|---|---|---|
| MT3 | -2.1421516 | 0.0390925 |
| PLA2G2A | -1.666964 | 0.0145296 |
| SOX13 | -1.3901869 | 0.0194534 |
| LRRN2 | -1.3594998 | 0.0229775 |
| SLC13A3 | -1.2960257 | 0.0026692 |
| PSMB10 | -1.1817549 | 0.0189279 |
| TIMP4 | -1.1515896 | 0.0413052 |
| LHFPL3 | -1.074258 | 0.0297385 |
| CDO1 | -1.0589175 | 0.0150565 |
| GSTM1 | -1.0539138 | 0.0360231 |
| PCDH9 | -1.0202208 | 0.0490553 |
| OXTR | -0.9761209 | 0.0462061 |
| MT1F | -0.9454169 | 0.036464 |
| SCN7A | -0.9301521 | 0.0176824 |
| CPVL | -0.9192903 | 0.0116371 |
| SALL2 | -0.8955581 | 0.0298013 |
| LHFPL6 | -0.8819274 | 0.0361568 |
| CHPF2 | -0.8737764 | 0.0482079 |
| SNRPE | -0.8631275 | 0.0480092 |
| BANF1 | -0.8299606 | 0.0305625 |
| RASL10A | -0.8232662 | 0.0111404 |
| H2BU1 | -0.7964202 | 0.0195083 |
| PRR4 | -0.7909397 | 0.0355338 |
| TRIAP1 | -0.7906548 | 0.0203456 |
| FKBP9 | -0.7580489 | 0.0475727 |
| TPRKB | -0.7414154 | 0.0137872 |
| CFAP126 | -0.7382032 | 0.0377267 |
| ZNF85 | -0.7143841 | 0.048491 |
| CES3 | -0.70029 | 0.0161595 |
| TMEM167A | -0.6901024 | 0.0311815 |
| GPR137B | -0.6880465 | 0.0241592 |
| NTAN1 | -0.6854528 | 0.0441248 |
| LONRF2 | -0.6775738 | 0.045562 |
| ARHGEF6 | -0.6484924 | 0.0384931 |
| CHPT1 | -0.6382877 | 0.0378623 |
| TMEM127 | -0.6337225 | 0.0472188 |
| EFCAB2 | -0.6307732 | 0.0323613 |
| SERF1A | -0.6106473 | 0.0443024 |
| UMAD1 | -0.6098539 | 0.0164594 |
| RNF165 | -0.5946142 | 0.04946 |
| ST7 | -0.5836576 | 0.0441373 |

| ID | logFC | P.Value |
|---|---|---|
| TNRC6C | 0.4739628 | 0.037959 |
| ZNF292 | 0.4928466 | 0.0468701 |
| GIGYF1 | 0.5013454 | 0.0492093 |
| CCAR2 | 0.5521645 | 0.0374247 |
| ZC3H7B | 0.5566781 | 0.0498735 |
| MGA | 0.5622681 | 0.0447888 |
| PRRC1 | 0.6050351 | 0.0450925 |
| FNBP4 | 0.6068211 | 0.0235447 |
| MTMR10 | 0.6088259 | 0.0449231 |
| CIAPIN1 | 0.6106324 | 0.0238822 |
| DHRS13 | 0.6119795 | 0.0322429 |
| LDB1 | 0.6124621 | 0.0300263 |
| FBXO10 | 0.6354688 | 0.0330152 |
| AMD1 | 0.6373133 | 0.043775 |
| SLC35E3 | 0.6719942 | 0.0186545 |
| HSD17B4 | 0.6721253 | 0.025391 |
| ADGRL2 | 0.6874639 | 0.0489148 |
| TYRO3 | 0.691039 | 0.0411939 |
| UBR1 | 0.7079217 | 0.0282472 |
| TRH | 0.7102485 | 0.048704 |
| SECISBP2L | 0.7265894 | 0.0205694 |
| ANK3 | 0.7275353 | 0.0136832 |
| SPON2 | 0.7595807 | 0.0387246 |
| SERPINB1 | 0.7773809 | 0.0310664 |
| SLC12A2 | 0.8007084 | 0.0216942 |
| SEC14L5 | 0.8051085 | 0.0167621 |
| TP53BP1 | 0.8102527 | 0.0193492 |
| SETD9 | 0.8124451 | 0.0429878 |
| ANKS1B | 0.8174788 | 0.0437792 |
| QDPR | 0.8274325 | 0.0311425 |
| RNF130 | 0.833463 | 0.023403 |
| TGM2 | 0.8511526 | 0.0369642 |
| PDIA3 | 0.8685114 | 0.0297365 |
| ANKRD36C | 0.8778109 | 0.015123 |
| APOL2 | 0.8803492 | 0.0264469 |
| ABCA2 | 0.9888759 | 0.0184732 |
| MAP3K1 | 1.069982 | 0.0248654 |
| RARRES1 | 1.0815062 | 0.0203412 |
| PIP4K2A | 1.1280487 | 0.0420047 |
| PPP1R14A | 1.2417506 | 0.037527 |
| MOG | 1.2798443 | 0.0491946 |
| PADI2 | 1.2840005 | 0.0092517 |
| MAG | 1.2904431 | 0.0448388 |
| CERCAM | 1.2981156 | 0.0119356 |
| UGT8 | 1.362535 | 0.0238564 |
| TF | 1.3927989 | 0.0264311 |
| LMCD1 | 1.4255578 | 0.0070711 |

| | | |
|---|---|---|
| PRDM2 | 1.603932 | 0.0096114 |
| SOX10 | 1.6446727 | 0.021292 |
